# Mathematical genomics of SRY proteins and their associated HMG-Box domains across mammals

**DOI:** 10.64898/2026.10.18.759944

**Authors:** Sk. Sarif Hassan, Nabanita Mukherjee, Moumita Sil, Arunava Goswami, Guy W. Dayhoff, Andrey P. Anisimov, Vladimir N. Uversky

**Author notes:** Corresponding author Email addresses (Sk. Sarif Hassan). (Nabanita Mukherjee), (Moumita Sil), (Arunava Goswami), (Guy W. Dayhoff II), (Andrey P. Anisimov), (Vladimir N. Uversky).

## Abstract

Male sex determination in mammals depends on sex-determining region Y (SRY), a transcription factor whose high-mobility group box (HMG box) binds and bends DNA to activate SOX9. The divergent N- and C-terminal domains lack sequence homology and have been dismissed as passive linkers, yet alignment-based homology cannot reveal whether they retain a hidden compositional grammar. We applied quantitative genomics to 65 full-length mammalian SRY proteins and 38 HMG-box subsequences across 75 mammals, integrating amino acid and dipeptide compositions, Shannon entropy, dominant-residue profiles, hydropathy-based n-gram spectra, longest hydropathy stretches, and cumulative proximity. The HMG box is tightly constrained: basic residues dominated, cysteine was nearly absent, entropy clustered near 4.0, polar content averaged 58.44%, and 25 of 38 HMG boxes shared 57.97% polar residues, whereas full-length SRY varied more, with rodents showing high polar and glutamine-rich profiles and cetaceans low polar content. Proximity-based clustering was more taxonomically consistent at the HMG level, while terminal regions blurred domain-level signals. Across 65 SRY proteins and 38 HMG domains, intrinsic disorder was the dominant predicted state, and a polyQ-rich rodent SRY illustrated how low-complexity repeats can produce misleading ordered predictions, underscoring the need for consensus disorder analysis in rapidly evolving intrinsically disordered regions. Cumulative proximity analysis identified *Bos taurus* and *Bos grunniens* as the only pair proximally clustered at both the full-length SRY and SRY-HMG levels, establishing multivariate proximity as a stringent criterion of compositional similarity satisfied by a single mammalian pair. These results suggest that SRY and its HMG box each carry a distinct compositional grammar, quantifiable only through alignment-independent signatures, with compositional individuality being the rule rather than the exception. These quantitative signatures provide a framework for interpreting SRY variation, assessing clinical mutations, and understanding evolutionary pressures shaping sex determination. Denser taxon sampling, structural modeling, and experimental tests are needed to clarify whether HMG-box drift and terminal-region variation affect DNA binding, bending, or reproductive function.

## 1. Introduction

The process of male sex determination in mammals begins with the action of the sex determining region Y (SRY) gene that is responsible for triggering the differentiation of the bipotential embryonic gonad to become testis [1, 2]. The SRY protein mainly acts as a transcription factor that binds to certain enhancers and activates essential downstream targets such as SOX9 [3, 4, 5]. In terms of structure, the SRY protein has an HMG box that is surrounded by the highly variable N-terminal (NTD) and C-terminal (CTD) domains [2].

For many years, the paradigmatic model used in the study of mammalian sex determination has revolved around the HMG box. Such emphasis is mainly warranted by the importance of such biophysical functions in this domain; it allows for sequence-specific DNA binding, it causes a sharp topological bending in the chromatin structure and contains nuclear localization signals important for cellular transport [6]. The importance of this core region both clinically and evolutionarily is immense. The majority of known clinical mutations in the SRY gene, which vary from relatively minor missense changes to highly damaging frameshift mutations, occur in the HMG box, resulting in disorders of sex development (DSDs) conditions, such as XY gonadal dysgenesis and full sexual reversal [3, 7, 8]. Since the HMG box demonstrates very high evolutionary conservation in many different organisms, genomic analysis techniques have depended a lot on it for identifying the phylogeny of male sex determination.

In contrast, the flanking non-HMG domains on either side of the HMG region have traditionally been overlooked in structural sequence studies. As the NTD and CTD show extremely rapid evolutionary divergence and show no significant homology in primary sequences between close mammalian species, they were long presumed to act merely as passive structural linkers or vestigial extensions [2, 9]. On the other hand, contemporary functional genomics has triggered an immense paradigm shift indicating that these extremely different regions are in fact biologically essential [10, 11]. The latest molecular studies have shown that the NTD and CTD play an active role in ensuring proper binding conformations, forming base-specific interactions, and associating with important transport pathways such as importin-*β* and co-regulatory systems [2, 6, 7]. Consequently, isolating the HMG box provides an incomplete picture of SRY functionality; a comprehensive evolutionary and structural understanding requires investigating the entire sequence architecture.

To fill this critical gap, this study employs methods of mathematical genomics to determine the structural signature of both the conserved HMG-box domain and the highly variable non-HMG-box domains of the SRY gene product. Extending the use of computational genomics methods used to determine mutation hotspots in expansive genomic regions, this investigation leverages high-resolution mathematical modelling to quantify structural features throughout the whole sequence of SRY [12, 13]. Previous studies have been able to detect genomic regions; however, our new approach uses analysis of the full-length protein in tandem with its corresponding isolated HMG-box subsequences.

The SRY proteins for this study were sourced from a wide range of seventy-five individual mammalian species. Through the process of curation, several duplicate evolution instances were detected for certain taxa, resulting in a final collection consisting of sixty-five individual SRY sequences and thirty-eight unique HMG box subsequences. To understand the underlying structure behind such proteins, the current research applies an advanced mathematical genomics pipeline to uncover this hidden structure. This multidimensional framework bridges classical multiple sequence homology with high-resolution topological metrics, evaluating amino acid and dipeptide compositions, Shannon entropy, window-based dominant residue distributions, and homogeneous polystring frequencies. In order to characterize the exact physicochemical architecture, hydropathy-based n-gram mapping (encompassing 2-gram, 3-gram, and 4-gram spectra) was applied in addition to continuous polar and non-polar residue stretches. Furthermore, to link sequence topology with functional biophysics, this study maps clinically significant phenotype-associated mutations in human SRY, evaluates intrinsic protein disorder (IPD), and predicts liquid–liquid phase separation (LLPS) propensity and condensate-forming capacity using the FuzDrop algorithm [14, 15, 16]. Through the integration of these distance matrices into dendograms, a cumulative proximity analysis to map the phylogenetic trajectories of both the full-length protein and its isolated domains was performed. Ultimately, assessing the whole SRY sequence together with its subsequence HMG box provides a robust, highly sensitive framework designed to redefine the structural, evolutionary, and functional interplay within this master regulatory protein.

## 2. Data acquisition

In this study, sex-determining region Y (SRY) protein sequences (UniProt reviewed) were extracted from seventy-five organisms (Table 1). Among these, ten sequences were identified as duplicates and the duplicated SRY sequences correspond to the organisms listed in Table 2. Consequently, out of the seventy-five sequences analyzed, sixty-five were found to be unique (Table 3).

**Table 1:**
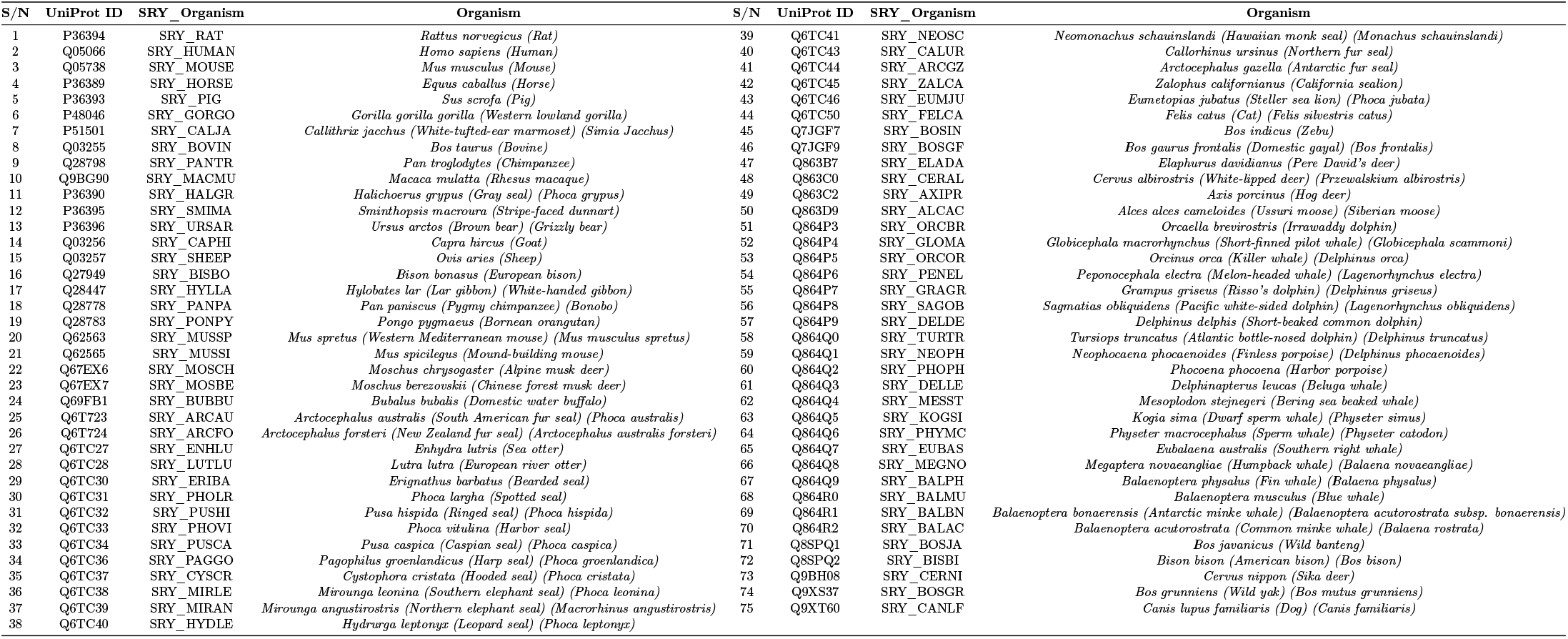
List of organisms from which SRY protein sequences were obtained.

**Table 2:**
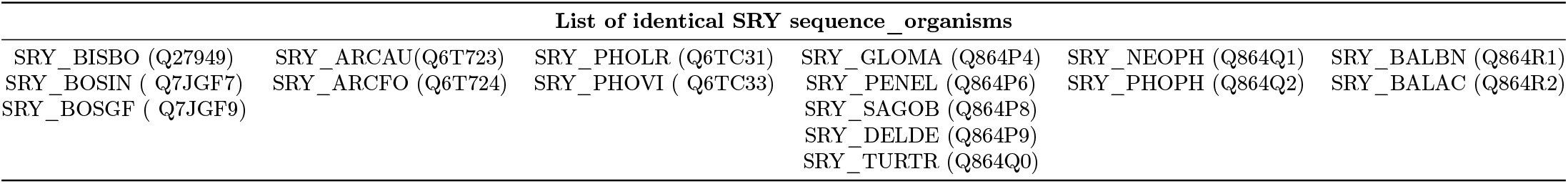
List of organisms in which identical SRY protein sequences were detected. Each column represents a distinct set of organisms in which identical SRY sequences were identified.

**Table 3:**
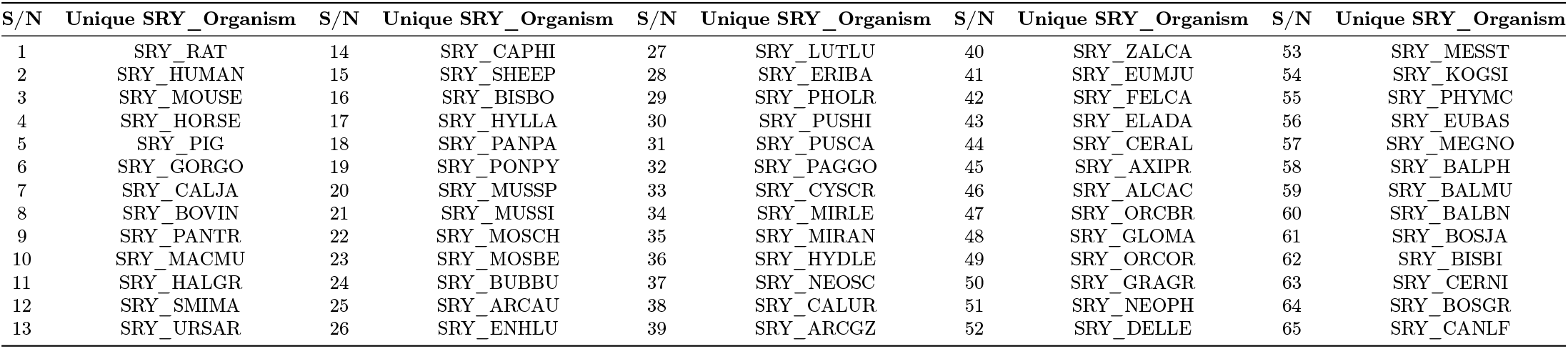
List of organisms from which sixty-five unique SRY protein sequences were identified.

From the sixty-five organisms examined, the high-mobility group box (HMG-box) subsequences were extracted from their corresponding SRY protein sequences. A total of thirty-eight unique HMG-box subsequences were identified (Table 4), whereas the remaining subsequences were found to be identical and are listed in Table 5. Henceforth, HMG-box domains derived from SRY sequences are referred to as ‘SRY-HMG’ and are designated as ‘SRY_organism_HMG’, where organism identifies the source organism.

**Table 4:**
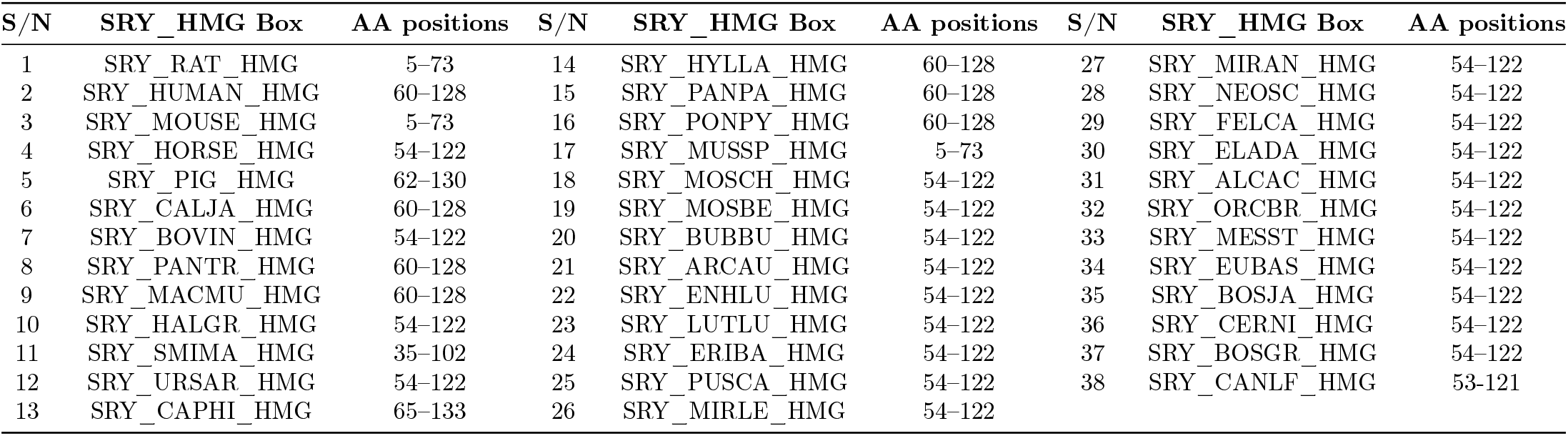
List of unique SRY-HMG subsequences, their associated organisms, and the corresponding amino acid positions within the SRY sequences.

**Table 5:**
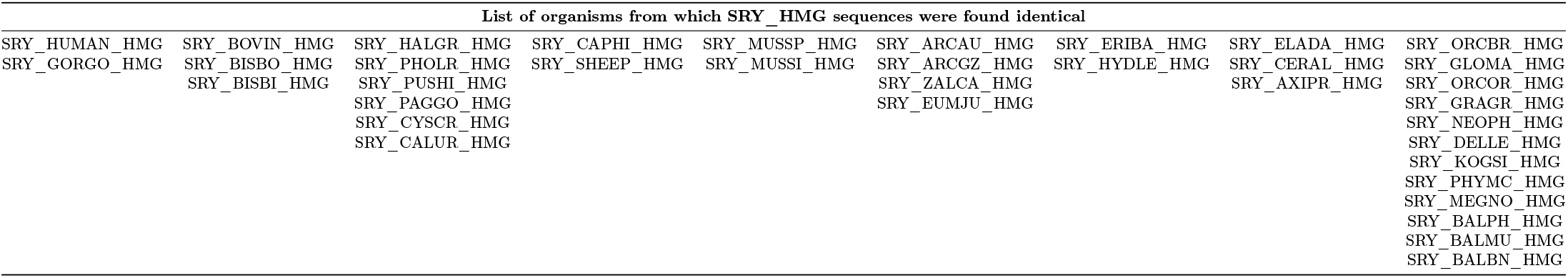
List of organisms with identical SRY-HMG subsequences. Each column represents a distinct set of organisms in which identical HMG subsequences were identified.

Therefore, in this study, sixty-five SRY protein sequences together with thirty-eight SRY-HMG subsequences were considered for subsequent analyses.

## 3. Methods

### 3.1. Amino acid composition based analyses of SRY and SRY-HMG sequences

#### 3.1.1. Amino acid relative frequency and correlation analysis across SRY sequences and SRY-HMG subsequences

The amino acid composition of each SRY sequence and its corresponding SRY-HMG was quantified by determining the frequency of occurrence of the twenty standard amino acids [17]. To normalize for variations in sequence length, the frequency of each amino acid was expressed as a percentage of the total sequence length, yielding its relative abundance within the corresponding SRY or SRY-HMG sequence [18, 19]. Thus, every SRY and SRY-HMG sequence was represented as a 20-dimensional feature vector comprising the relative frequencies of the standard amino acids.

Pearson correlation coefficients were subsequently computed between amino acid frequency profiles to evaluate pairwise associations among amino acid abundances [20]. This analysis enabled the identification of positively and negatively correlated amino acid pairs, providing insights into compositional relationships and patterns of amino acid usage across the SRY and SRY-HMG sequences.

#### 3.1.2. Dendrogram relationship of SRY and SRY-HMG sequences based on amino acid composition

To investigate compositional relationships among the SRY protein sequences and their corresponding SRY-HMG, each sequence was represented as a 20-dimensional vector comprising the relative frequencies of the standard amino acids [20, 21]. Pairwise Aitchison distances were then calculated between all sequence vectors to quantify compositional dissimilarities, while accounting for the compositional nature of amino acid frequency data. The resulting distance matrices were symmetric with dimensions 65 *×* 65 for the SRY sequences and 38 *×* 38 for the SRY-HMG subsequences, respectively, thereby summarizing the pairwise compositional differences among the sequences. Using these Aitchison distance matrices, dendrograms were constructed with the MATLAB phytree function to visualize the compositional clustering and dendrogram relationships among the SRY sequences and SRY-HMG domains.

#### 3.1.3. Shannon entropy-based analysis of compositional diversity in SRY and SRY-HMG sequences

Sequence compositional diversity in SRY protein sequences and their corresponding SRY-HMG was quantified using Shannon entropy, a measure of uncertainty derived from amino acid frequency distributions [22, 23]. For a given protein sequence, let *p_i_* denote the relative frequency of the *i*th amino acid (*i* = 1*, . . . ,* 20). The Shannon entropy (*H*) is computed as

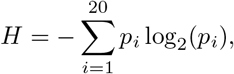

Shannon entropy provides a quantitative measure of amino acid compositional diversity within a sequence. Lower entropy values indicate reduced diversity and a predominance of specific amino acids, whereas higher values reflect a more uniform distribution of residues. For protein sequences composed of the twenty standard amino acids, the theoretical maximum entropy is 4.322, which occurs when all amino acids are equally represented [22, 24]. Thus, Shannon entropy serves as an effective metric for comparing compositional complexity and conservation among SRY and SRY-HMG sequences.

### 3.2. Frequency-dominant amino acid distribution across consecutive sequence windows in SRY and SRY-HMG sequences

Each SRY sequence was partitioned into consecutive, non-overlapping 20-residue windows to investigate local amino acid composition patterns [20]. For SRY sequences whose terminal segment contained fewer than five residues, the remaining residues were merged with the preceding window to avoid biases associated with very short segments. Terminal segments containing more than five residues were treated as independent windows.

For each window, the frequencies of the 20 standard amino acids were calculated, and the amino acid with the highest occurrence was identified as the *dominant residue*. The dominance of a residue within a window was quantified as

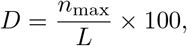

where *n*_max_ denotes the frequency of the most abundant amino acid in the window and *L* represents the length of the corresponding window. Thus, *D* measures the percentage contribution of the dominant amino acid to the local SRY sequence composition.

For every window, the dominant amino acid, its percentage abundance, and its positions within the original sequence were recorded. The resulting dominance profiles were visualized as window-wise distributions, in which the dominant amino acid percentage was plotted across consecutive sequence windows. To facilitate direct comparison among different sequences, all plots were generated using a standardized y-axis scale ranging from 0% to 100%. Background color coding was employed to distinguish dominant amino acid types across windows, thereby enabling visualization of local compositional heterogeneity and residue dominance patterns within the SRY sequences.

In the similar fashion, each SRY-HMG subsequences was partitioned into consecutive non-overlapping windows of 10 amino acid residues and analysis was carried out to determine frequency-dominant amino acid in each window.

### 3.3. Multiple sequence alignment-based phylogenetic analysis of SRY and SRY-HMG sequences

Evolutionary relationships among the SRY proteins and their corresponding SRY-HMG were investigated using multiple sequence alignment performed with Clustal Omega [25]. The alignments facilitated the identification of conserved residues, variable positions, and amino acid substitutions across the analyzed sequences, thereby providing a framework for assessing sequence homology and evolutionary divergence [26]. Multiple sequence alignments were generated using the default settings of Clustal Omega [26]. Amino acid substitutions were scored using the *Gonnet* substitution matrix, with a gap opening penalty of 10 and a gap extension penalty of 0.20. End-gap penalties were not applied, in accordance with the standard alignment procedure implemented by the Clustal Omega. The resulting alignments were subsequently used to infer phylogenetic relationships and to examine sequence conservation patterns among the SRY and SRY-HMG sequences.

### 3.4. Compilation of clinically significant variants and phenotype-associated mutations in human SRY

Clinically significant variants of SRY_HUMAN were compiled from Ensembl, ClinVar, and UniProt. Each variant was recorded with its amino acid change, mutation type, and reported clinical significance (pathogenic or likely pathogenic). Variant positions were mapped onto the SRY_HUMAN sequence to examine their distribution relative to the HMG box and the NTD and CTD. A separate set of NTD and CTD mutations with proband phenotypes was assembled from published clinical reports. For each mutation, the affected domain, proband karyotype and sex, and clinical phenotype were recorded. Premature stop codons in the NTD were excluded.

### 3.5. Prediction of liquid-liquid phase separation propensity for SRY and SRY-HMG sequences

The propensity for spontaneous liquid-liquid phase separation (LLPS) was predicted for the full-length SRY sequences and the SRY-HMG subsequences using the FuzDrop web server [15, 27, 16]. SRY/SRY-HMG sequence was submitted to the FuzDrop server with default parameters.

For every input sequence, the server returns a probability score for spontaneous phase separation (*p_LLP S_*), identifies droplet-promoting regions (DPRs), aggregation hot-spots, and regions with context-dependent interactions (*S_bind_*). Following the classification scheme of Hardenberg et al. [15], sequences with *p_LLP S_ ≥* 0.60 were considered droplet drivers, while those with lower scores were considered droplet clients or non-LLPS proteins. The positions of the predicted DPRs and aggregation hot-spots were recorded, and the *S_bind_* regions were noted as indicators of context-dependent binding modes. The analysis was performed separately for the full-length SRY_HUMAN and respective SRY_HUMAN_HMG set to compare their LLPS propensities and to map the droplet-promoting features onto the sequence architecture.

### 3.6. Dendrogram relationships among SRY and SRY-HMG sequences based on dipeptide composition analysis

Dipeptide composition was calculated for each SRY/SRY-HMG protein sequence to capture the frequency of adjacent amino acid pairs. For a protein of length *L*, this yields a 400-dimensional vector representing all possible ordered dipeptides from the twenty standard amino acids. Overlapping dipeptides were extracted across each sequence, and the frequency of each dipeptide *ij* was computed as:

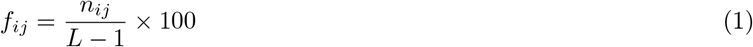

where *n_ij_* is the count of dipeptide ‘*ij*’ and *L −* 1 is the total number of dipeptide bonds.

Dipeptide prevalence was assessed across all SRY/SRY-HMG sequences using the dipeptide compositions across SRy/SRY-HMG sequences. For each dipeptide, presence was defined as a non-zero frequency in a given sequence. The following statistics were calculated: the number of sequences containing the dipeptide, prevalence (%), mean, median, standard deviation, minimum, and maximum frequency (%).

Prevalence was computed as:

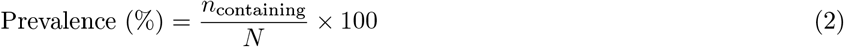

where *n*_containing_ is the number of sequences containing the dipeptide and *N* is the total number of SRY/SRY-HMG sequences. Dipeptides were then classified into five prevalence categories: absent (0%), rare (*≤*25%), moderately prevalent (26–50%), common (51–75%), and highly prevalent (>75%). Dipeptides absent from all sequences were identified separately. In addition, pairwise Aitchison distances were computed as Euclidean distances between the dipeptide frequency vectors for each pair of SRY/SRY-HMG sequences. Hierarchical clustering used average linkage, and a dendrogram was generated for SRY and SRY-HMG sequences.

### 3.7. Longest polystring analysis and dendrogram relationships across SRY and SRY-HMG sequences

Homopolymer runs were characterized by computing, for each of the 20 standard amino acids across all SRY and SRY-HMG sequences, the longest polystring, defined as the longest consecutive stretch of a single amino acid residue. The longest run for each amino acid was then identified; where multiple runs shared the same maximum length, the first occurrence was retained. For each amino acid in each SRY/SRY-HMG sequence, the longest run length, the corresponding polystring, a normalized longest-run percentage were recorded, the latter computed as *L*_run_*/L ×* 100, where *L*_run_ is the longest run length for that amino acid and *L* is the total sequence length.

Based on the the normalized longest polystring lengths for SRY/SRY-HMG sequences, a UPGMA dendrogram was constructed. The normalized percentages for each of the 20 amino acids were assembled into a 20-dimensional feature vector per sequence, and pairwise Euclidean distances were calculated from this matrix to generate a full distance matrix. Hierarchical clustering was performed using average linkage, and the fidelity of the resulting dendrogram was assessed. Clusters were defined by cutting the dendrogram at a distance threshold of 0.5, and cluster membership was recorded for each sequence. Branch colouring was applied using the same threshold; no threshold line was drawn, as the cutoff was used solely for visual grouping and cluster assignment.

### 3.8. Hydropathy-based n-gram frequency analysis of SRY and SRY-HMG sequences

Local hydropathy patterns were captured by partitioning the 20 standard amino acid residues (Σ) into two disjoint sets:

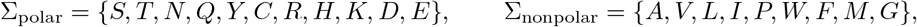

such that Σ = Σ_polar_ *∪* Σ_nonpolar_ and Σ_polar_ *∩* Σ_nonpolar_ = Ø. Each residue *x_i_* in a sequence *x* = (*x*_1_*, x*_2_*, . . . , x_L_*) of length *L* was then encoded as a binary indicator:

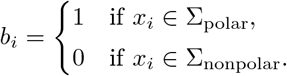

An *n-gram* was defined as any contiguous subsequence of length *n* in this binary encoding, i.e. a word in *{*0, 1*}^n^* representing a local hydropathy pattern. For each sequence, all overlapping *n*-grams were enumerated by sliding a window of length *n* across the binary sequence, yielding the complete word set *W_n_* = *{*0, 1*}^n^* of size 2*^n^*. For each word *w ∈ W_n_*, the count and relative frequency were defined as

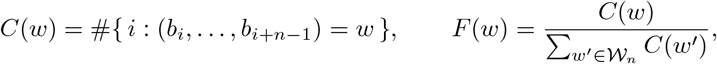

so that *F* (*w*) represents the proportion of *w* among all observed overlapping *n*-grams.

As an illustration, consider the binary sequence *b* = (0, 1, 0, 0, 1, 1, 0, 1, 0, 0, 1, 0). Sliding a window of length 2 yields the overlapping 2-grams 01, 10, 00, 01, 11, 10, 01, 10, 00, 01, 10, with counts *C*(00) = 2, *C*(01) = 4, *C*(10) = 4, *C*(11) = 1 and relative frequencies *F* (00) = 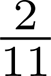, *F* (01) = 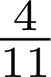, *F* (10) = 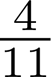 *F* (11) = 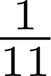. Similarly, sliding a window of length 3 yields 010, 100, 001, 011, 110, 101, 010, 100, 001, 010, with counts *C*(010) = 3, *C*(100) = 2, *C*(001) = 2, *C*(011) = 1, *C*(110) = 1, *C*(101) = 1 and the remaining 3-grams having count zero. The same procedure generalises to larger *n*.

Finally, using the relative frequencies of independent *n*-grams (*n* = 2, 3, and 4), pairwise Euclidean distances were computed between all SRY sequences and between all SRY-HMG subsequences. This produced square, symmetric distance matrices of dimension 65 *×* 65 and 38 *×* 38, respectively, representing the compositional dissimilarities among the 65 SRY sequences and the 38 SRY-HMG subsequences. Hierarchical clustering was then applied to each distance matrix, and dendrograms were constructed to visualise sequence proximities. Proximal clusters were identified by applying a distance threshold to the dendrogram, thereby delineating groups of sequences with similar hydropathy-based *n*-gram profiles.

### 3.9. Hydropathy profile-based clustering of SRY and SRY-HMG sequences

For each SRY/SRY-HMG sequence, the percentages of polar and nonpolar residues were calculated. In addition, the longest continuous stretch of polar residues and the longest continuous stretch of nonpolar residues were identified by scanning the sequence for the maximal uninterrupted run of residues belonging to each set. For every SRY/SRY-HMG sequence, the longest polar and nonpolar stretches were recorded as absolute lengths and then normalized by dividing by the total sequence length to account for variation in sequence size. These normalized values were used to construct a two-dimensional feature vector for each sequence. The feature vectors were then subjected to density-based clustering using DBSCAN with Euclidean distance. The parameters *ɛ* (neighborhood radius=0.05) and Minpts (minimum points per cluster=3) were optimized empirically to balance sensitivity and cluster stability. DBSCAN was chosen because it does not require pre-specification of the number of clusters and can identify outliers as noise, thereby enabling robust grouping of SRY and SRY-HMG sequences based solely on their longest polar and nonpolar domain architecture.

### 3.10. Intrinsic disorder profiles analysis of SRY and SRY-HMG sequences

To evaluate intrinsic disorder, the Rapid Intrinsic Disorder Analysis Online (RIDAO) platform was employed, utilizing six distinct computational models (PONDR® VLXT, PONDR® VL3, PONDR® VSL2B, IUPred short/long, and PONDR® FIT). A consensus mean disorder prediction (MDP) was subsequently derived by calculating the average output across all individual predictors [28]. Protein residues were classified based on their disorder scores: disordered (0.5–1.0), flexible (0.15–0.5), or ordered (<0.15). The per-residue outputs generated by these tools are used to calculate the average disorder score (ADS) and percent of intrinsically disordered residues (PPIDR). These parameters are used for classification disorder status of query proteins. Here, PPIDR values classify proteins as mostly ordered (<10%), moderately disordered (10–30%), or highly disordered (>30%), while ADS values provide a complementary classification, where ADS < 0.15, 0.15 *≤* ADS < 0.5, and ADS *≥* 0.5 correspond to highly ordered, moderately disordered/flexible, and highly disordered regions, respectively [29, 30]. Because PPID and ADS measure different aspects of protein structures, combining them creates a highly effective method for classifying tricky or edge-case proteins. This combined approach introduces two additional “plus” categories: “mostly ordered plus” (low PPIDR < 10% and high ADS *≥* 0.15) and “moderately disordered plus” (moderate ADS between 0.15 and 0.5, but high PPIDR > 30%). These specific classifications help differentiating between proteins featuring only a few highly disordered regions and those with widespread, slightly unstable characteristics.

3D structures of SRY_ENHLU, SRY_HUMAN, and SRY_MOUSE were modeled by the state-of-the-art AI-based tool AlphaFold 2 developed by Google DeepMind, which generates the “expected” structure of a query protein in a form of a static snapshot corresponding to the lowest-energy, most stable 3D conformation [31]. In other words, it generates highly precise static snapshots rather than modeling the dynamic, moving processes of proteins in action. To address this limitation and generate a conformational ensemble rather than a single static structure, we utilized the AFflecto web server [32]. AFflecto produces comprehensive conformational ensembles encompassing both structured domains and intrinsically disordered regions (IDRs) directly from AlphaFold (AF) input models [32]. The platform leverages pLDDT scores and structural context of AFflecto to identify IDRs, categorizing them into loops, linkers, or tails. To map the conformational landscape of these flexible segments, the server applies highly efficient stochastic sampling algorithms [32]. Additionally, AFflecto features a built-in mechanism to detect conditionally folded IDRs that might be erroneously predicted as rigid elements by AlphaFold. Consequently, the resulting ensembles offer a biologically realistic representation of protein structural heterogeneity [32]. In our analysis, we used the default AFflecto parameters, such as PLDDT cutoff of 0.7, contact distance of 8, and shortest rigid of 3, to generate 25 models for each protein.

### 3.11. Determining proximal sets of SRY/SRY-HMG subsequences

For each SRY or SRY-HMG sequence, six types of quantitative features were computed: (i) amino acid frequency distribution, (ii) amino acid sequence homology, (iii) dipeptide composition, (iv) longest polystring analysis, (v) polar and nonpolar frequency distribution, and (vi) hydropathy-based *n*-gram frequency distribution. Each feature type yielded a feature vector for every sequence, with the dimensionality depending on the feature (e.g., 20 for longest polystring of amino acids, 2 for polar/nonpolar composition, and so on).

To compare sequences, pairwise Euclidean distances were calculated between their feature vectors for each feature type separately. For a given feature, the distance between two sequences reflected their dissimilarity in that feature space. These distances were then used to identify SRY/SRY-HMG subsequences that were proximal to one another. For each feature, a suitable distance threshold was applied to define the most proximal sets of sequences. Sequences falling within the threshold were considered part of the same proximal set for that feature.

### 3.12. Cumulative feature-based clustering of SRY and SRY-HMG sequences

Robust proximal SRY and SRY-HMG sequences were identified by performing eight independent feature-based clustering analyses. The features used were amino acid frequency, sequence homology, dipeptide composition, longest polystring, 2-, 3-, and 4-gram hydropathy profiles, and longest polar/nonpolar stretches. For each feature, sequences were clustered using hierarchical clustering with Euclidean distance and average linkage, and clusters were defined at a fixed threshold. The clusters obtained from all eight analyses were then intersected, and only those groups of sequences that co-clustered in all eight analyses were retained as cumulative proximal clusters. This intersection ensured that the final clusters were robust to the choice of feature and reflected consistent sequence similarity across multiple compositional and hydropathy descriptors. Sequences that did not co-cluster with any other sequence in all eight analyses were retained as singletons. This integrative approach provides a multidimensional perspective on sequence similarity, capturing complementary signals from composition, hydropathy, and local order. The analysis was performed separately for the 65 full-length SRY sequences and the 38 SRY-HMG subsequences.

## 4. Results and analyses

### 4.1. Amino acid frequency distribution across SRY and SRY-HMG

#### 4.1.1. Amino acid frequency distribution across SRY sequences

Amino acid frequency analysis of 65 SRY sequences from 65 organisms revealed distinct enrichment patterns. Four sequences—SRY_MOUSE, SRY_RAT, SRY_MUSSP, and SRY_MUSSI—showed the highest glutamine frequency. Notably, SRY_MOUSE uniquely contained a polyglutamine tract that stabilized the protein and functioned as a transcriptional activation domain, essential for Sox9 upregulation and male sex determination [11]. Glutamine-rich sequences conferred protein stability and transcriptional activation capacity, enabling organism-specific regulation [11]. In contrast, only one sequence, SRY_SMIMA, was lysine-rich (Figure 1). Lysine acetylation had been shown to regulate nuclear import of SRY, whereas deacetylation promoted its cytoplasmic export [33, 34]. In line with this, the lysine-rich sequence SRY_SMIMA was likely to enhance chromatin interaction and nuclear localization, providing a unique layer of regulatory control in this organism. Another subset of SRY sequences exhibited arginine enrichment, specifically SRY_HORSE, SRY_PIG, SRY_ENHLU, SRY_ERIBA, SRY_ORCBR, SRY_GLOMA, SRY_ORCOR, SRY_GRAGR, SRY_NEOPH, SRY_DELLE, SRY_MESST, SRY_KOGSI, SRY_PHYMC, SRY_EUBAS, SRY_MEGNO, SRY_BALPH, and SRY_BALMU (Figure 1). This observation was consistent with established findings that arginine residues within the HMG box mediated minor groove binding and induced sharp DNA bending, a mechanism critical for transcriptional activation of SOX9 [6, 35]. Accordingly, the enrichment of arginine in these sequences was inferred to strengthen DNA binding affinity, thereby ensuring robust sex-determining activity across these organisms [6, 35]. The remaining sequences predominantly displayed serine-rich profiles, underscoring serine as the most frequent amino acid across the majority of SRY proteins from various organisms. This pattern corresponded with established findings that serine residues frequently acted as phosphorylation sites, facilitating dynamic modulation of protein activity [36]. The predominance of serine therefore indicated that SRY proteins were subject to extensive post-translational regulation, enabling precise temporal control of their function during sex determination [36].

**Figure 1:**
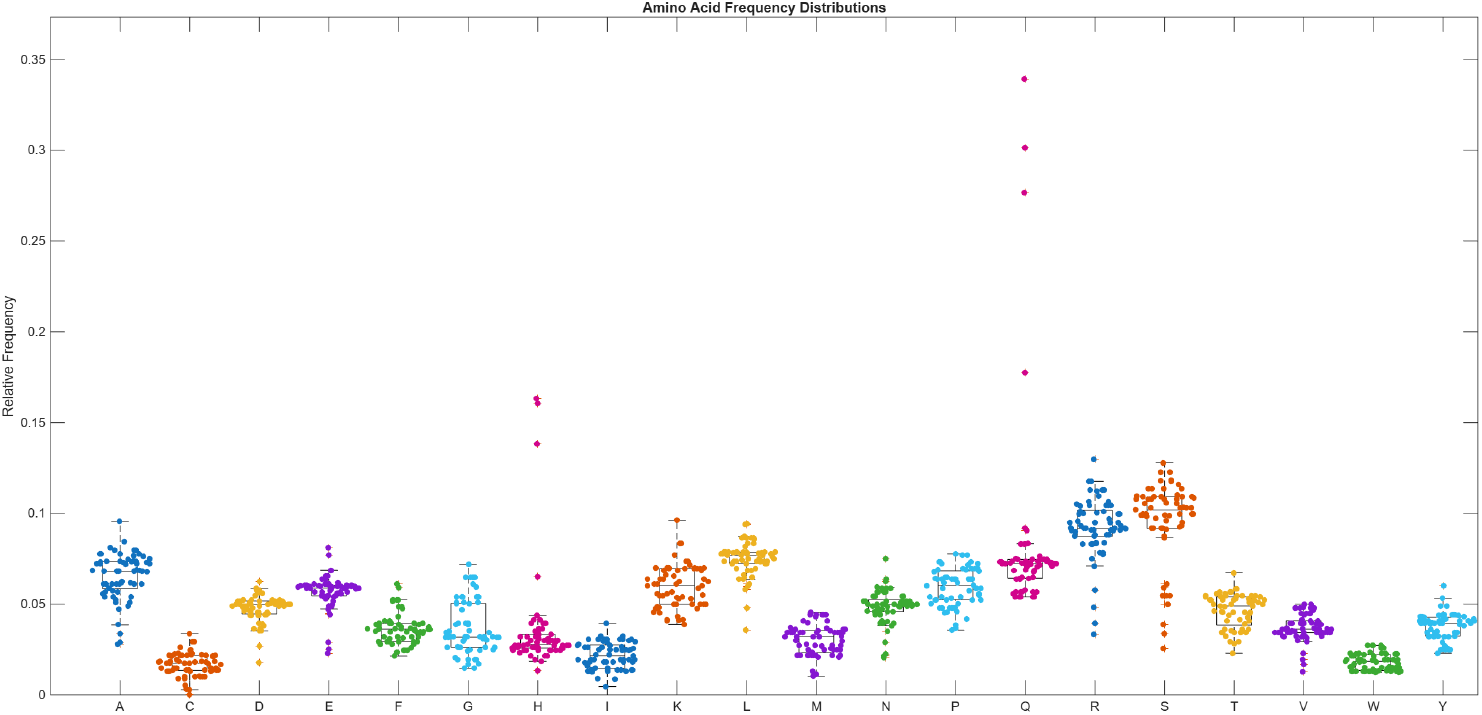
Boxplot representation of amino acid frequency distributions in 38 SRY-HMG subsequences. The central line denotes the median, the box represents the interquartile range, whiskers indicate data spread, and points correspond to frequencies observed in individual subsequences.

Analysis of amino acid frequencies across all SRY sequences also revealed distinct depletion patterns. SRY_PANTR had the lowest glycine content among all SRY proteins analyzed. This observation was consistent with reports that glycine residues conferred structural flexibility, whereas their depletion enforced rigidity within DNA-binding domains. Accordingly, the scarcity of glycine in primate SRY suggested organism-specific structural constraints that potentially influenced the conformational dynamics required for sex-determining activity. Six SRY sequences—SRY_URSAR, SRY_PANPA, SRY_MIRLE, SRY_MIRAN, SRY_HYDLE, and SRY_NEOSC—had the lowest isoleucine content (Figure 1). This finding corresponded with established evidence that isoleucine contributed to hydrophobic core stability, and its depletion reflected reduced reliance on hydrophobic packing [37]. Consequently, the scarcity of isoleucine in these sequences was inferred to favor greater surface exposure of DNA-binding motifs, potentially enhancing accessibility and regulatory interactions during sex determination [38]. A number of SRY sequences, including those from rodents and whales, namely SRY_RAT, SRY_MOUSE, SRY_CALJA, SRY_HALGR, SRY_MUSSP, SRY_MUSSI, SRY_ARCAU, SRY_LUTLU, SRY_ERIBA, SRY_PHOLR, SRY_PUSHI, SRY_PUSCA, SRY_PAGGO, SRY_CYSCR, SRY_CALUR, SRY_ARCGZ, SRY_ZALCA, SRY_EUMJU, SRY_ORCBR, SRY_GLOMA, SRY_ORCOR, SRY_GRAGR, SRY_NEOPH, SRY_DELLE, SRY_MESST, SRY_KOGSI, SRY_PHYMC, SRY_EUBAS, SRY_MEGNO, SRY_BALPH, and SRY_BALMU, had the lowest cysteine content. Cysteine residues often formed disulfide bonds, and their absence indicated reliance on non-covalent stabilization mechanisms. Consequently, the depletion of cysteine across these SRY sequences supported the presence of a flexible protein architecture, favoring dynamic conformational changes over rigid stabilization in SRY function [39, 40]. In the majority of SRY sequences, tryptophan residues were markedly depleted (Figure 1). A conserved tryptophan within the HMG box was indispensable for DNA binding, and mutations at this site impaired sex determination [41]. The widespread depletion of tryptophan outside the HMG domain therefore suggested a streamlined protein design, minimizing bulky aromatic residues while retaining the essential DNA-binding tryptophan required for functional activity [42, 43, 44].

Pairwise correlation analysis of amino acid frequencies across the 65 SRY sequences revealed strong positive associations between glutamine and histidine (Q-H, *r* = 0.9475), leucine and serine (L-S, *r* = 0.6516), arginine and glycine (R-G, *r* = 0.6506), and cysteine and serine (C-S, *r* = 0.6005) (Figure 2). In contrast, strong negative correlations were observed for glutamine and glutamic acid (Q-E, *r* = *−*0.7885), glutamic acid and histidine (E-H, *r* = *−*0.7744), arginine and glutamine (R-Q, *r* = *−*0.7553), arginine and histidine (R-H, *r* = *−*0.7011), glutamine and serine (Q-S, *r* = *−*0.6806), glutamine and leucine (Q-L, *r* = *−*0.6403), asparagine and histidine (N-H, *r* = *−*0.6386), and histidine and serine (H-S, *r* = *−*0.6317) (Figure 2). Furthermore, it was noteworthy that these correlations provide a descriptive overview of amino acid co-variation within the SRY sequences but do not, by themselves, imply direct evolutionary, structural, or functional coupling between residues.

**Figure 2:**
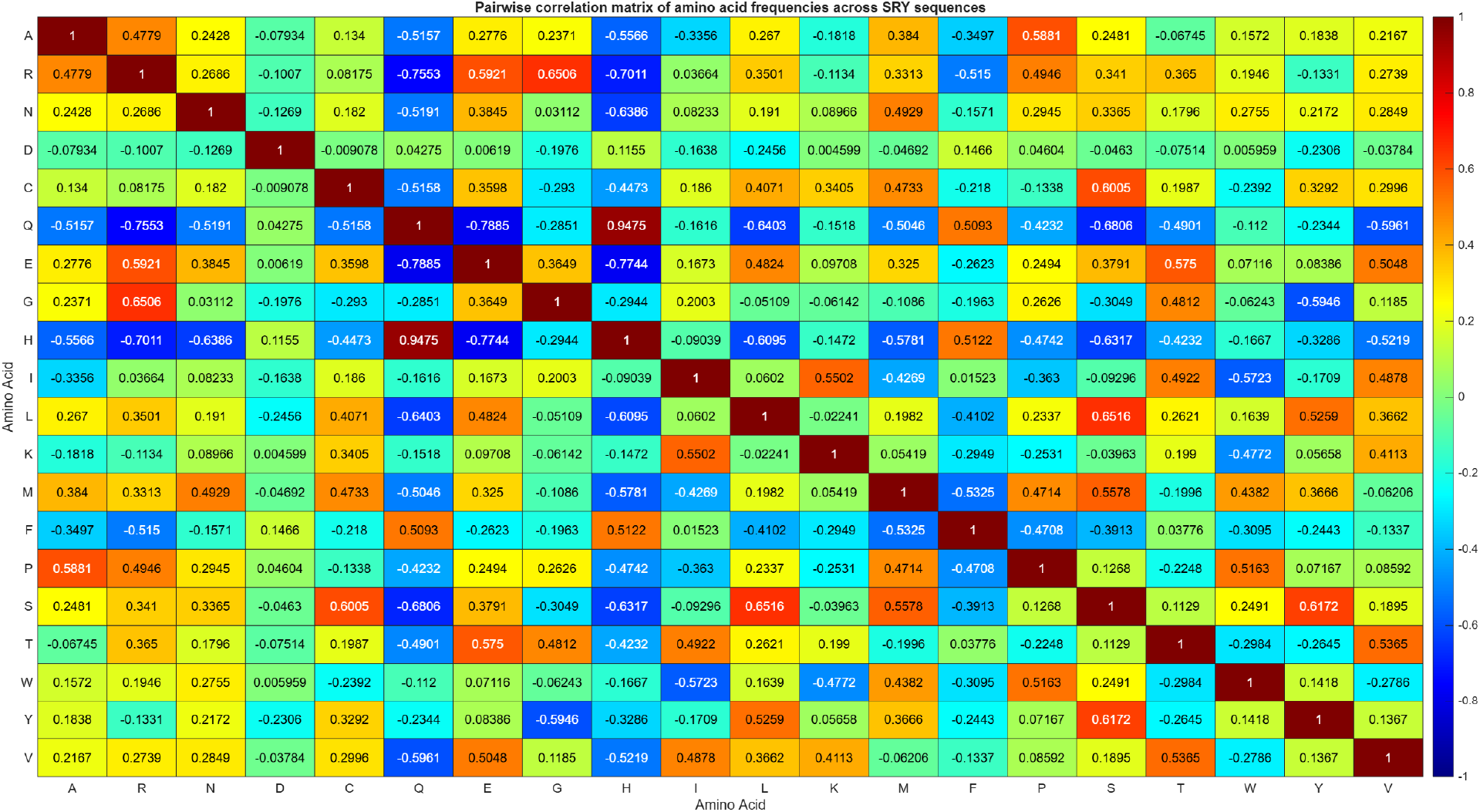
Heatmap showing the pairwise Pearson correlation coefficients among amino acid frequencies in 65 SRY sequences. Color intensity represents the magnitude and direction of the correlations, with values ranging from *−*1 (negative correlation) to 1 (positive correlation).

A dendrogram illustrating the relationships among the 65 SRY sequences was constructed based on the similarity of their amino acid frequency profiles (Figure 3). Using a distance threshold cutoff of 1.0, the SRY sequences from the 65 organisms were grouped into 13 distinct clusters, as summarized in Table 6. Furthermore, it was noted that closely related SRY sequences belonging to the same cluster were distinguished by different colors in the dendrogram (Figure 3). A total of 46 sequences were included within these clusters, while the remaining 19 sequences remained unclustered and were considered outliers according to the selected distance threshold.

**Figure 3:**
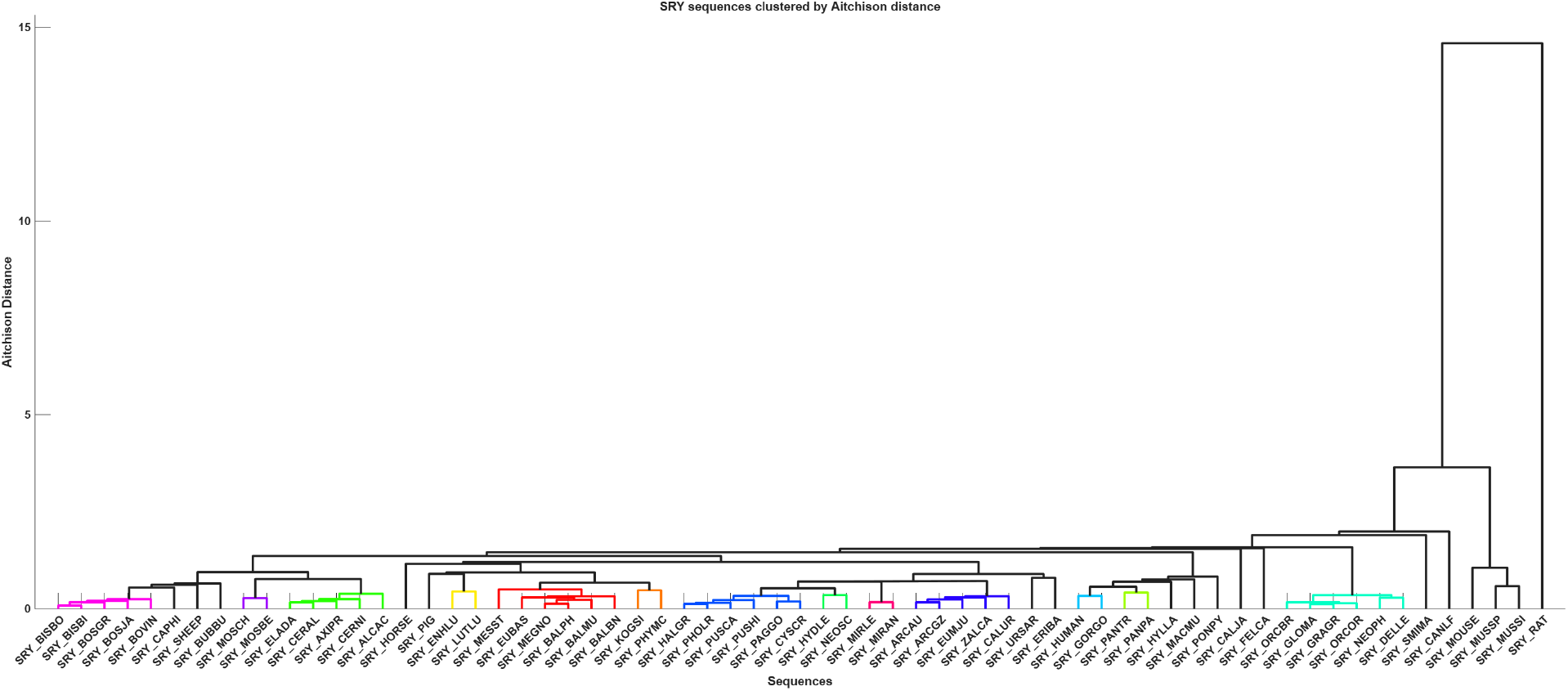
Hierarchical clustering of 65 SRY sequences based on amino acid frequency composition profiles using the Aitchison distance metric. Distinct clusters identified at a distance threshold of 1.0 are highlighted in different colors, whereas unclustered sequences are shown in black. The vertical axis represents the Aitchison distance, reflecting compositional dissimilarity among the SRY sequences.

**Table 6:**
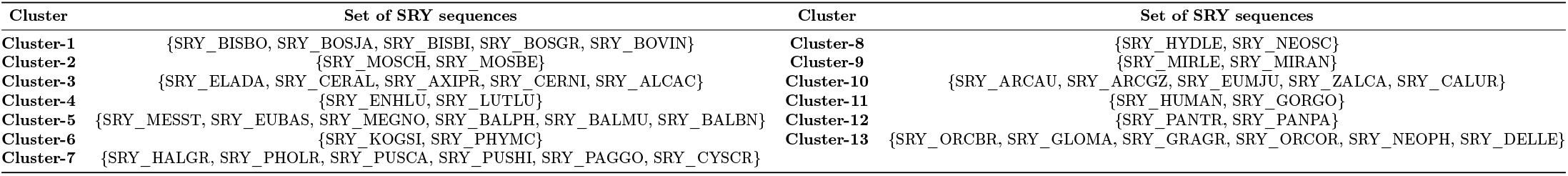
List of proximal clusters based on proximity derived from amino acid frequency profile of 65 SRY sequences.

#### 4.1.2. Amino acid frequency distribution across SRY-HMG subsequences

Amino acid frequency analysis of the 38 SRY-HMG subsequences derived from 65 organisms revealed distinct enrichment patterns, similar to those observed for the full-length SRY sequences (Figure 4). Lysine was the most abundant amino acid in 14 SRY-HMG subsequences, namely SRY_RAT_HMG, SRY_HUMAN_HMG, SRY_MOUSE_HMG, SRY_HORSE_HMG, SRY_PIG_HMG, SRY_CALJA_HMG, SRY_BOVIN_HMG, SRY_PANTR_HMG, SRY_SMIMA_HMG, SRY_HYLLA_HMG, SRY_PANPA_HMG, SRY_PONPY_HMG, SRY_MUSSP_HMG, and SRY_MACMU_HMG. On the other side, glutamine exhibited the highest frequency in only three SRY-HMG subsequences, namely SRY_HALGR_HMG, SRY_URSAR_HMG, and SRY_FELCA_HMG. The remaining 21 SRY-HMG subsequences exhibited arginine enrichment, with arginine being the most abundant amino acid in each case.

**Figure 4:**
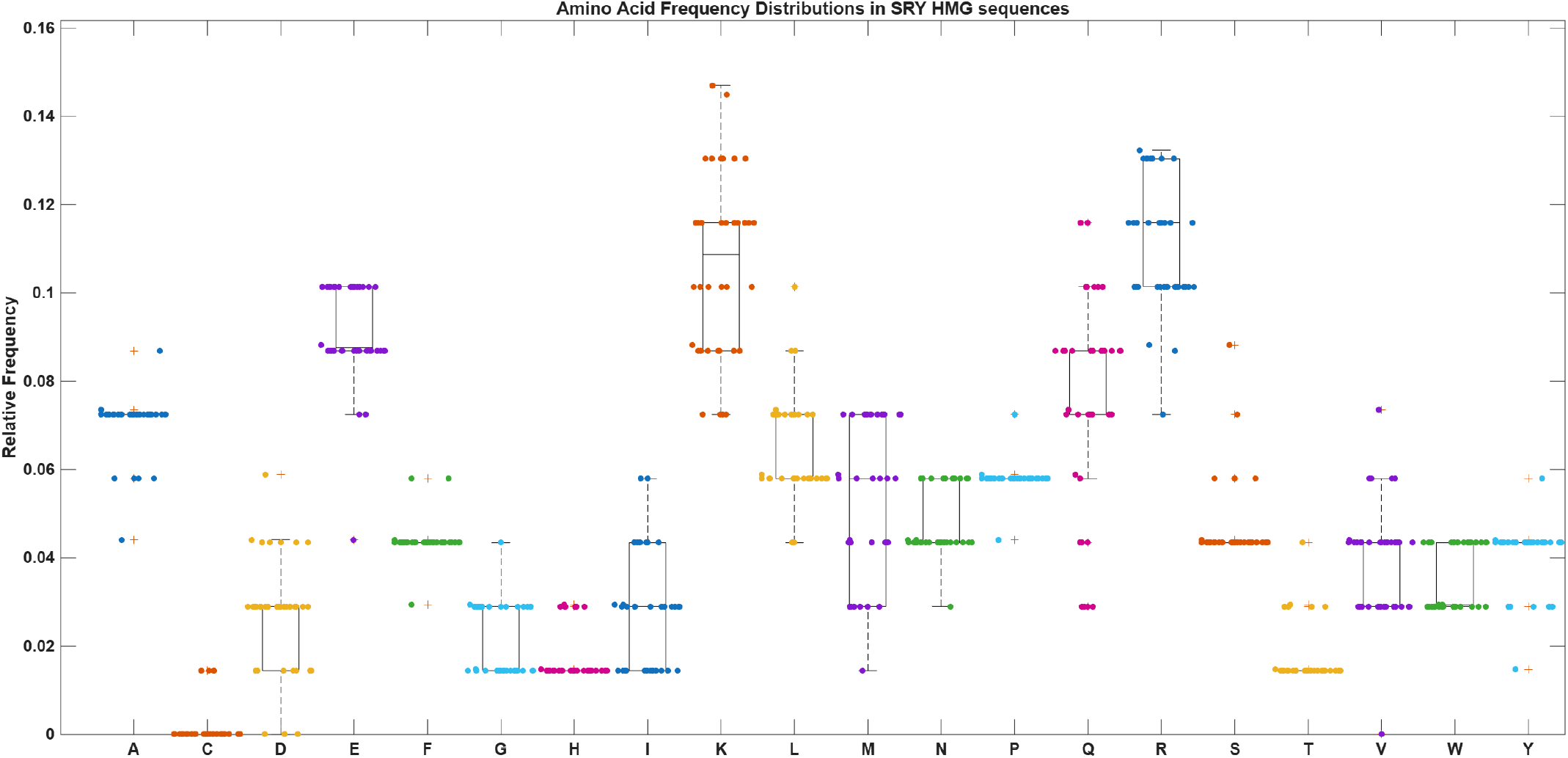
Boxplot representation of amino acid frequency distributions in 38 SRY-HMG subsequences. The central line denotes the median, the box represents the interquartile range, whiskers indicate data spread, and points correspond to frequencies observed in individual subsequences.

Among the 38 SRY-HMG subsequences, aspartic acid was the least represented amino acid, with a frequency of 0%, only in SRY_RAT_HMG, SRY_MOUSE_HMG, and SRY_MUSSP_HMG (Figure 4). Given the conserved role of the HMG box in DNA binding and male sex determination, the absence of aspartic acid in these rodent sequences suggests that this residue is not strictly required for maintenance of HMG-domain function in all mammalian lineages [45, 46]. In SRY_HORSE_HMG and SRY_PIG_HMG, cysteine, glycine, isoleucine, and threonine shared the lowest frequency of 1.4% each (Figure 4). The remaining SRY-HMG subsequences were devoid of cysteine residues, indicating that cysteine is not universally required for maintenance of the highly conserved HMG domain responsible for DNA binding and sex-determining activity [6, 35]. Its occurrence in only a small number of sequences may therefore reflect lineage-specific sequence variation rather than a core functional requirement.

Pairwise correlation analysis of amino acid frequencies in the 38 SRY-HMG subsequences revealed several notable associations (Figure 5). Strong positive correlations were observed between glutamine and methionine (Q-M, *r* = 0.7341), methionine and tryptophan (M-W, *r* = 0.6870), histidine and threonine (H-T, *r* = 0.6507), glutamine and tryptophan (Q-W, *r* = 0.6362), serine and threonine (S-T, *r* = 0.6229), and alanine and phenylalanine (A-F, *r* = 0.6217). In contrast, several amino acid pairs exhibited strong negative correlations, most notably alanine and serine (A-S, *r* = *−*0.9069). Additional negatively correlated pairs included glycine and methionine (G-M, *r* = *−*0.7595), asparagine and aspartic acid (N-D, *r* = *−*0.7068), glutamine and isoleucine (Q-I, *r* = *−*0.6824), leucine and methionine (L-M, *r* = *−*0.6545), glutamic acid and lysine (E-K, *r* = *−*0.6488), alanine and threonine (A-T, *r* = *−*0.6404), glutamine and arginine (Q-R, *r* = *−*0.6373), histidine and tyrosine (H-Y, *r* = *−*0.6321), glycine and tryptophan (G-W, *r* = *−*0.6290), and isoleucine and tryptophan (I-W, *r* = *−*0.6226) (Figure 5).

**Figure 5:**
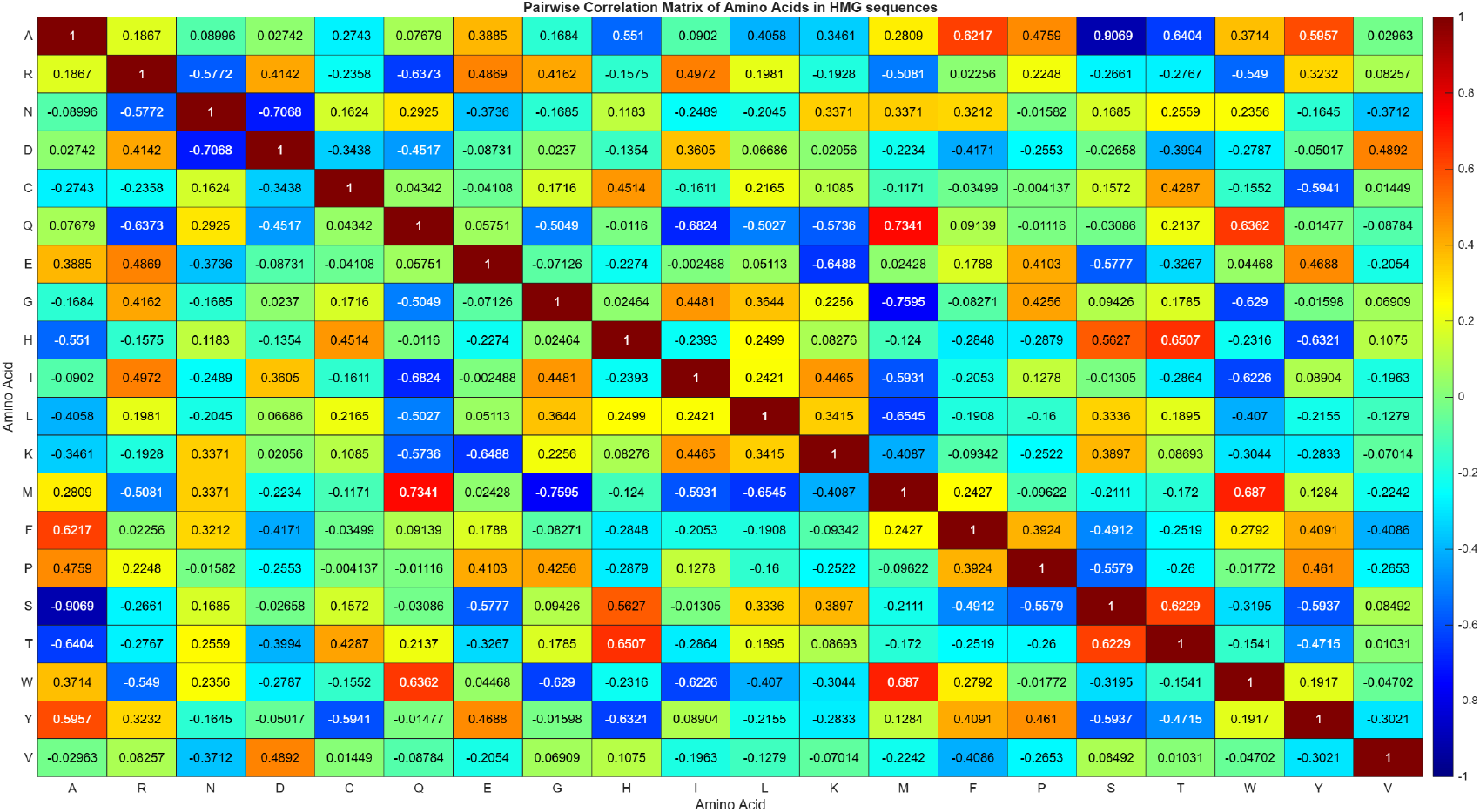
Heatmap of pairwise Pearson correlation coefficients among amino acid frequencies in 38 SRY-HMG subsequences. The analysis reveals patterns of co-occurrence and inverse association among amino acids within the highly conserved HMG domain of SRY proteins across different organisms.

Given that the HMG box constitutes the DNA-binding domain responsible for SRY-mediated activation of the testis-determining pathway, the observed amino acid co-variation patterns likely reflect evolutionary constraints acting on domain architecture rather than random sequence divergence [47]. The clustering of amino acid residues with similar physicochemical characteristics and the reciprocal variation of others may contribute to preserving the structural features required for DNA recognition and bending, both of which are essential for SRY function during male sex determination [6].

A dendrogram depicting the relationships among the 38 SRY-HMG subsequences was generated using their amino acid frequency composition profiles (Figure 6). At a distance threshold of 0.8, twenty-two subsequences were classified into seven distinct clusters, whereas the remaining sixteen SRY-HMG subsequences remained unclustered and were considered outliers. The identified clusters were distinguished by different colors in the dendrogram (Figure 6), and their compositions are presented in Table 7.

**Figure 6:**
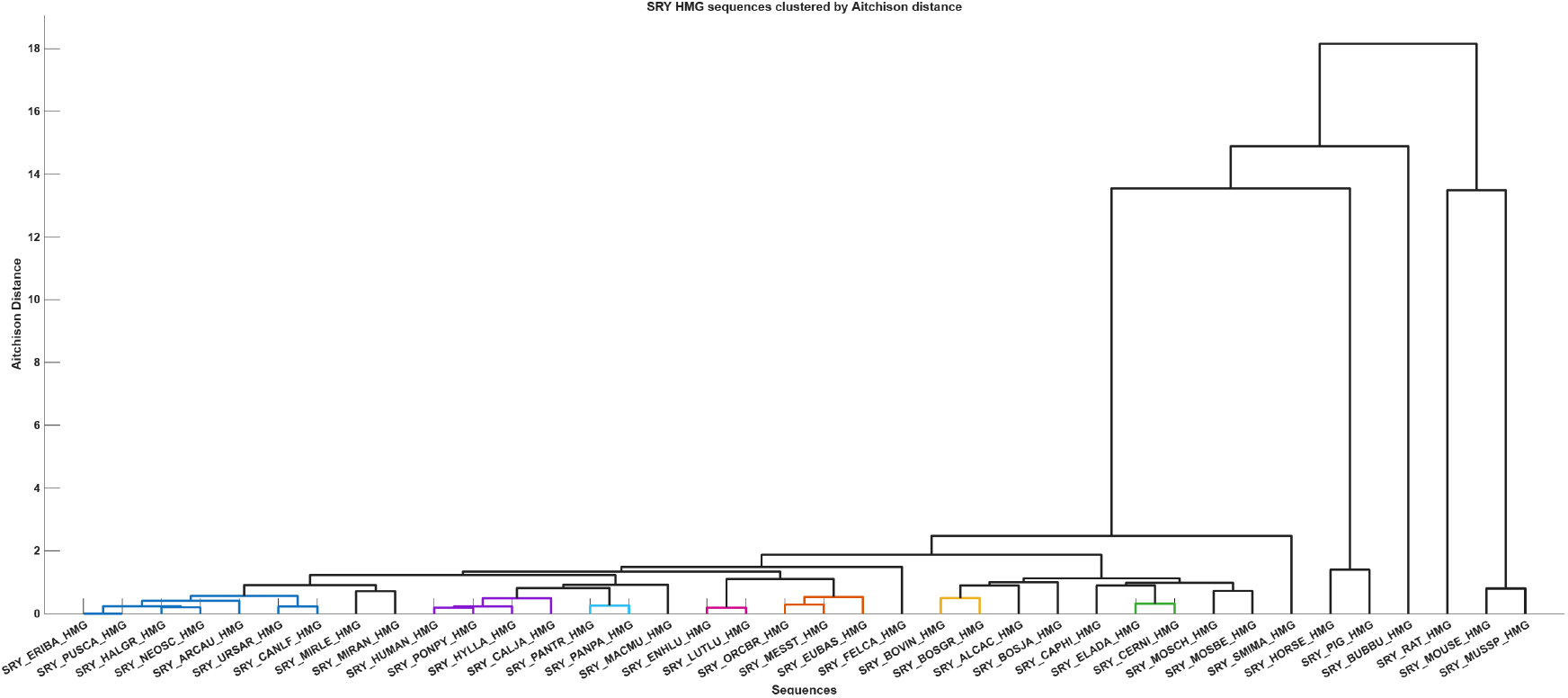
Dendrogram depicting the relationships among 38 SRY-HMG subsequences based on their amino acid frequency composition profiles using the Aitchison distance metric. Distinct clusters identified at a distance threshold of 0.8 are highlighted by different colors, whereas unclustered subsequences are shown in black.

**Table 7:** List of proximal clusters containing 22 SRY-HMG subsequences based on proximity derived from amino acid frequency profiles of 38 SRY sequences.

| Cluster | Sets of SRY-HMG subsequences |
| --- | --- |
| Cluster-1 | {SRY_ERIBA_HMG, SRY_PUSCA_HMG, SRY_HALGR_HMG, SRY_NEOSC_HMG, SRY_ARCAU_HMG, SRY_URSAR_HMG, SRY_CANLF_HMG} |
| Cluster-2 | {SRY_HUMAN_HMG, SRY_PONPY_HMG, SRY_HYLLA_HMG, SRY_CALJA_HMG} |
| Cluster-3 | {SRY_PANTR_HMG, SRY_PANPA_HMG} |
| Cluster-4 | {SRY_ENHLU_HMG, SRY_LUTLU_HMG} |
| Cluster-5 | {SRY_ORCBR_HMG, SRY_MESST_HMG, SRY_EUBAS_HMG} |
| Cluster-6 | {SRY_BOVIN_HMG, SRY_BOSGR_HMG} |
| Cluster-7 | {SRY_ELADA_HMG, SRY_CERNI_HMG} |

#### 4.1.3. Amino acid frequency based Shannon entropy analysis of SRY and SRY-HMG sequences

Shannon entropy analysis revealed differences in sequence complexity between the full-length SRY proteins and their corresponding HMG domains (Figure 7). Among the 65 SRY sequences studied, entropy values ranged from 3.4272 to 4.1901. The highest entropy was observed in SRY_SMIMA (4.1901), followed closely by SRY_ENHLU (4.1844) and SRY_MOSCH (4.1827), indicating a high degree of amino acid frequency diversity. In addition, SRY sequences from rodent such as SRY_MOUSE (3.4272), SRY_MUSSP (3.5515), and SRY_MUSSI (3.6958) showed notably lower entropy values, reflecting reduced sequence complexity relative to other taxa. The 38 SRY-HMG domains displayed a narrower entropy range, from 3.8965 to 4.1063, compared to the full-length SRY protein sequences. The highest entropy within the SRY-HMG subsequences was recorded for SRY_PIG_HMG (4.1063), while SRY_BUBBU_HMG (3.8965) had the lowest value. Most SRY-HMG subsequences had entropy values around 4.0, indicating relatively uniform amino acid diversity across organisms. Comparative analysis confirmed that full-length SRY proteins exhibit greater variability in sequence complexity than their SRY-HMG domains. This broader entropy range among full-length SRY sequences reflects increased diversification outside the SRY-HMGs, whereas the narrower range found in SRY-HMG domains highlights stronger conservation of this DNA-binding region. Despite these differences, both sequences maintained consistently high entropy values, underscoring substantial amino acid diversity across mammalian SRY proteins and their SRY-HMG domains [23].

**Figure 7:**
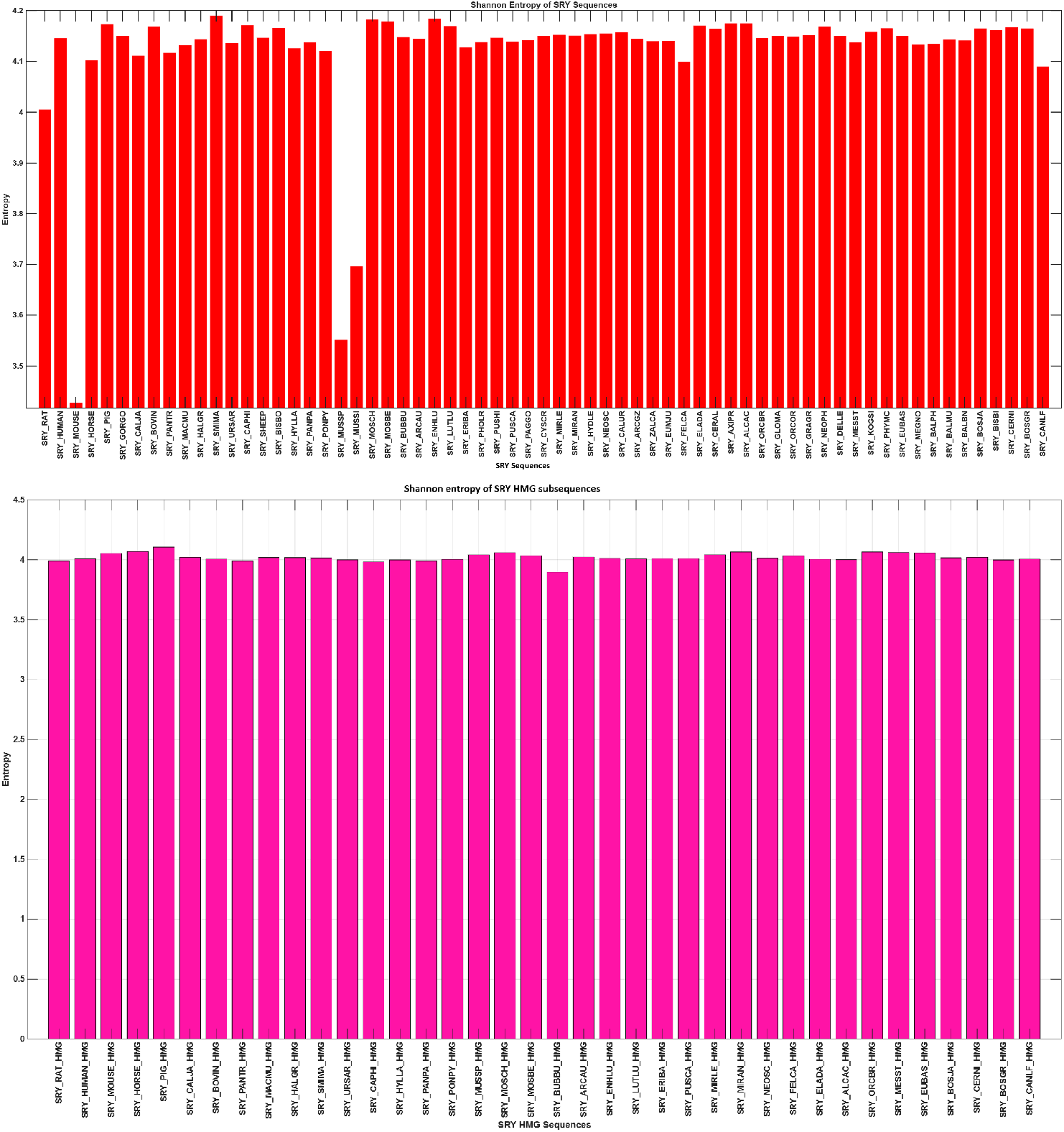
Shannon entropy profiles of full-length SRY proteins (top) and SRY-HMG domains (bottom) across mammalian organisms. Full-length SRY sequences exhibited a broader entropy distribution (3.4272–4.1901), with notably lower entropy in several rodent sequences, whereas SRY-HMG domains showed a narrower range of variation (3.8965–4.1063). The reduced variability of the HMG domains indicates greater compositional conservation compared with the full-length SRY proteins.

### 4.2. Analysis of frequency-dominant amino acid distribution over SRY and SRY-HMG sequences

Analysis of dominant amino acid profiles across the 65 SRY protein sequences from their respective organisms revealed substantial compositional diversity. The SRY sequences were distributed among 49 classes, with most clusters containing only a small number of organisms. The largest cluster comprised five bovine sequences, namely SRY_BOVIN, SRY_BISBO, SRY_BOSJA, SRY_BISBI, and SRY_BOSGR, all sharing the dominant amino acid profile ‘V-S-E-R-K-R-R-H-S-S-A-Y’ (Table 8). Several additional clusters were formed by closely related taxa, whereas many organisms possessed unique dominant amino acid patterns.

**Table 8:**
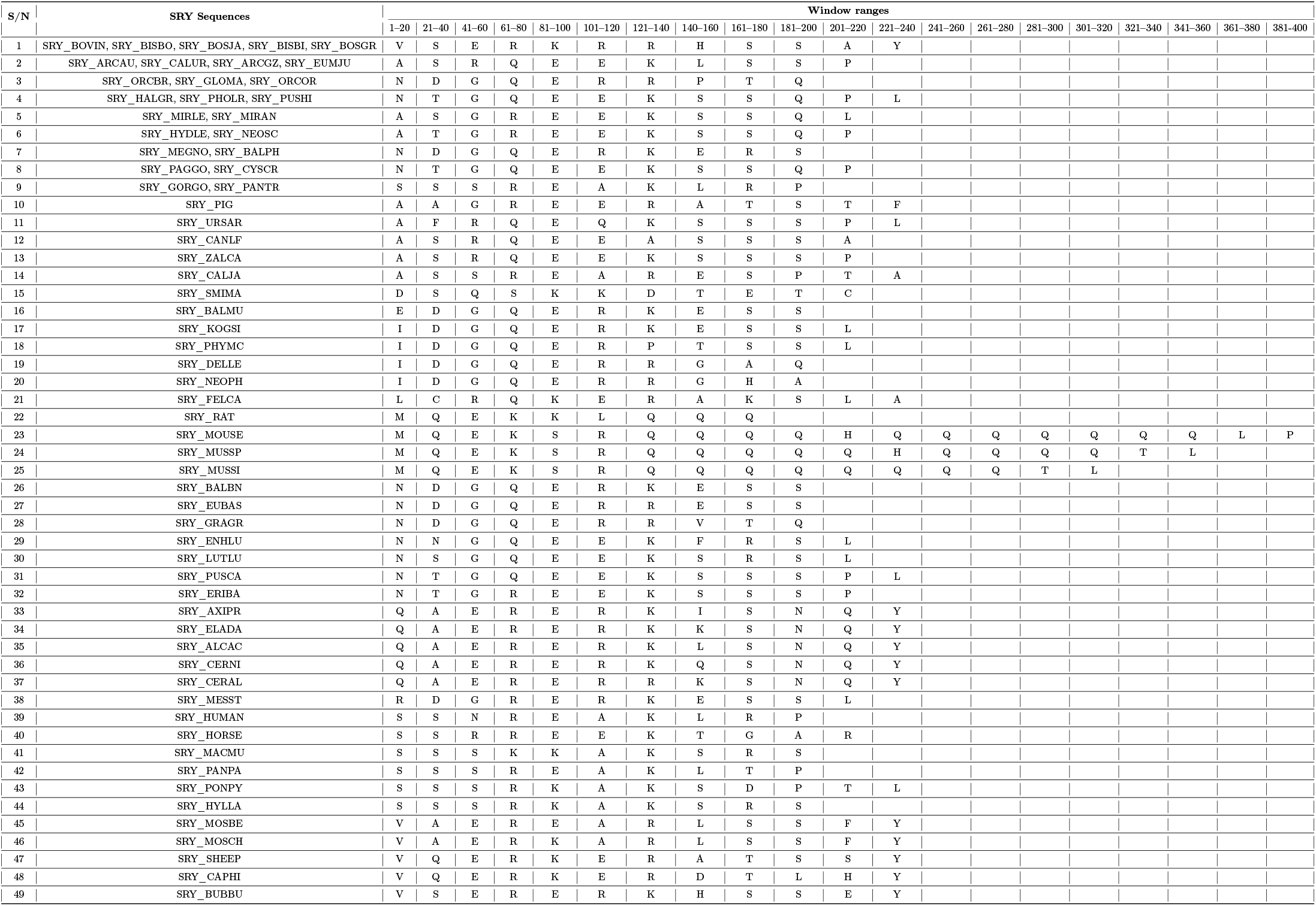
Frequency dominant amino acid profiles of 65 SRY protein sequences based on successive 20-residue windows. SRY sequences sharing identical profiles were grouped together.

Despite this diversity, recurring compositional features were evident across the set of SRY sequences from various organisms. Glutamic acid, arginine, lysine, serine, and glutamine repeatedly appeared as dominant residues in the central regions of the proteins, indicating the presence of a conserved compositional framework. Furthermore, the N-terminal and C-terminal regions exhibited considerably greater variation, contributing substantially to interorganisms differences in dominant amino acid profiles. A particularly distinctive pattern was observed in murine SRY proteins, especially SRY_MOUSE, SRY_MUSSP, and SRY_MUSSI, which displayed extended stretches of glutamine-dominated windows not observed in most other mammals (Table 8).

Based on the dominant amino acid present within successive 10-residue windows, the 38 SRY-HMG subsequences were partitioned into eighteen compositional clusters, indicating a high degree of overall conservation accompanied by measurable sequence-specific variation (Table 9). The largest cluster comprised nine sequences, all sharing an identical dominant amino acid profile (‘M-R-E-K-E-A-K’) (Table 9). Despite this conservation, most SRY-HMG subsequences exhibited unique dominant amino acid patterns, resulting in numerous small clusters and singleton groups, reflecting subtle lineage-specific divergence within an otherwise conserved domain.

**Table 9:** Frequency dominant amino acid profiles of 38 unique SRY-HMG subsequences based on successive 10-residue windows. SRY-HMG subsequences sharing identical profiles were grouped together.

| S/N | SRY-HMG subsequences | Window ranges |  |  |  |  |  |  |
| --- | --- | --- | --- | --- | --- | --- | --- | --- |
|  |  | 1-10 | 11-20 | 21-30 | 31-40 | 41-50 | 51-60 | 61-70 |
| 1 | SRY_HORSE_HMG, SRY_HALGR_HMG, SRY_ARCAU_HMG, SRY_ENHLU_HMG, SRY_LUTLU_HMG, SRY_ERIBA_HMG, SRY_MIRLE_HMG, SRY_MIRAN_HMG, SRY_NEOSC_HMG | M | R | E | K | E | A | K |
| 2 | SRY_HUMAN_HMG, SRY_CALJA_HMG, SRY_HYLLA_HMG, SRY_PONPY_HMG | V | R | E | K | E | Q | K |
| 3 | SRY_CAPHI_HMG, SRY_MOSCH_HMG, SRY_ELADA_HMG, SRY_CERNI_HMG | V | R | E | K | R | A | K |
| 4 | SRY_RAT_HMG, SRY_MOUSE_HMG, SRY_MUSSP_HMG | M | R | Q | K | E | L | K |
| 5 | SRY_PIG_HMG, SRY_ORCBR_HMG, SRY_EUBAS_HMG | V | R | E | K | E | A | K |
| 6 | SRY_URSAR_HMG, SRY_PUSCA_HMG | M | R | E | Q | E | A | K |
| 7 | SRY_BOVIN_HMG, SRY_BOSGR_HMG | V | R | K | K | R | A | K |
| 8 | SRY_PANPA_HMG | F | R | E | K | A | Q | K |
| 9 | SRY_PANTR_HMG | F | R | E | K | E | Q | K |
| 10 | SRY_MOSBE_HMG | I | R | E | K | R | A | K |
| 11 | SRY_BUBBU_HMG | I | R | E | K | R | E | K |
| 12 | SRY_SMIMA_HMG | M | Q | N | K | L | D | K |
| 13 | SRY_FELCA_HMG | M | R | E | K | A | A | R |
| 14 | SRY_CANLF_HMG | R | R | E | K | E | A | K |
| 15 | SRY_MACMU_HMG | V | K | E | K | A | Q | K |
| 16 | SRY_MESST_HMG | V | R | E | K | E | A | Y |
| 17 | SRY_ALCAC_HMG | V | R | E | K | R | A | D |
| 18 | SRY_BOSJA_HMG | V | R | K | K | R | A | H |

Across the clusters, the 11–20, 31–40, and 61–70 windows were highly conserved, being predominantly occupied by arginine, lysine, and lysine, respectively (Table 9). The 21–30 and 41–50 windows showed moderate variability, but remained largely enriched in glutamic acid and either glutamic acid or arginine. Greater variation was observed in the 1– 10 and 51–60 windows, where dominant residues included valine, methionine, phenylalanine, isoleucine, arginine, alanine, glutamine, leucine, glutamic acid, and aspartic acid (Table 9). The persistent enrichment of arginine and lysine across multiple windows underscores the strong conservation of positively charged residues associated with sequence-specific DNA binding and bending by the HMG-box domain. Thus, while the overall compositional architecture of the SRY-HMG domain remains highly conserved across mammals, the presence of numerous cluster-specific dominant residue profiles suggests limited evolutionary diversification that preserves the fundamental structural and functional requirements of the domain [48, 10].

These frequency-dominant amino acid analyses suggested that evolutionary diversification of SRY protein sequences from 65 organisms had occurred predominantly outside the HMG-box, while the dominant amino acid composition of the SRY-HMG domain had remained comparatively constrained across organisms, consistent with its conserved functional role.

Figure 8 illustrated the dominant amino acid profiles of SRY_HUMAN and its corresponding HMG-box domain (SRY_HUMAN_HMG). While the full-length SRY protein exhibited variation in the dominant residues across successive windows (‘S-S-N-R-E-A-K-L-R-P’), the respective HMG-box domain displayed a more compact profile (‘V-R-E-K-E-Q-K’) characterized by recurrent enrichment of arginine and lysine. These representative profiles exemplified the greater compositional complexity of the full-length SRY protein relative to the more conserved HMG-box region. Compared with the comparatively diverse dominant profile of SRY_HUMAN, the SRY_HUMAN_HMG domain exhibited a shorter and more conserved profile, consistent with the overall pattern of stronger conservation observed among SRY-HMG sequences than among full-length SRY protein sequences.

**Figure 8:**
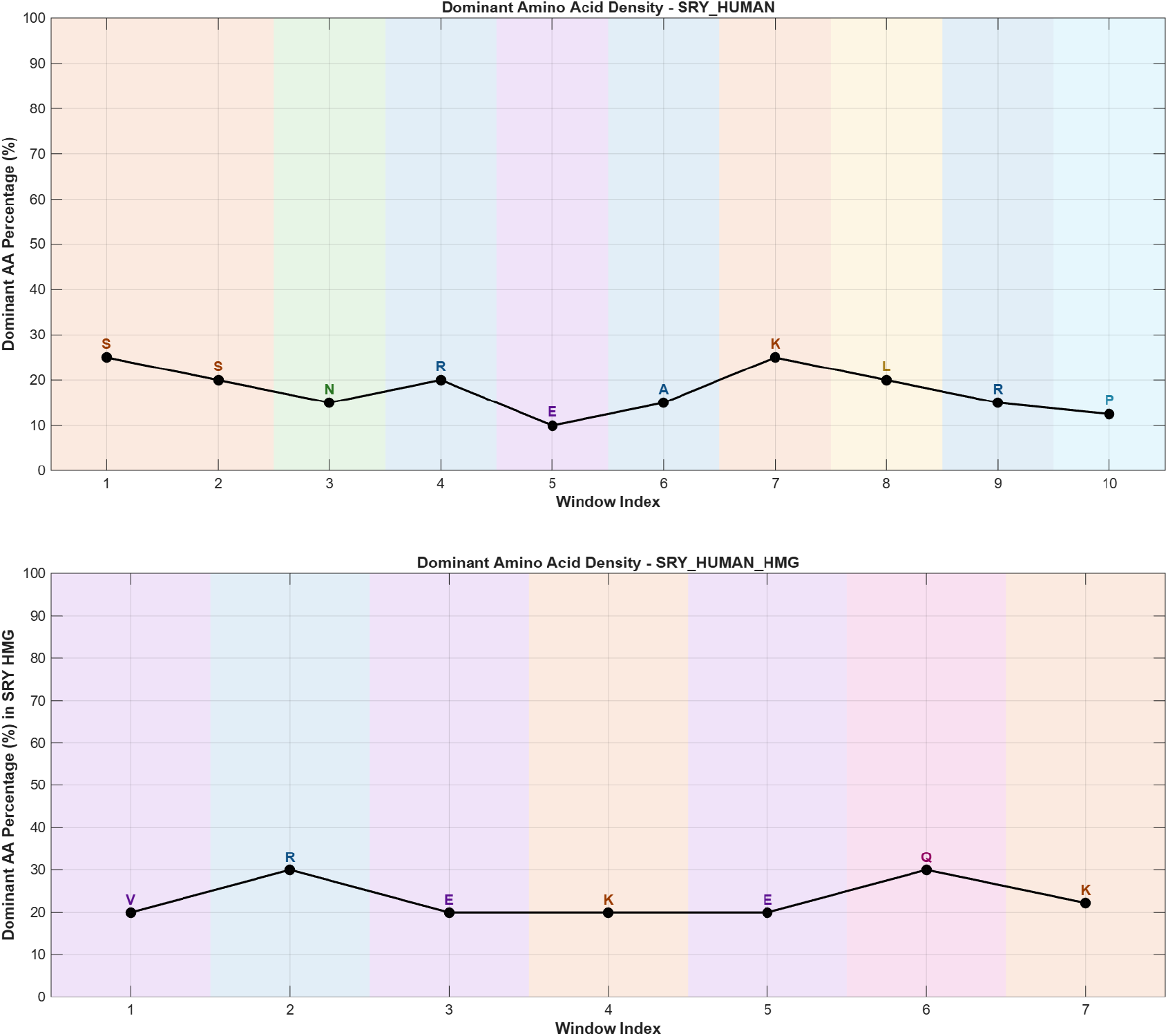
Dominant amino acid profile of SRY_HUMAN based on successive 20-residue windows. The dominant profile was S-S-N-R-E-A-K-L-R-P. Dominant amino acid profile of the SRY_HUMAN_HMG domain based on successive 10-residue windows. The dominant profile was V-R-E-K-E-Q-K.

### 4.3. Amino acid sequence homology-based phylogenetic analysis of SRY and SRY-HMG sequences

#### 4.3.1. Amino acid sequence homology-based phylogenetic analysis of SRY sequences

Positional evolutionary pressure and mutational tolerance at single-residue resolution across the full-length protein were quantified by mapping a position-specific Shannon entropy profile across SRY sequences. The computed entropy values spanned a dynamic range from an absolute floor of 0.00 bits (total invariance) to a maximum peak of 2.83 bits (extreme hyper-variability) (Figure 9). The structural landscape revealed a highly polarized, bipartite architecture: a highly variable, structurally plastic N-terminal region followed by a broad, rigidly conserved functional domain. Specifically, the first half of the alignment exhibited severe architectural volatility (0.50–2.81 bits) with prominent mutational spikes (>2.00 bits) clustered at coordinates 30–60, 140–180, and 190–240, reaching a global maximum of 2.81 bits at position 157. This extensive variation reflected a severe relaxation of purifying selection, corresponding biologically to poorly structured N-terminal flanking loops and highly divergent, lineage-specific insertions.

**Figure 9:**
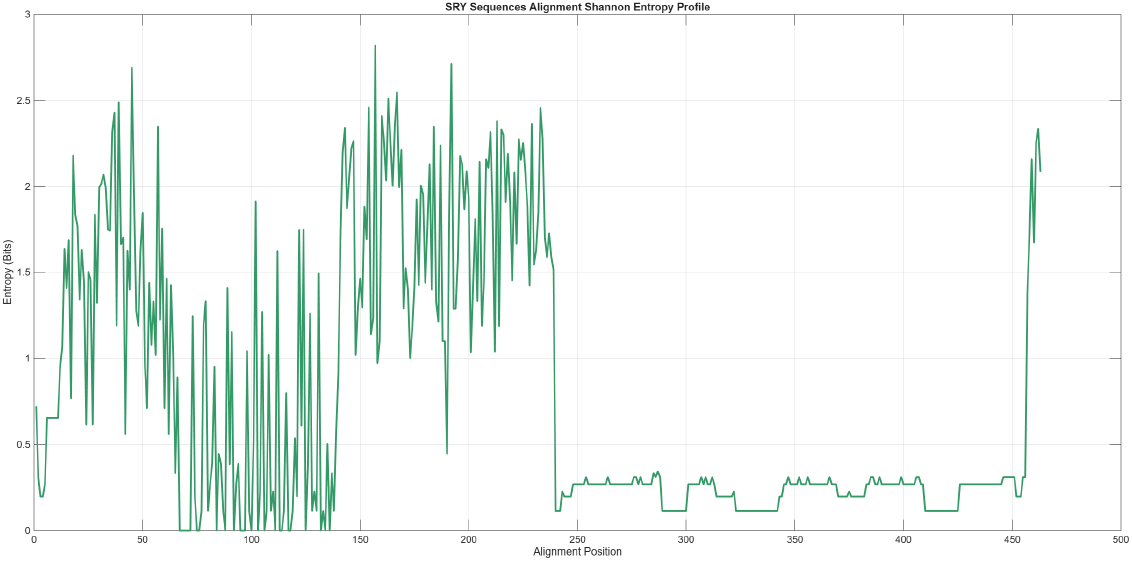
Position-specific Shannon entropy profile of the SRY sequences across 65 organisms.

**Figure 10:**
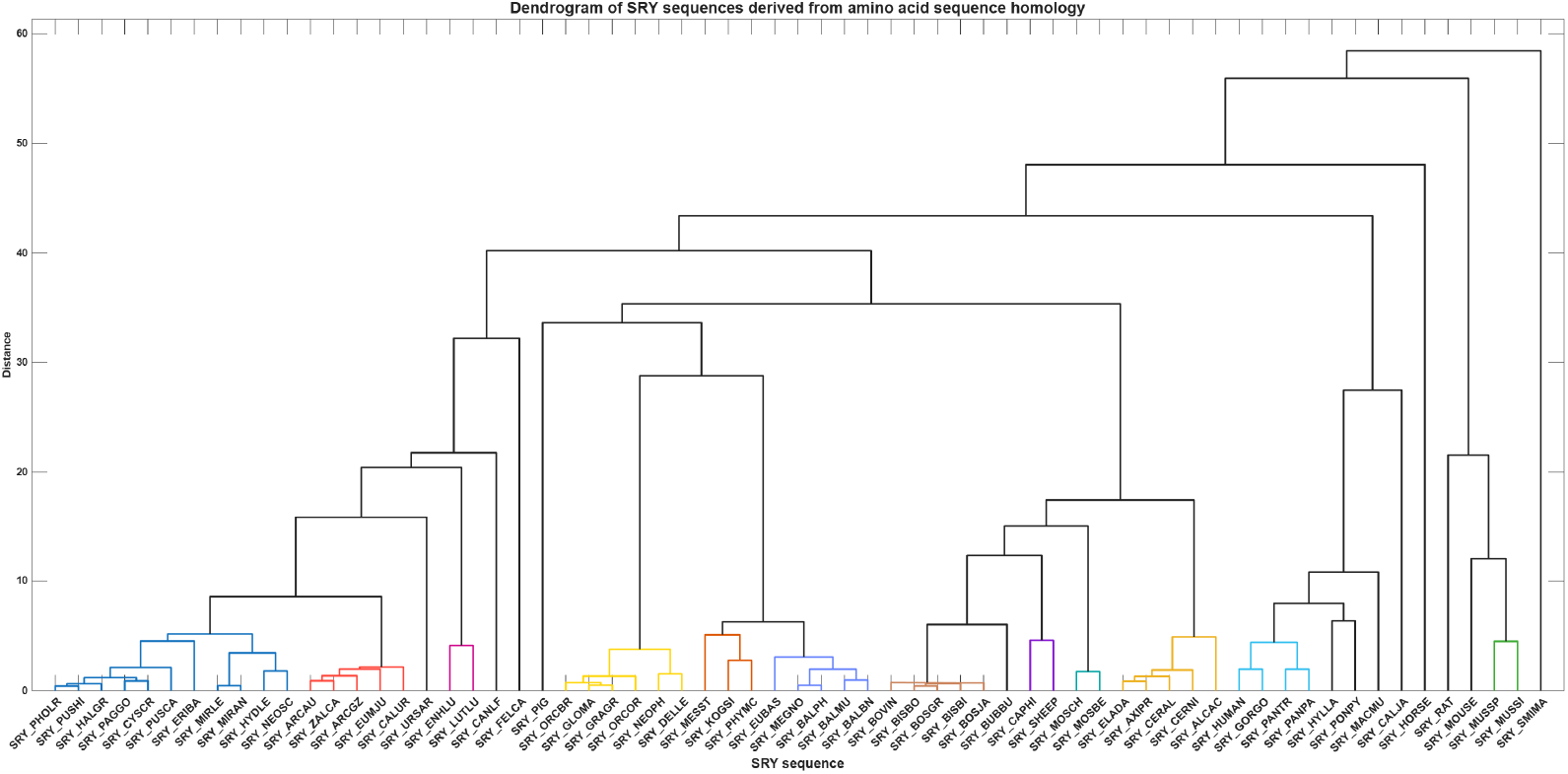
Dendrogram showing homology relationships among 65 SRY sequences from 65 organisms based on amino acid sequence similarity.

It was observed that a transition occurred near alignment coordinate 240, where the entropy profile dropped into an exceptional, prolonged conservation plateau that maintained a tight ceiling between 0.15 and 0.35 bits up to position 455. Within this region, absolute zero-entropy valleys (H = 0.00 bits) were tightly concentrated around positions 242, 288–298, 320–330, 342, and 410–420. This massive, unbroken 215-residue compression provided direct quantitative evidence of intense purifying selection locking the critical functional infrastructure of the SRY transcription factor. Biologically, these invariant sites governed internal hydrophobic core packing and crucial DNA minor-groove binding/intercalation. Finally, a sharp terminal spike up to 2.35 bits at position 462 marked the flexible, lineage-specific C-terminal tail, confirming that while flanking domains accommodated high evolutionary diversity, the spatial configuration required for DNA architectural bending remained under absolute negative selection across all 38 organisms. Consequently, non-synonymous changes within these ultra-conserved valleys directly disrupted DNA bending and transcriptional activation, directly explaining the molecular etiology of clinical phenotypes such as Swyer syndrome (46,XY gonadal dysgenesis) [49, 50].

A total of 65 SRY amino acid sequences were subjected to hierarchical clustering based on sequence homology, and the resulting dendrogram was cut at a distance threshold of 5 to define clusters (Figure 12). This yielded 12 distinct clusters (Cluster-1 to Cluster-12), varying in size from 2 to 11 sequences (Table 10). Cluster-1 was the largest (11 sequences), while Clusters 3, 8, 9, and 12 were the smallest (2 sequences each). The clusters showed clear taxonomic congruence, grouping the Bovini (Cluster-7), Caprini (Cluster-8), Moschidae (Cluster-9), Cervidae (Cluster-10), Hominini (Cluster-11), Mus (Cluster-12), and Balaenopteridae (Cluster-6) (Table 10).

**Figure 11:**
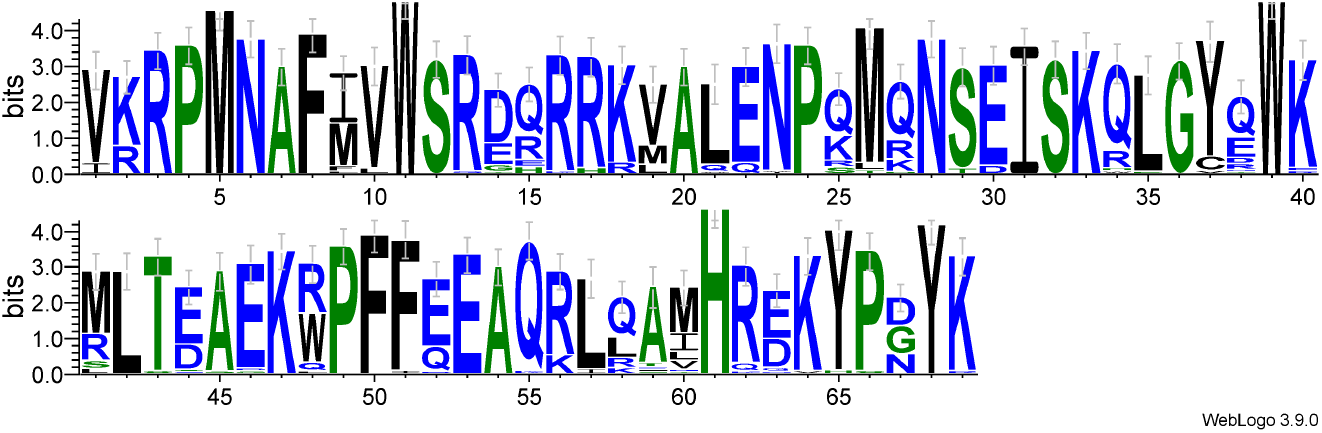
Sequence-logo showing positional conservation and amino-acid composition of the SRY-HMG subsequences.

**Figure 12:**
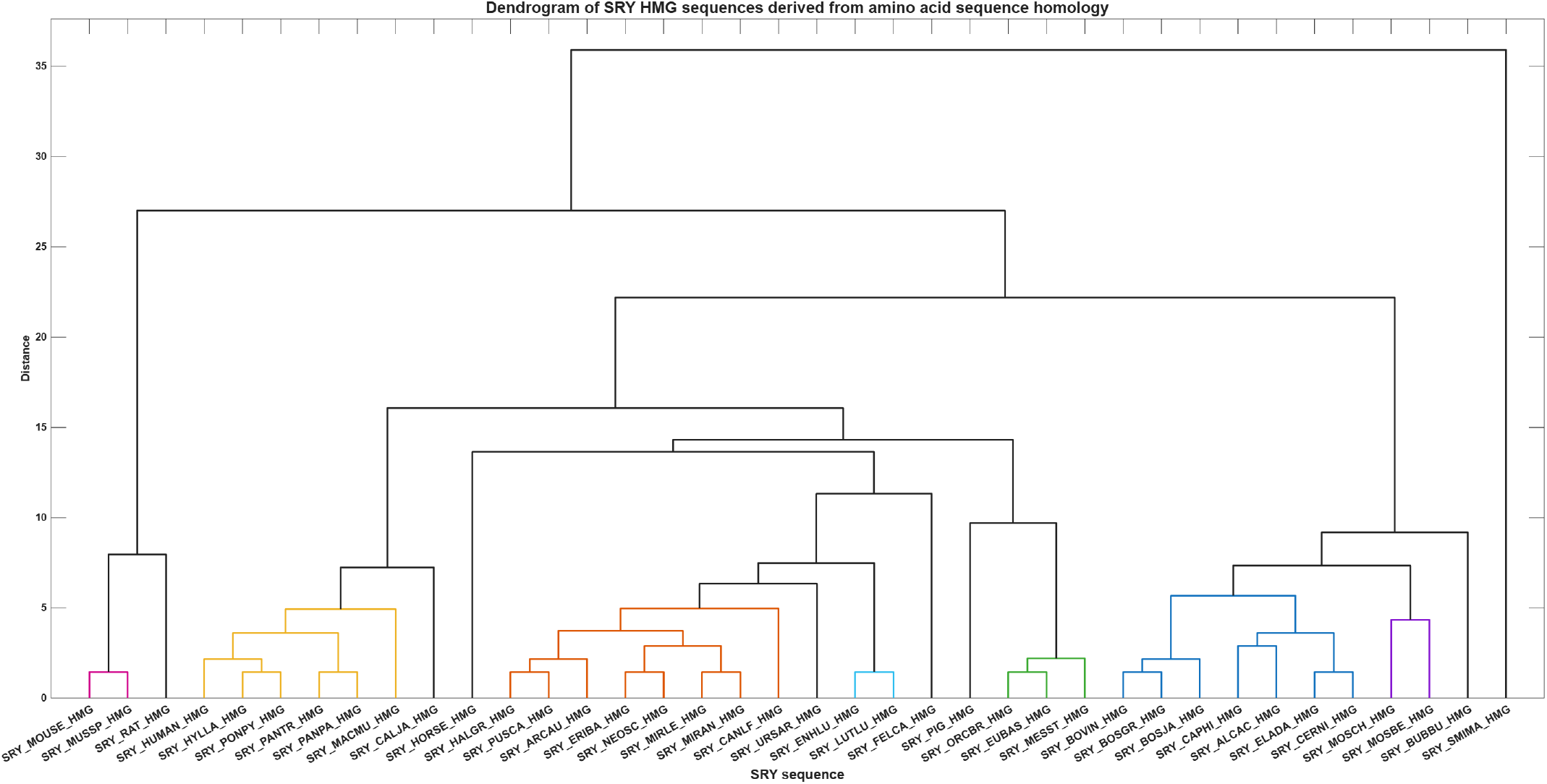
Dendrogram showing homology relationships among 38 SRY-HMG subsequences based on amino-acid sequence similarity.

**Table 10:**
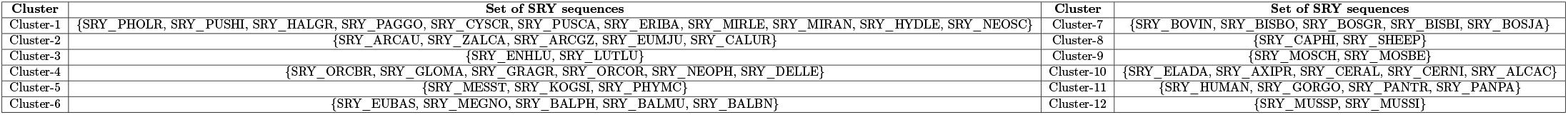
List of cluster containing proximal SRY sequences from various organisms derived from amino acid sequence homology by setting a distance cutoff of 5.

#### 4.3.2. Amino acid sequence homology-based phylogenetic analysis of SRY-HMG subsequences

Multiple sequence alignment of the HMG-box domain of SRY orthologs from a broad range of mammalian organisms was performed, and sequence conservation was visualized with WebLogo (Figure 11). The aligned region comprised 69 amino-acid positions and exhibited a high overall degree of sequence identity. It was found that the total number of strictly invariant residues (identical amino acid in every sequence) was 25. In the sequence-logo these sites appeared as single letters of maximal information content (4 bits), confirming absolute conservation across the examined taxa (Figure 11). At position 1 of the alignment, isoleucine was observed exclusively in four SRY-HMG sequences (SRY_MOSBE_HMG, SRY_BUBBU_HMG, SRY_ENHLU_HMG and SRY_LUTLU_HMG), whereas all remaining SRY-HMGs contained valine at this site. Both isoleucine and valine were hydrophobic amino acids; consequently, this substitution was chemically conservative and was not expected to alter the overall physicochemical properties of the domain. Likewise, at position 2 of the alignment the majority of SRY-HMG sequences contained lysine, whereas eleven orthologs (SRY_HALGR_HMG, SRY_URSAR_HMG, SRY_ARCAU_HMG, SRY_ENHLU_HMG, SRY_LUTLU_HMG, SRY_ERIBA_HMG, SRY_PUSCA_HMG, SRY_MIRLE_HMG, SRY_MIRAN_HMG, SRY_NEOSC_HMG and SRY_CANLF_HMG) possessed arginine. Both lysine and arginine were positively charged basic residues; the substitution was therefore chemically conservative and preserved the electrostatic character of this site [51].

In the similar fashion, ten additional positions displayed only chemically conservative substitutions belonging to the same physicochemical class (Table 11). The majority of variable sites tolerated only conservative changes (hydrophobic-for-hydrophobic or polar-for-polar), while a smaller subset showed more pronounced amino-acid diversity. These variable positions mapped predominantly to solvent-exposed loops and the extreme termini of the domain and were reflected in the logo as multi-letter stacks of reduced height (Figure 11). The resulting conservation pattern was therefore highly non-random: the 25 invariant residues concentrated on the DNA-binding surface and the hydrophobic core required to maintain the characteristic L-shaped HMG-box architecture, whereas variable residues occupied peripheral locations that were under weaker selective constraint [52].

**Table 11:** Positions (sequence-logo in Figure 11) in the SRY HMG-box alignment that exhibit only physicochemically conservative substitutions. For each position the observed amino acids, their shared physicochemical class, and the nature of the substitution are indicated.

| Position | Residues changes | Physicochemical class | Nature of substitution |
| --- | --- | --- | --- |
| 9 | M / I / V / T / F | Predominantly hydrophobic | Conservative |
| 10 | V / M / L / F / I | Hydrophobic | Fully conservative |
| 12 | S / R / G / D / H / Q / T / N | Polar/charged | Moderately conservative |
| 14 | D / E / R / Q / H / G | Charged/ polar | Conservative |
| 15 | Q / R / H / E / G | Polar/ charged | Conservative |
| 20 | V / M / A / R / L / Q | Predominantly hydrophobic | Conservative |
| 21 | A / V / M / L / Q / R | Predominantly hydrophobic | Conservative |
| 27 | Q / K / R / S / T / L / D | Polar/charged | Conservative |
| 42 | M / R / S / L | Hydrophobic/ polar | Limited conservative |
| 59 | Q / K / R / L / A / V / M | Polar/charged/ hydrophobic | Conservative (volume/charge) |
| 60 | A / M / T / L / V / I | Predominantly hydrophobic | Conservative |
| 67 | D / N / G / H / R / Q / S | Polar/charged | Conservative |

An amino acid sequence similarity-based analysis was performed to establish the relationships among the 38 distinct SRY-HMG subsequences, resulted in the dendrogram shown in Figure 12. Using a distance threshold of 5.0, the 30 subsequences were grouped into seven clusters of proximal sequences. Table 12 summarized the proximal SRY-HMG subsequences identified across the different organisms. Cluster-3 contained the largest group (8 subsequences), followed by Cluster-6 (7) and Cluster-2 (6), whereas Clusters-1, 4, and 7 each contained 2 subsequences and Cluster-5 contained 3. This clustering pattern indicated distinct groups of closely related SRY-HMG amino acid subsequences among the organisms.

**Table 12:** List of proximal clusters containing 30 SRY-HMG subsequences based on amino acid sequence homology of 38 SRY sequences.

| Cluster | Sets of SRY-HMG subsequences |
| --- | --- |
| Cluster-1 | {SRY_MOUSE_HMG, SRY_MUSSP_HMG} |
| Cluster-2 | {SRY_HUMAN_HMG, SRY_HYLLA_HMG, SRY_PONPY_HMG, SRY_PANTR_HMG, SRY_PANPA_HMG, SRY_MACMU_HMG} |
| Cluster-3 | {SRY_HALGR_HMG, SRY_PUSCA_HMG, SRY_ARCAU_HMG, SRY_ERIBA_HMG, SRY_NEOSC_HMG, SRY_MIRLE_HMG, SRY_MIRAN_HMG, SRY_CANLF_HMG} |
| Cluster-4 | {SRY_ENHLU_HMG, SRY_LUTLU_HMG} |
| Cluster-5 | {SRY_ORCBR_HMG, SRY_EUBAS_HMG, SRY_MESST_HMG} |
| Cluster-6 | {SRY_BOVIN_HMG, SRY_BOSGR_HMG, SRY_BOSJA_HMG, SRY_CAPHI_HMG, SRY_ALCAC_HMG, SRY_ELADA_HMG, SRY_CERNI_HMG} |
| Cluster-7 | {SRY_MOSCH_HMG, SRY_MOSBE_HMG} |

### 4.4. Clinically significant variants of SRY_HUMAN

A total of 49 distinct sequence alterations in SRY_HUMAN were compiled from Ensembl, ClinVar, and UniProt annotations. Of these, 44 were classified as pathogenic and 5 as likely pathogenic. The variants comprised 33 missense changes, 12 nonsense (STOP) mutations, and 4 deletions (denoted as MISSING in the source data) (Table 13).

**Table 13:** Clinically significant variants of human SRY from Ensembl, ClinVar, and UniProt.

| Change | Clinical Significance | Change | Clinical Significance | Change | Clinical Significance |
| --- | --- | --- | --- | --- | --- |
| Q2STOP | Pathogenic (Ensembl, ClinVar) | S71Y | Likely pathogenic (Ensembl, ClinVar) | L101H | Pathogenic (UniProt) |
| S3L | Pathogenic (UniProt) | R75M | Pathogenic (UniProt) | T102I | Likely pathogenic (ClinVar) |
| Y4STOP | Pathogenic (Ensembl, ClinVar) | R76L | Likely pathogenic (Ensembl, ClinVar) | K106I | Pathogenic (Ensembl, ClinVar, UniProt) |
| S18N | Pathogenic (Ensembl, ClinVar) | R76S | Pathogenic (UniProt) | W107STOP | Pathogenic (Ensembl, ClinVar) |
| R30I | Pathogenic (Ensembl, ClinVar) | M78T | Pathogenic (UniProt) | P108R | Pathogenic (UniProt) |
| G49STOP | Pathogenic (Ensembl, ClinVar) | N87Y | Pathogenic (UniProt) | F109MISSING | Pathogenic (ClinVar) |
| Q57STOP | Pathogenic (Ensembl, ClinVar) | S88STOP | Pathogenic (Ensembl, ClinVar) | F109S | Pathogenic (Ensembl, ClinVar, UniProt) |
| R59MISSING | Pathogenic (ClinVar) | E89MISSING | Pathogenic (ClinVar) | Q111STOP | Pathogenic (Ensembl, ClinVar) |
| V60A | Pathogenic (Ensembl, ClinVar) | I90M | Pathogenic (Ensembl, ClinVar, UniProt) | A113T | Pathogenic (Ensembl, ClinVar, UniProt) |
| V60L | Pathogenic (Ensembl, ClinVar, UniProt) | S91T | Pathogenic (UniProt) | E122MISSING | Pathogenic (ClinVar) |
| R62G | Pathogenic (UniProt) | S91G | Likely pathogenic (ClinVar) | P125L | Pathogenic (UniProt) |
| M64I | Pathogenic (UniProt) | K92STOP | Pathogenic (Ensembl, ClinVar) | Y127C | Pathogenic (Ensembl, ClinVar, UniProt) |
| M64R | Pathogenic (UniProt) | Q93STOP | Pathogenic (Ensembl, ClinVar) | Y127F | Pathogenic (Ensembl, ClinVar, UniProt) |
| A66P | Likely pathogenic (Ensembl, ClinVar) | G95E | Pathogenic (Ensembl, ClinVar, UniProt) | P131R | Pathogenic (UniProt) |
| F67V | Pathogenic (UniProt) | G95R | Pathogenic (Ensembl, ClinVar, UniProt) | R133W | Pathogenic (Ensembl, ClinVar, UniProt) |
| I68T | Pathogenic (Ensembl, ClinVar, UniProt) | Y96STOP | Pathogenic (Ensembl, UniProt) |  |  |
| W70STOP | Pathogenic (Ensembl, ClinVar) | W98STOP | Pathogenic (Ensembl, UniProt) |  |  |

The distribution of these alterations along the protein was markedly non-uniform. Only six variants mapped to the N-terminal region preceding the HMG box: Q2STOP, S3L, Y4STOP, S18N, R30I, and G49STOP. One nonsense mutation, Q57STOP, occurred at the boundary of the HMG box. The overwhelming majority—38 of the 49 variants—fell within the HMG box itself, spanning residues 59 to 125. This included all four deletions (R59MISSING, E89MISSING, F109MISSING, and E122MISSING), the remaining nonsense mutations (W70STOP, S88STOP, K92STOP, Q93STOP, Y96STOP, W98STOP, W107STOP, and Q111STOP), and numerous missense changes. Only four variants were located C-terminal to the HMG box: Y127C, Y127F, P131R, and R133W. Several positions appeared to be mutational hotspots, with more than one independent alteration reported. Codon 60 carried both V60A and V60L; codon 64 carried M64I and M64R; codon 76 carried R76L and R76S; codon 91 carried S91T and S91G; codon 95 carried G95E and G95R; codon 109 carried F109MISSING and F109S; and codon 127 carried Y127C and Y127F. Many of these recurrent positions correspond to residues that are highly conserved across species and are known to be critical for the structure or DNA-binding function of the HMG box [53].

Furthermore, a separate set of N-terminal domain (NTD) and C-terminal domain (CTD) mutations with associated proband phenotypes was examined (Table 14). In the NTD, excluding premature stop codons, four missense mutations were identified: S3L, S18N, R30I, and Q57R. These were associated with 46,XY pure or partial gonadal dysgenesis, and in one case with Turner syndrome with gonadal dysgenesis [54]. It was noted that S18N was observed in two probands with different karyotypes and phenotypes: one 46,XY female with partial gonadal dysgenesis and one 45, XO/46, XY female with Turner syndrome and gonadal dysgenesis [54]. R30I was reported in a proband described as 46,XY male or female with pure or partial gonadal dysgenesis. Q57R and S3L were each associated with pure gonadal dysgenesis in 46, XY females [54, 55]. In the CTD, five mutations were recorded: S143G, S143C, Q158fsX180, Q159fsX167, and L163X. These included two missense changes (S143G and S143C), two frameshift mutations (Q158fsX180 and Q159fsX167), and one nonsense mutation (L163X) [56]. All were associated with pure gonadal dysgenesis except Q159fsX167, which was found in a 45,XO/46,XY proband with Turner syndrome and gonadal dysgenesis [56].

**Table 14:** Phenotype-associated mutations in the NTD and CTD of human SRY.

| Region | Mutation | Proband | Phenotype |
| --- | --- | --- | --- |
| NTD | S3L | 46,XY female | Pure gonadal dysgenesis [57, 58, 59] |
| NTD | S18N | 46,XY female | Partial gonadal dysgenesis [57, 58, 59] |
| NTD | S18N | 45,XO/46,XY female | Turner syndrome with gonadal dysgenesis [57, 59, 60] |
| NTD | R30I | 46,XY male or female | Pure/partial gonadal dysgenesis [57, 58, 59] |
| NTD | Q57R | 46,XY female | Pure gonadal dysgenesis [57, 58, 59] |
| CTD | S143G | 46,XY female | Pure gonadal dysgenesis [57, 58, 59] |
| CTD | S143C | 46,XY female | Pure gonadal dysgenesis [57, 58, 59] |
| CTD | Q158fsX180 | 46,XY female | Pure gonadal dysgenesis [57, 58, 59] |
| CTD | Q159fsX167 | 45,XO/46,XY female | Turner syndrome with gonadal dysgenesis [57, 59, 60] |
| CTD | L163X | 46,XY female | Pure gonadal dysgenesis [57, 58, 59] |

Therefore, this information reinforced the functional importance of the HMG box as the primary target of clinically significant alterations in SRY_HUMAN, while also confirming that mutations in the NTD and CTD can impair gonadal development. The concentration of pathogenic changes within the HMG box was consistent with its essential role in sequence-specific DNA recognition. Truncating mutations and deletions within this domain would be expected to abolish DNA binding entirely, while missense changes at conserved positions might impair folding or reduce binding affinity. In addition, the smaller number of variants in the N- and C-terminal regions may affect other aspects of SRY function, including the disordered terminal segments that were previously identified as droplet-promoting and aggregation-prone in the LLPS analysis.

### 4.5. Liquid-liquid phase separation propensity of SRY_HUMAN and its SRY_HUMAN_HMG

The SRY_HUMAN sequence was analysed for its propensity to undergo spontaneous liquid-liquid phase separation using the FuzDrop web server. The predicted probability of phase separation was *p_LLP S_* = 0.5734, indicating a moderate-to-high propensity, although it fell just below the 0.60 threshold conventionally used to classify a protein as a droplet driver (Figure 13a). Two droplet-promoting regions (DPRs) were identified, located at the N-terminus (residues 1–23) and the C-terminus (residues 179–204). These regions corresponded to intrinsically disordered terminal segments. Two aggregation hot-spots were also predicted, at residues 15–23 and 187–199, both of which overlapped with the identified DPRs, suggesting that these disordered regions might also have driven irreversible maturation of the droplet state. Multiple regions with context-dependent binding modes (*S_bind_*) were identified, including residues 15–34, 61–66, 71–80, 95–105, 154–162, and 187–199.

**Figure 13:**
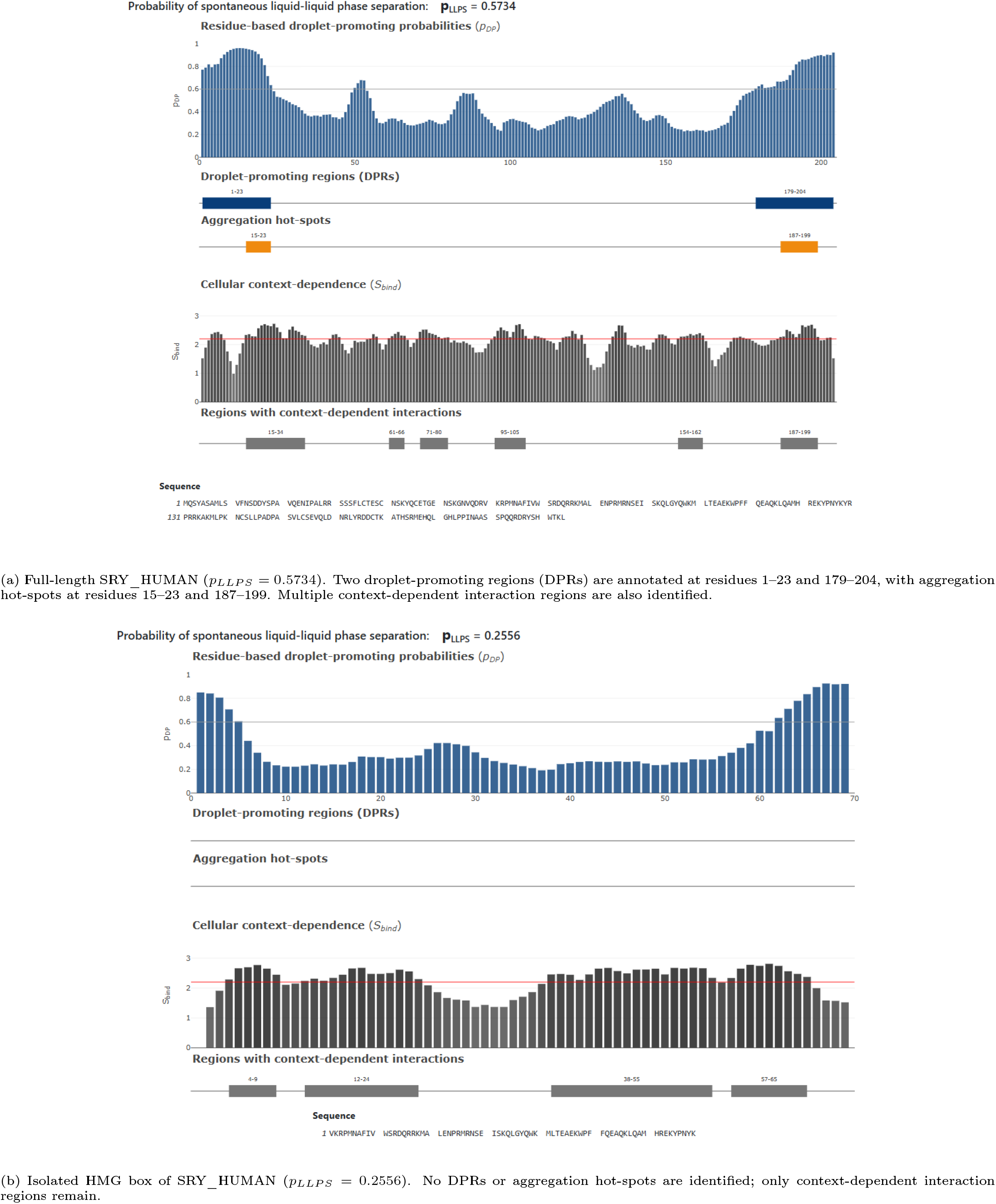
FuzDrop prediction of liquid-liquid phase separation (LLPS) propensity for SRY_HUMAN and its isolated HMG box. Top: full-length SRY_HUMAN, showing a moderate-to-high LLPS propensity driven by its disordered terminal regions. Bottom: the isolated HMG box, showing a markedly lower LLPS propensity with no droplet-promoting or aggregation-prone regions.

On the other side, the isolated HMG box domain of SRY_HUMAN (SRY_HUMAN_HMG) exhibited a much lower propensity for spontaneous phase separation, with *p_LLP S_* = 0.2556 (Figure 13b). No DPRs or aggregation hot-spots were identified in this domain. Four regions with context-dependent interactions were predicted (residues 4–9, 12–24, 38–55, and 57–65), which likely corresponded to the DNA-binding surface and protein interaction interfaces of the HMG box [61].

The human HMG box was a structured, conserved domain with low LLPS propensity, consistent with its role in sequence-specific DNA binding. The flanking N- and C-terminal regions, by contrast, were intrinsically disordered, contained the DPRs and aggregation hot-spots, and were responsible for the higher LLPS propensity of the full-length protein. This architectural arrangement was characteristic of transcription factors that formed biomolecular condensates at specific genomic loci. The marked difference between the two FuzDrop profiles (Figures 13a and 13b) visually confirmed that the phase separation behaviour of SRY_HUMAN was driven primarily by its terminal disordered regions rather than by its DNA-binding domain.

### 4.6. Dendrogram relationships among SRY and SRY-HMG sequences based on dipeptide composition analysis

#### 4.6.1. Prevalence of dipeptide frequency and associated dendrogram of SRY protein sequences

The 400 possible dipeptides were classified into five prevalence categories based on their occurrence across SRY protein sequences (Table 15). The distribution was markedly skewed toward low prevalence. The largest proportion of dipeptides fell into the rare category (*≤* 25%), followed by a substantial number that were completely absent from all SRY sequences, indicating that a large fraction of amino acid pair combinations does not occur in the SRY protein. In contrast, only a small subset was highly prevalent (>75%). When examined as amino acid couples, the highly prevalent dipeptides were dominated by combinations in which both partners are polar or charged, or in which a polar residue is paired with a small nonpolar residue - a pattern consistent with solvent-exposed and DNA-contacting surfaces. The common category (51-75%) followed the same trend, with polar–polar and polar–charged couples prevailing, while moderately prevalent dipeptides (26–50%) showed a more balanced mix of polar–nonpolar and nonpolar–nonpolar couplings. In addition, the rare and absent categories were enriched for couples in which both partners are bulky nonpolar residues, or that involve cysteine and other residues with limited compatibility in the SRY fold. Overall, the prevalence distribution revealed a clear compositional bias: polar- and charged-coupled dipeptides are widely conserved, whereas nonpolar–nonpolar and sterically demanding couples are largely excluded, consistent with strong purifying selection acting on the functional core of the protein.

**Table 15:**
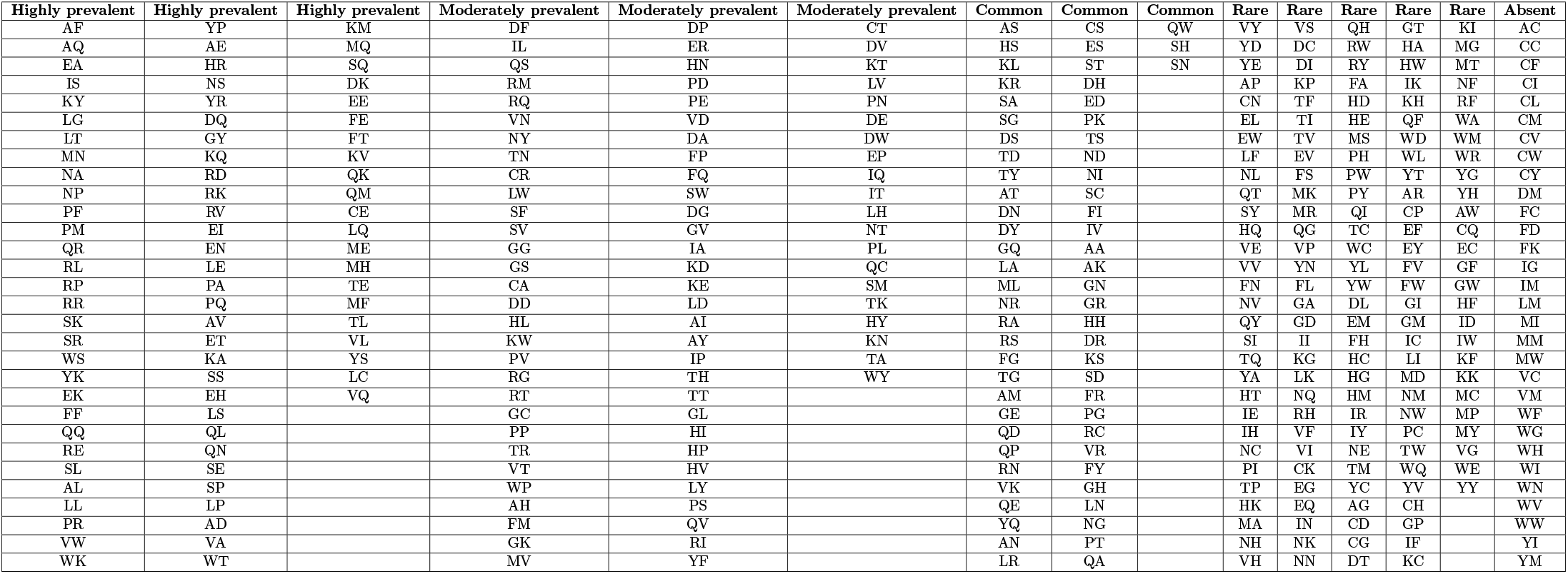
Prevalence categories of the 400 dipeptides across SRY protein sequences. Categories: highly prevalent (>75%), common (51–75%), moderately prevalent (26–50%), rare (*≤*25%), and absent (0%).

Based on frequency of dipeptide composition across 65 SRY sequences from various organisms led to a dendrogram relationship among SRY sequences (Figure 14). By setting a distance cutoff of 3.0, thirteen clusters of proximal SRY sequences were derived from the dendrogram. The list of clusters were presented in Table 16.

**Figure 14:**
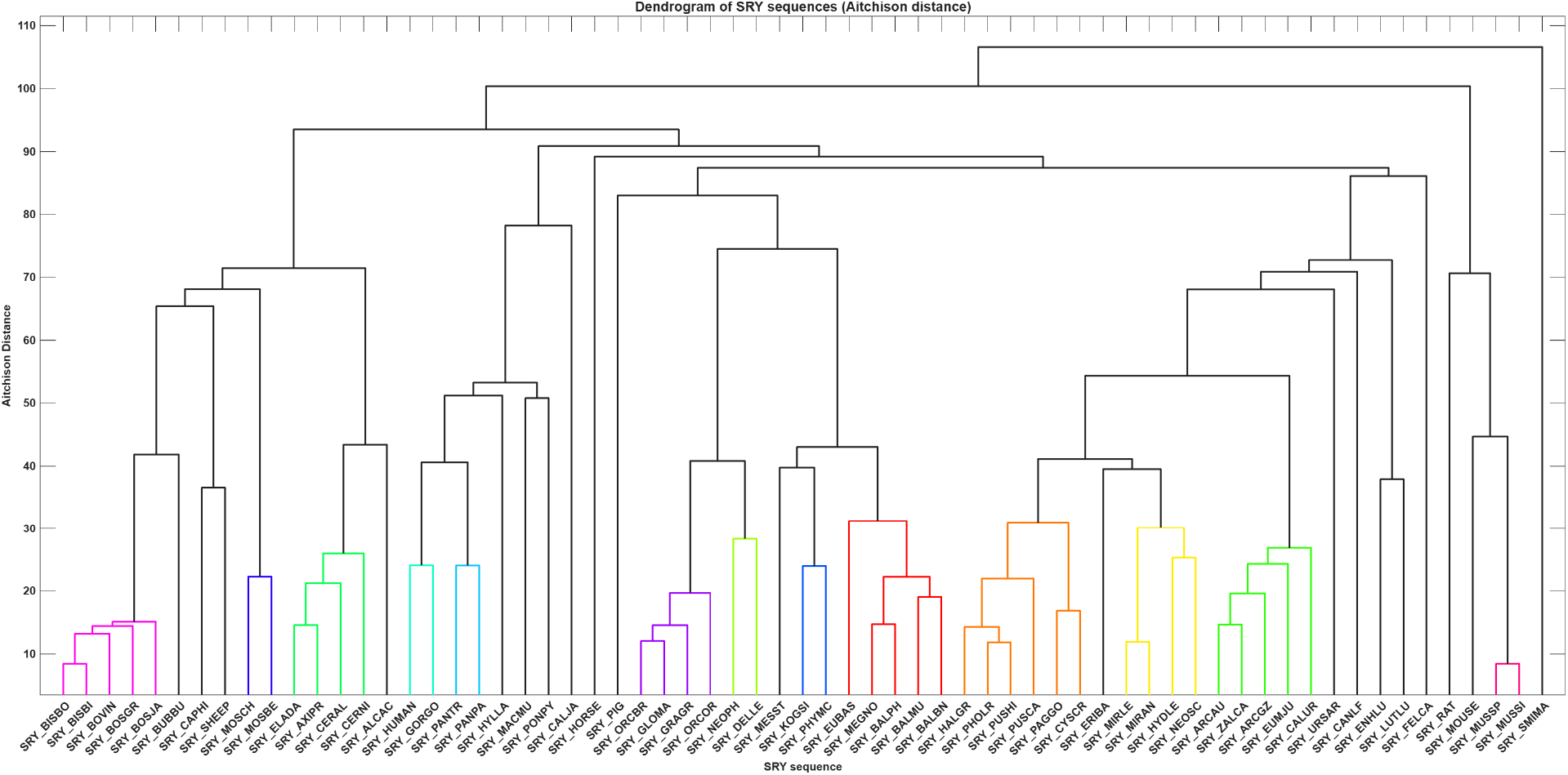
Dendrogram of SRY protein sequences based on dipeptide composition. Hierarchical clustering was performed using average linkage on pairwise Aitchison distances derived from centered log-ratio (CLR) transformed dipeptide frequencies. Colored branches highlight sequence groups resolved at lower distances, revealing clear taxonomic structuring among SRY proteins.

**Table 16:** List of proximal SRY sequences based on dipeptide frequency composition proximity.

| Cluster | Sets of SRY sequences | Cluster | Sets of SRY sequences |
| --- | --- | --- | --- |
| Cluster-1 | {SRY_BISBO, SRY_BISBI, SRY_BOVIN, SRY_BOSGR, SRY_BOSJA} | Cluster-8 | {SRY_KOGSI, SRY_PHYMC} |
| Cluster-2 | {SRY_MOSCH, SRY_MOSBE} | Cluster-9 | {SRY_EUBAS, SRY_MEGNO, SRY_BALPH, SRY_BALMU, SRY_BALBN} |
| Cluster-3 | {SRY_ELADA, SRY_AXIPR, SRY_CERAL, SRY_CERNI} | Cluster-10 | {SRY_HALGR, SRY_PHOLR, SRY_PUSHI, SRY_PUSCA, SRY_PAGGO, SRY_CYSCR} |
| Cluster-4 | {SRY_HUMAN, SRY_GORGO} | Cluster-11 | {SRY_MIRLE, SRY_MIRAN, SRY_HYDLE, SRY_NEOSC} |
| Cluster-5 | {SRY_PANTR, SRY_PANPA} | Cluster-12 | {SRY_ARCAU, SRY_ZALCA, SRY_ARCGZ, SRY_EUMJU, SRY_CALUR} |
| Cluster-6 | {SRY_ORCBR, SRY_GLOMA, SRY_GRAGR, SRY_ORCOR} | Cluster-13 | {SRY_MUSSP, SRY_MUSSI} |
| Cluster-7 | {SRY_NEOPH, SRY_DELLE} |  |  |

#### 4.6.2. Prevalence of dipeptide frequency and associated dendrogram of SRY-HMG subsequences

The 400 dipeptides were classified into five prevalence categories based on their occurrence across the 38 SRY-HMG domain subsequences (Table 17). The distribution was still skewed toward low prevalence, though it was somewhat less extreme than that observed for the full-length SRY proteins, a pattern that reflected the shorter and more uniform length of the HMG domain subsequences. The rare category (*≤*25%) still contained the largest number of dipeptides, followed by a substantial set that was completely absent, which indicated that many amino acid couple combinations did not occur within the HMG domain. Only a small subset was highly prevalent (>75%), representing couples that were strongly enriched in this domain.

**Table 17:** Prevalence categories of the 400 dipeptides across SRY-HMG domain subsequences. Categories: highly prevalent (>75%), common (51–75%), moderately prevalent (26–50%), and rare (*≤*25%).

| Highly prevalent | Highly prevalent | Highly prevalent | Moderately prevalent | Moderately prevalent | Common | Rare | Rare | Rare | Rare | Rare | Rare |
| --- | --- | --- | --- | --- | --- | --- | --- | --- | --- | --- | --- |
| AF | SR | SE | QM | DA | EE | RN | SD | SM | TQ | HN | WE |
| EA | KY | RK | YQ | EW | FE | VR | VH | YD | YR | HP | WL |
| IS | NS | RE | IV | FQ | VA | IH | DH | CR | AA | ID | WR |
| MN | VW | RL | KW | NY | QN | MA | DI | FV | AD | KI |  |
| NA | WK | QL | WP | PN | TE | MR | DW | IK | AG | LP |  |
| PF | YK | KM | FM | TD | VK | QK | GE | IL | AK | LV |  |
| PM | YP | KR | DK | YE | ML | AI | HK | IQ | AT | NK |  |
| RP | AE |  | MV |  | KV | AV | HQ | IR | CK | QV |  |
| SK | EI |  | DY |  | DQ | KN | KH | KT | CQ | RQ |  |
| WS | HR |  | KL |  | LQ | LR | KS | LS | DE | SI |  |
| AQ | RR |  | PG |  | MH | MK | LK | LW | DS | SQ |  |
| EK | AL |  | ER |  | MQ | GC | PR | NT | EM | TA |  |
| FF | GY |  | LA |  | PQ | LH | PS | QD | EY | TH |  |
| LG | EN |  | PD |  | AM | QP | QQ | QT | FL | TW |  |
| LT | KQ |  | LL |  | FI | RA | RG | RH | GK | VE |  |
| NP | LE |  | PK |  | QA | RM | RW | RI | GV | VS |  |
| QR | RD |  | QE |  | QW | RV | SL | TL | HL | VT |  |

When examined as amino acid couples, the highly prevalent dipeptides were dominated by polar–polar and polar–charged combinations, along with several couples in which a polar residue was paired with a small nonpolar residue. This pattern was consistent with the solvent-exposed and DNA-contacting surfaces of the HMG box. The common category (51–75%) followed the same trend, in which polar–polar and polar–charged couples prevailed. Moderately prevalent dipeptides (26–50%) showed a more balanced mix of polar–nonpolar and nonpolar–nonpolar couplings. Furthermore, the rare and absent categories were enriched for couples in which both partners were bulky nonpolar residues, or that involved cysteine and other residues with limited compatibility in the SRY-HMG fold.

Therefore, the prevalence distribution revealed a clear compositional bias within the SRY-HMG domain: polar- and charged-coupled dipeptides were widely conserved, whereas nonpolar–nonpolar and sterically demanding couples were largely excluded, consistent with strong purifying selection acting on the functional core of the domain. Notably, this bias was more pronounced than in the SRY proteins, where a higher proportion of rare and absent dipeptides was observed; the difference reflected the fact that the HMG domain was a shorter, structurally homogeneous unit, whereas the SRY sequences included the variable N- and C-terminal regions that diluted the conserved compositional signature.

Furthermore, eight clusters of SRY-HMG subsequences were derived from the dendrogram relationship derived from dipeptide composition of amino acids across 38 SRY-HMG subsequences (Figure 15, Table 18).

**Figure 15:**
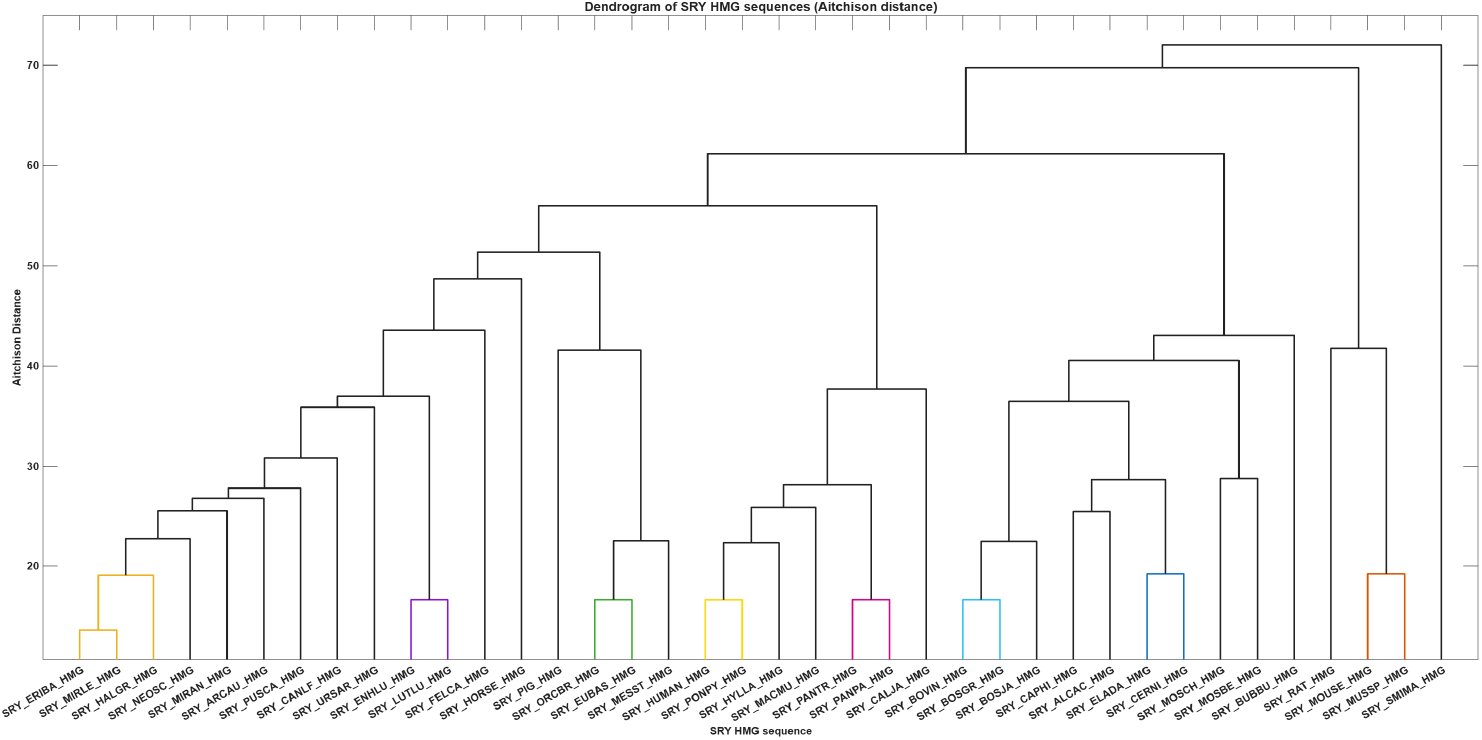
Dendrogram of SRY HMG sequences based on dipeptide composition. Hierarchical clustering was performed using average linkage on pairwise Aitchison distances derived from dipeptide frequencies. Sequence labels correspond to SRY HMG protein sequences from 38 mammalian organisms.

**Table 18:**
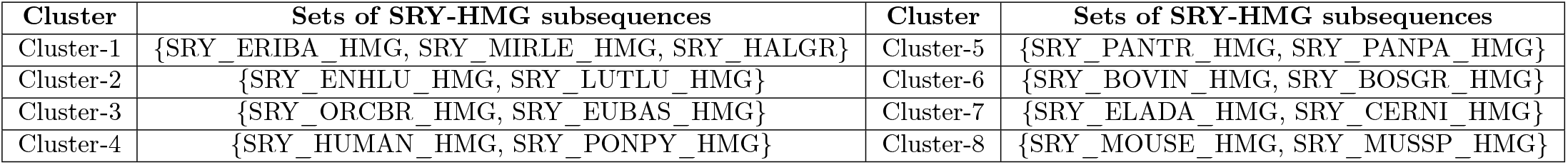
List of proximal SRY-HMG subsequences based on dipeptide frequency composition proximity.

| Cluster | Sets of SRY-HMG subsequences | Cluster | Sets of SRY-HMG subsequences |
| --- | --- | --- | --- |
| Cluster-1 | {SRY_ERIBA_HMG, SRY_MIRLE_HMG, SRY_HALGR} | Cluster-5 | {SRY_PANTR_HMG, SRY_PANPA_HMG} |
| Cluster-2 | {SRY_ENHLU_HMG, SRY_LUTLU_HMG} | Cluster-6 | {SRY_BOVIN_HMG, SRY_BOSGR_HMG} |
| Cluster-3 | {SRY_ORCBR_HMG, SRY_EUBAS_HMG} | Cluster-7 | {SRY_ELADA_HMG, SRY_CERNI_HMG} |
| Cluster-4 | {SRY_HUMAN_HMG, SRY_PONPY_HMG} | Cluster-8 | {SRY_MOUSE_HMG, SRY_MUSSP_HMG} |

### 4.7. Dendrogram relationships across SRY and SRY-HMG sequences derived from the longest polystring analysis

#### 4.7.1. Dendrogram relationships across SRY sequences derived from the longest polystring analysis

A longest polystring analysis was performed for each of the 20 standard amino acids across 65 SRY protein sequences. Sequence lengths ranged from 169 residues (SRY_RAT) to 392 residues (SRY_MOUSE), with most sequences between 200 and 240 residues; the extreme length of mouse SRY reflected its extended C-terminal region. Across all sequences, homopolymer runs were generally short, with most amino acids showing maximum run lengths of 1–2 residues and normalized polystring lengths below 1.5%.

The longest runs were observed for glutamine and histidine, and were largely restricted to the murine lineage (Figure 16). Glutamine runs reached 13 residues in SRY_MOUSE, 8 in SRY_MUSSP, and 7 in SRY_RAT and SRY_MUSSI. Histidine runs reached 5 residues in SRY_MOUSE and 4 in SRY_MUSSP and SRY_MUSSI. Moderate runs of 3 residues occurred for leucine (SRY_MESST, SRY_KOGSI, SRY_PHYMC), valine (SRY_ORCBR, SRY_GLOMA, SRY_ORCOR, SRY_GRAGR), arginine (Bovini, Caprini, Moschidae, Cervidae, and several carnivore and phocid sequences), and serine (primates and several other lineages). Additionally, cysteine, tryptophan, and tyrosine never exceeded a run length of 1 in any sequence (Figure 16).

**Figure 16:**
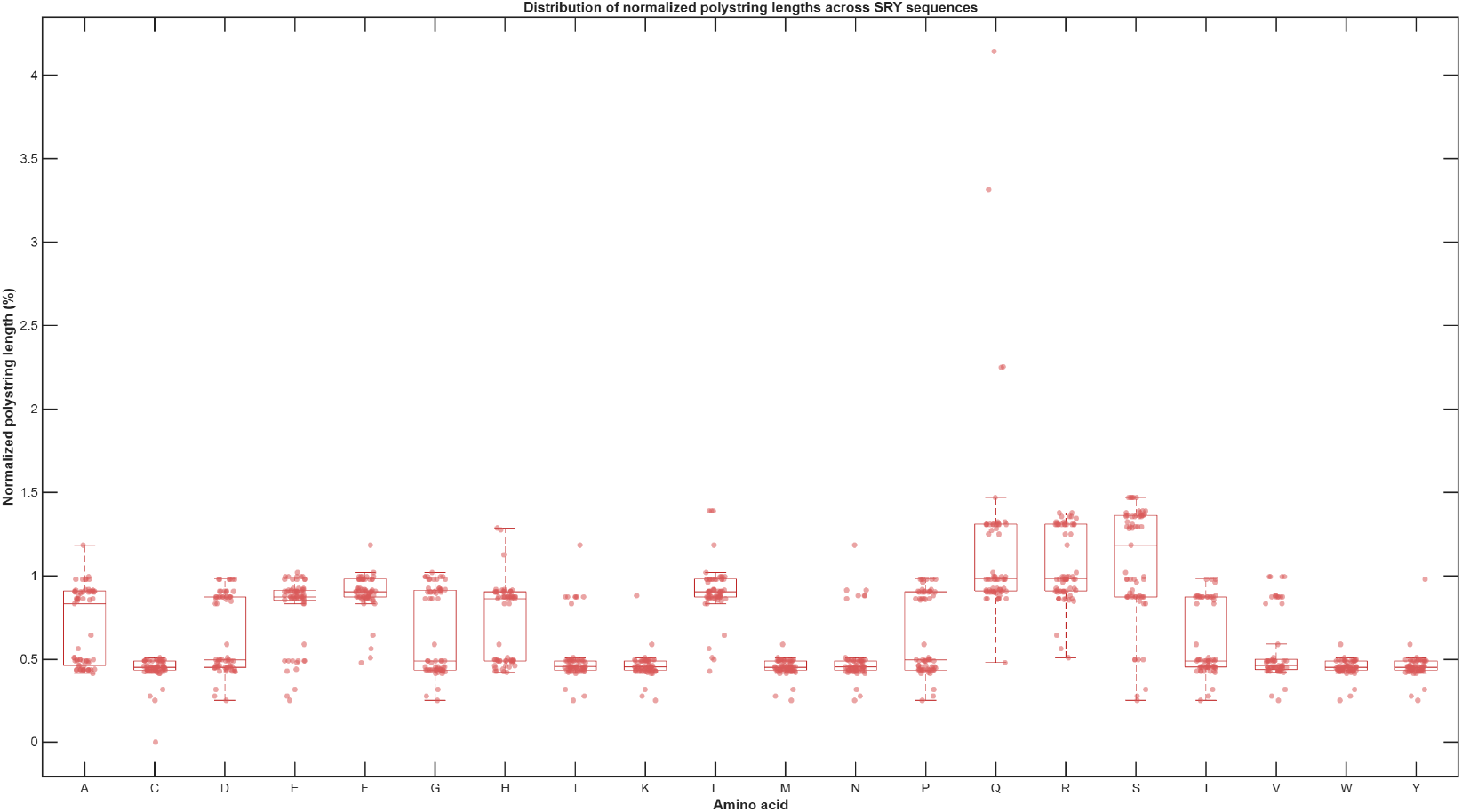
Distribution of normalized polystring lengths across SRY protein sequences. Each box represents the interquartile range (IQR) for one of the 20 standard amino acids, with the central line indicating the median and whiskers extending to the most extreme values within 1.5*×*IQR.

The distribution of normalized polystring lengths was strongly right-skewed. Q, S, R, and P showed the highest medians and widest spread, while cysteine showed the lowest values overall (Figure 16). The single highest normalized value was Q in SRY_RAT (*∼*4.14%), arising because the SRY_RAT sequence was the shortest (169 residues); although SRY_MOUSE carried the longest absolute Q run (13 residues), its longer length (392 residues) yielded a lower normalized value (*∼*3.32%).

It was noticed that SRY homopolymer runs were short and compositionally constrained, consistent with the structural demands of the HMG box, and the only extended runs occurred in the murine lineage, reflecting its distinctive Q-rich and H-rich C-terminal region. The depletion of cysteine, tryptophan, and tyrosine runs, together with the predominance of polar and small residues, was consistent with the strong structural constraints on the domain. Because the normalized metric was length-dependent, the same absolute run length contributed a higher percentage in shorter sequences such as SRY_RAT, a point to consider when comparing across taxa.

Based on the proximity of the longest normalized frequuency of polystring of 20 amino acid profiles, a dendrogram relationship among SRY sequences was developed (Figure 17). By setting a distance threshold of 0.5, 13 clusters were obtained, which was summarized in Table 19.

**Figure 17:**
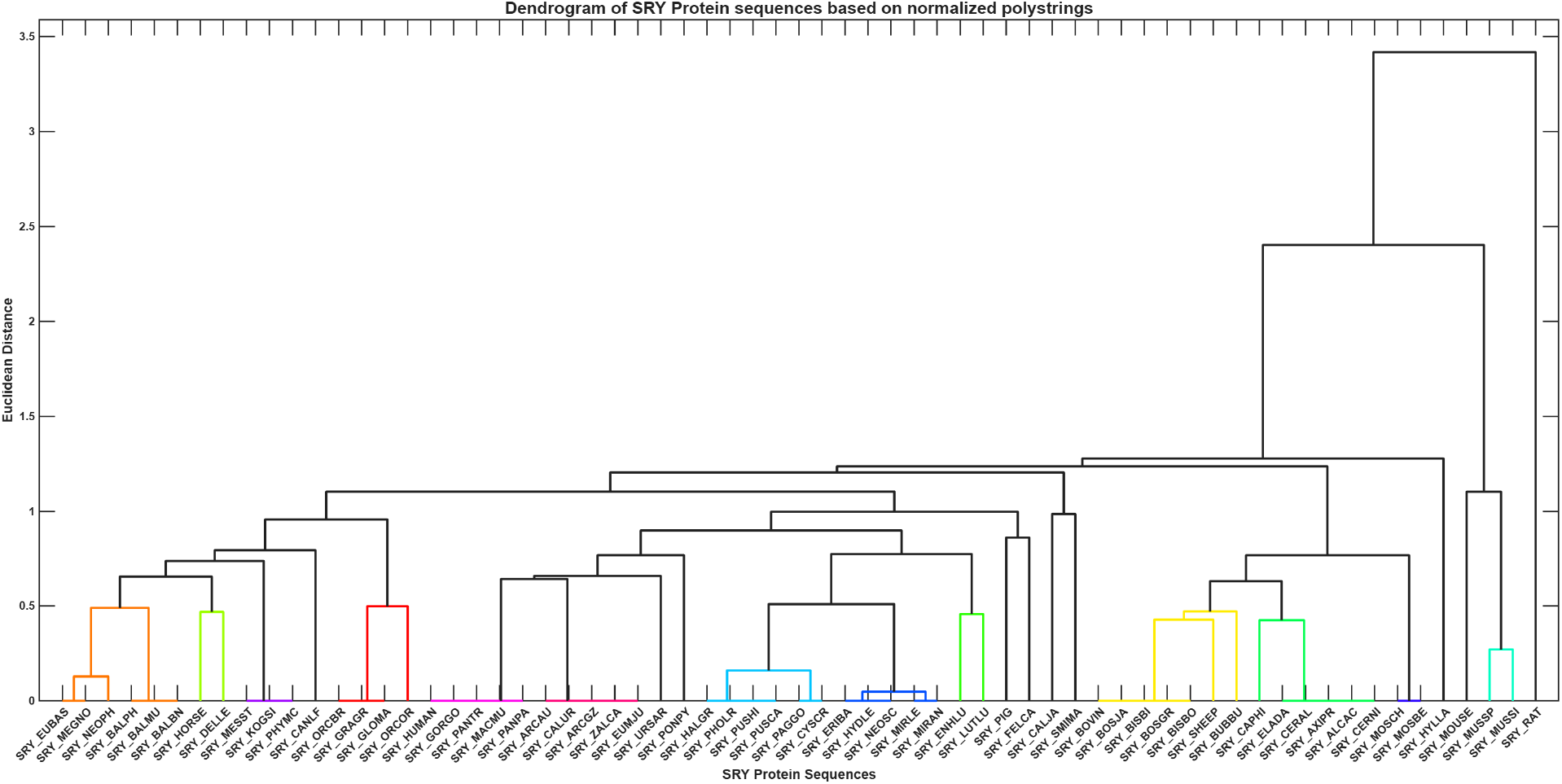
Dendrogram of SRY protein sequences based on the longest normalized polystring lengths. Hierarchical clustering was performed using average linkage (UPGMA). Colored branches indicate subclusters defined at a distance threshold of 0.5, used solely for visual grouping.

**Table 19:**
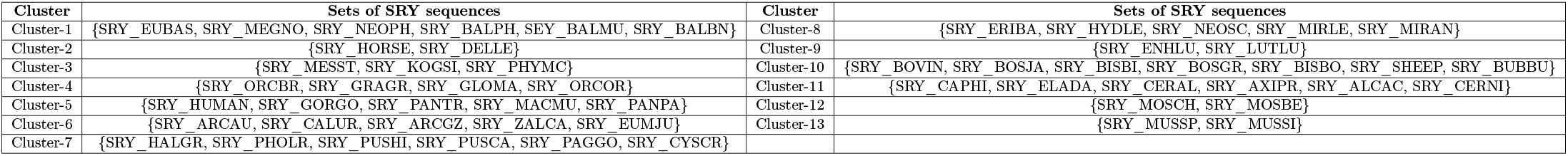
List of proximal of SRY sequences based on the longest normalized polystring lengths.

#### 4.7.2. Dendrogram relationships across SRY-HMG subsequences derived from the longest polystring analysis

A longest polystring analysis was performed for each of the 20 standard amino acids across 38 SRY-HMG domain subsequences. Across all SRY-HMG subsequences, homopolymer runs were uniformly short, with maximum run lengths of 1–3 residues and normalized polystring lengths generally between 1.4% and 4.4% (Figure 18).

**Figure 18:**
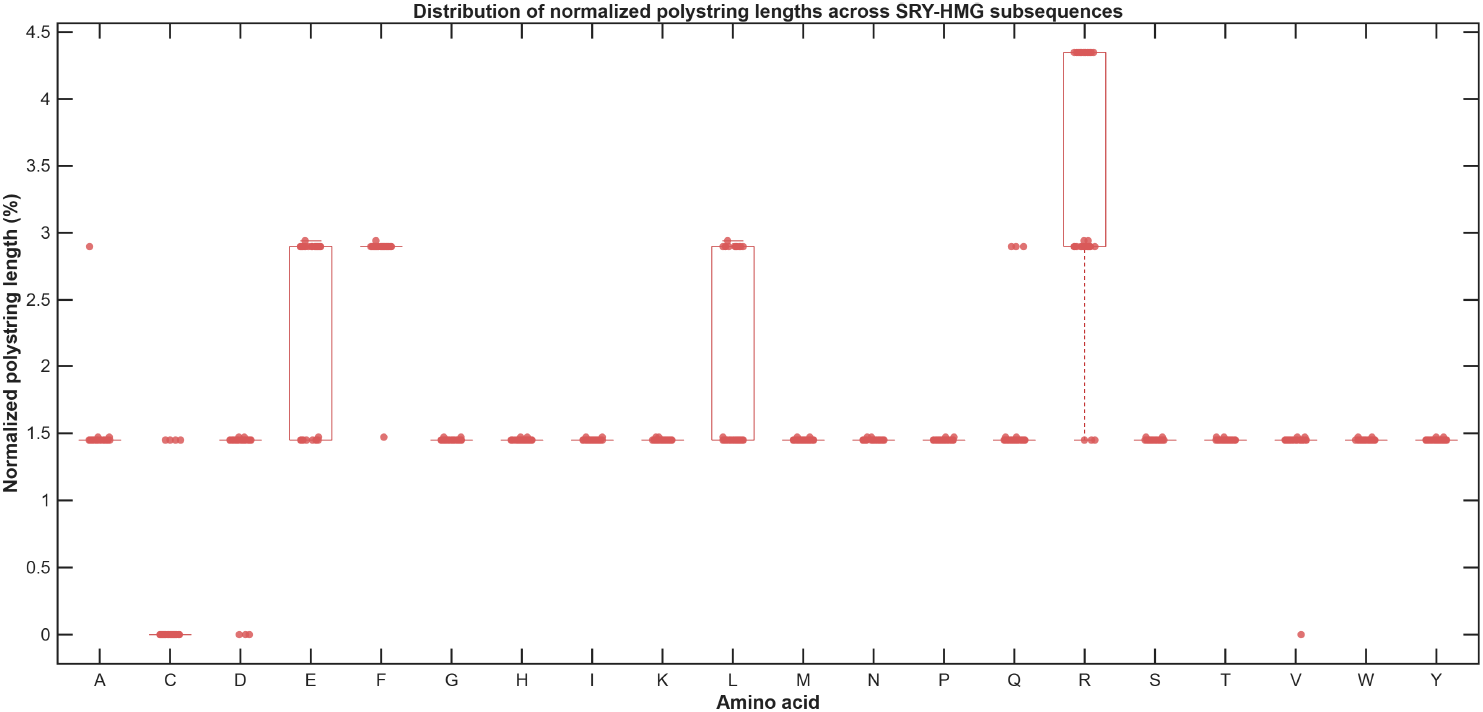
Distribution of normalized polystring lengths across SRY-HMG subsequences. Boxes show the interquartile range with median lines; individual values are overlaid as jittered points.

The longest runs were restricted to a small number of amino acids and lineages. Arginine reached 3 residues (RRR) in a defined set of sequences representing the Bovini (SRY_BOVIN_HMG, SRY_BOSJA_HMG, SRY_BOSGR_HMG, SRY_BISBI_HMG), Caprini (SRY_CAPHI_HMG, SRY_SHEEP_HMG), Moschidae (SRY_MOSCH_HMG, SRY_MOSBE_HMG), Cervidae (SRY_ELADA_HMG, SRY_ALCAC_HMG, SRY_CERNI_HMG), and several carnivore and phocid sequences (SRY_ERIBA_HMG, SRY_MIRLE_HMG, SRY_MIRAN_HMG, SRY_NEOSC_HMG). Glutamine reached 2 residues (QQ) only in the rodent sequences (SRY_RAT_HMG, SRY_MOUSE_HMG, SRY_MUSSP_HMG), and leucine reached 2 residues (LL) in 12 sequences spanning primates, ruminants, and carnivores. The remaining amino acids never exceeded a run length of 1. Cysteine was entirely absent (C = 0) in 34 of the 38 sequences, and valine was absent in SRY_BUBBU_HMG only (Figure 18). The distribution of normalized polystring lengths was narrow and strongly skewed toward low values. The highest normalized value was arginine (*∼*4.35%) in the sequences carrying the RRR motif, followed by leucine (*∼*4.35%) in the LL sequences and glutamine (*∼*2.90–2.94%) in the rodent sequences. The lowest values were observed for cysteine, reflecting its near or complete absence in the HMG domain (Figure 18).

It was found that SRY-HMG subsequences display a highly restricted polystring profile: runs are short, predominantly 1–2 residues, and the only recurrent 3-residue motif is RRR, confined to specific artiodactyl and carnivore lineages. The near absence of cysteine and the absence of tryptophan and tyrosine runs reflected the strong structural and compositional constraints on the HMG box, consistent with its role as a sequence-specific DNA-binding domain.

Two key differences were noticed while we comparing the polystring profiles between full-length SRY and associated SRY-HMG subsequences. Full-length SRY contained extended runs of up to 13 residues for glutamine and 5 for histidine, all confined to the murine C-terminal tail, while SRY-HMG runs were limited to 3 residues (RRR) and 2 residues (QQ, LL). Secondly, although cysteine, tryptophan, and tyrosine never exceeded a run length of 1 in either SRY or SRY-HMG sequences, the constraint was stronger in SRY-HMG, where cysteine was entirely absent in 34 of 38 sequences and the only recurrent 3-residue motif was RRR, compared with LLL, VVV, and RRR in full-length SRY. Consequently, the full-length SRY polystring profile was dominated by its variable N- and C-terminal regions, whereas the HMG domain displayed a short, uniform, and compositionally constrained signature.

Based on the proximity of the longest normalized frequuency of polystring of 20 amino acid profiles, a dendrogram relationship among SRY-HMG subsequences was developed (Figure 19). By setting a distance threshold of 0.5, six clusters were obtained, which was summarized in Table 20.

**Figure 19:**
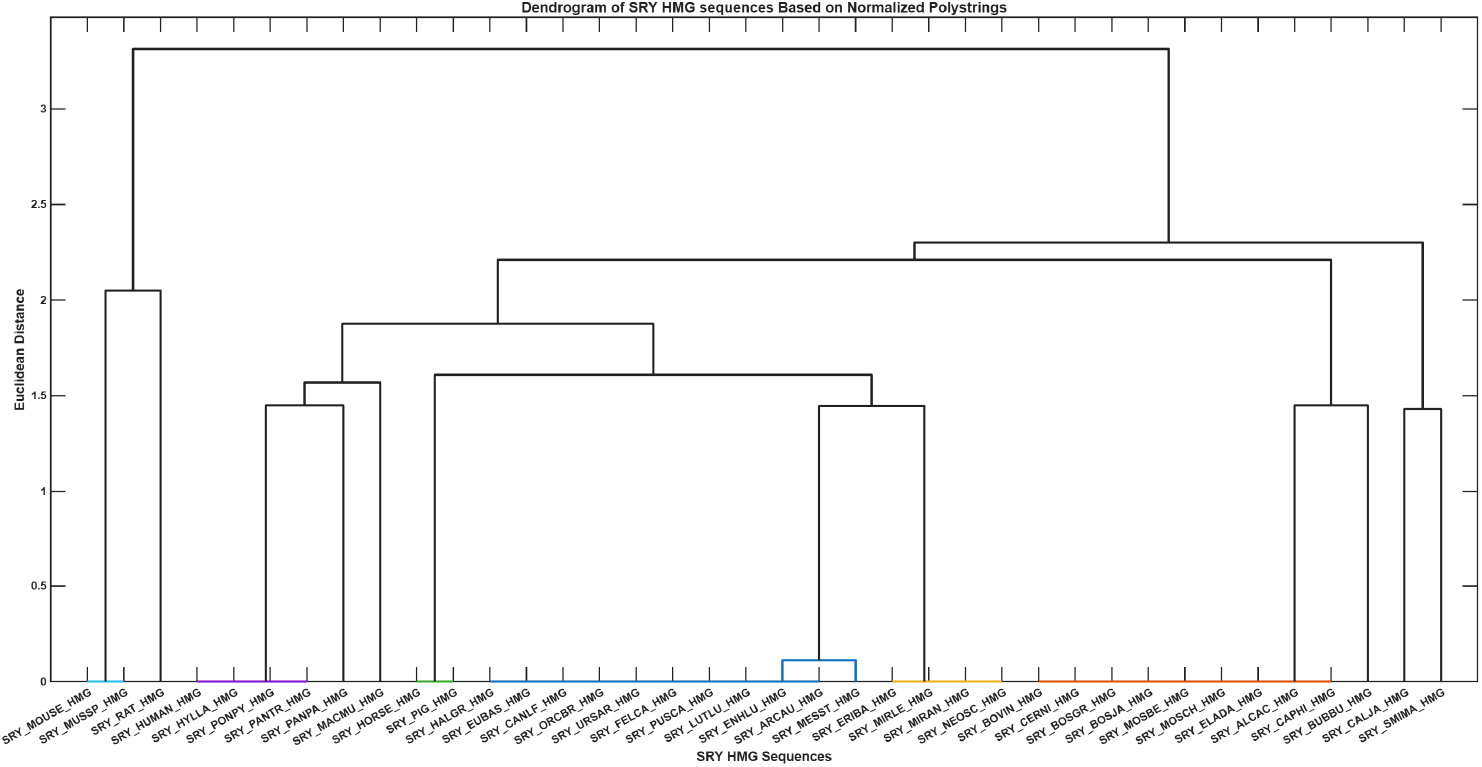
Dendrogram of SRY-HMG subsequences based on normalized polystring lengths. Euclidean distances were computed from a 20-dimensional feature vector per sequence, and clustering used average linkage.

**Table 20:**
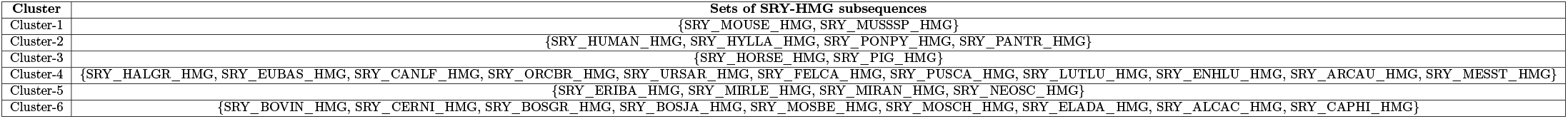
List of proximal SRY-HMG subsequences based on the longest normalized polystring lengths.

### 4.8. Analysis of hydropathy-based n-gram frequency across SRY and SRY-HMG sequences

#### 4.8.1. Analysis of hydropathy-based n-gram frequency across SRY sequences

##### Relative frequency of 2-grams and associated phylogenetic relationships of SRY sequences

For each of the 65 SRY sequences, the binary hydropathy encoding was used to enumerate all overlapping 2-grams. Let *F_w_* denote the relative frequency of the 2-gram *w ∈ {*00, 01, 10, 11*}*, where 0 denotes a nonpolar residue and 1 denotes a polar residue. Thus *F*_00_, *F*_01_, *F*_10_, and *F*_11_ correspond to the relative frequencies of polar–polar, nonpolar–polar, polar– nonpolar, and nonpolar–nonpolar pairs, respectively. These frequencies were computed and summarized (Table 21). The 2-gram profile was dominated by 11, with a mean relative frequency of 0.4026 (range 0.3250–0.5934). The transition words 01 and 10 were the next most frequent, with means of 0.2198 (range 0.1714–0.2600) and 0.2168 (range 0.1714–0.2550), respectively (Figure 20). The polar–polar word 00 was least frequent, with a mean of 0.1608 (range 0.0639–0.2078). The close agreement between 01 and 10 frequencies (mean difference = 0.0030) is consistent with the binary encoding of linear sequences, in which the counts of the two transition types can differ by at most one.

**Figure 20:**
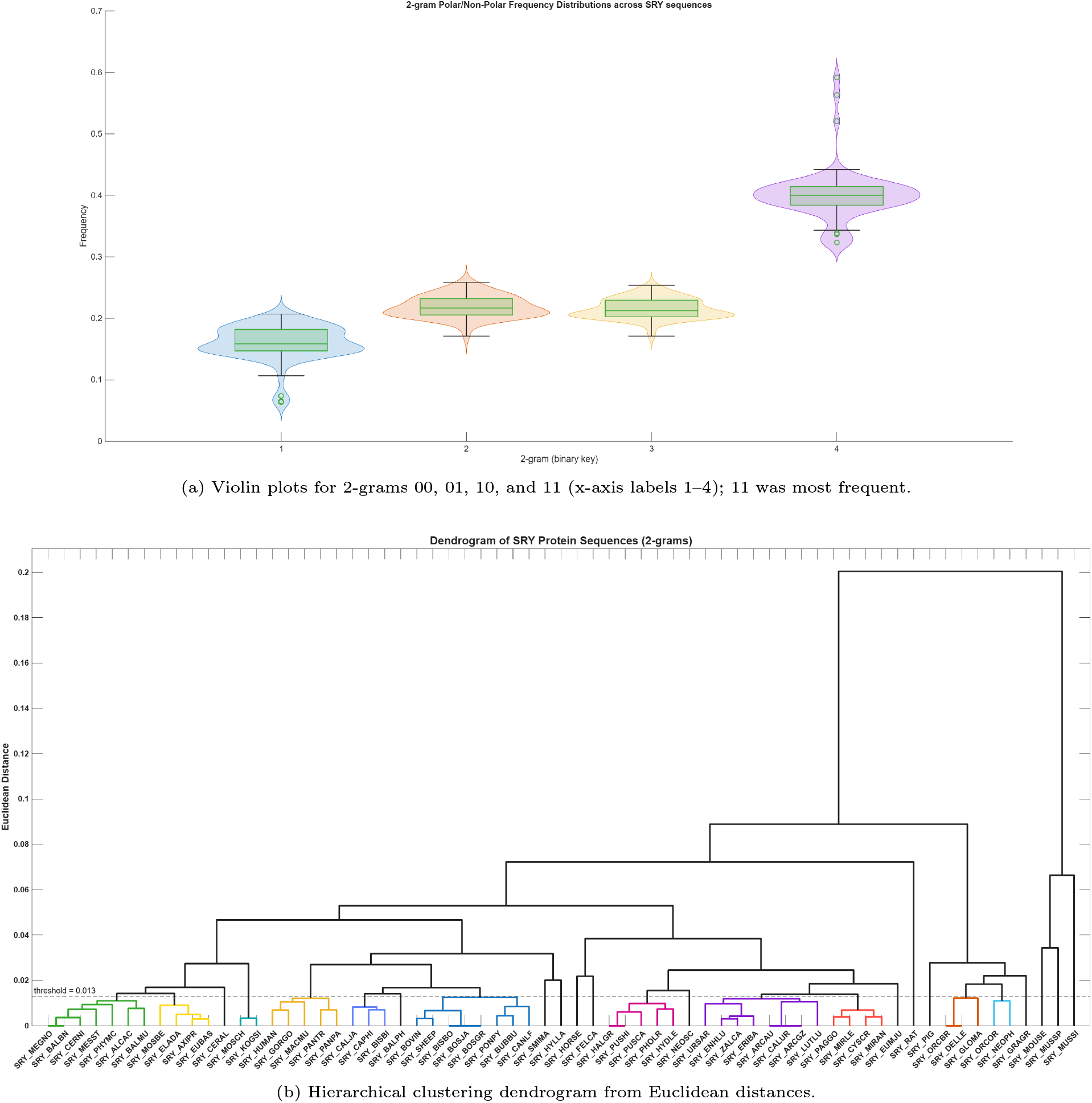
Hydropathy-based 2-gram frequency profiles of 65 SRY sequences.

**Table 21:** Summary of hydropathy-based 2-gram relative frequencies across 65 SRY sequences.

| 2-gram | Hydropathy pattern | Mean | Range | Sequence(s) with minimum / maximum |
| --- | --- | --- | --- | --- |
| 00 | Polar–polar | 0.1608 | 0.0639–0.2078 | Min: SRY_MOUSE; Max: SRY_PUSCA |
| 01 | Nonpolar–polar | 0.2198 | 0.1714–0.2600 | Min: SRY_MOUSE; Max: SRY_GRAGR |
| 10 | Polar–nonpolar | 0.2168 | 0.1714–0.2550 | Min: SRY_MOUSE; Max: SRY_GRAGR |
| 11 | Nonpolar–nonpolar | 0.4026 | 0.3250–0.5934 | Min: SRY_ORCBR, SRY_GLOMA, SRY_GRAGR, SRY_DELLE; Max: SRY_MOUSE |

Substantial variation was observed among SRY sequences. The most nonpolar-enriched profiles were found in SRY_MOUSE (*F*_11_ = 0.5934), SRY_MUSSP (0.5650), and SRY_MUSSI (0.5226), which also had the lowest 00 frequencies (0.0639, 0.0650, and 0.0742, respectively) and the lowest 01/10 frequencies (SRY_MOUSE: 0.1714*/*0.1714). On the other side, the lowest *F*_11_ values (0.3250) occurred in SRY_ORCBR, SRY_GLOMA, SRY_GRAGR, and SRY_DELLE, with SRY_GRAGR showing the highest transition frequencies (*F*_01_ = 0.2600, *F*_10_ = 0.2550) Figure 20). The highest *F*_00_ was observed in SRY_PUSCA (0.2078), followed by SRY_HALGR and SRY_PUSHI (0.2035 each), indicating a relatively higher proportion of consecutive polar residues in these sequences.

Therefore, the 65 SRY sequences exhibited a nonpolar-biased hydropathy 2-gram profile, with nonpolar–nonpolar pairs being the most common local pattern, transitions being intermediate, and polar–polar pairs being the least common. These relative frequencies were used as feature vectors to compute pairwise Euclidean distances among all 65 SRY sequences, producing the 65 *×* 65 distance matrix that was subsequently subjected to hierarchical clustering as presented in Figure 20). List of proximal SRY sequences based on proximity of 2-grams hydropathy profiles was summarized in Table 22.

**Table 22:**
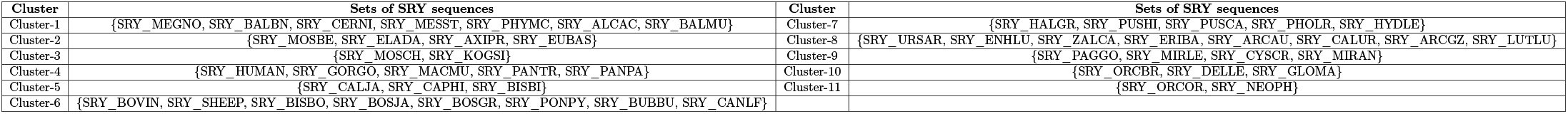
List of proximal SRY sequences based on proximity of 2-grams hydropathy profiles.

##### Relative frequency of 3-grams and associated phylogenetic relationships of SRY sequences

For each of the 65 SRY sequences, all overlapping 3-grams were enumerated from the binary hydropathy encoding. Let *F_w_* denote the relative frequency of the 3-gram *w ∈ {*000, 001*, . . . ,* 111*}*, where 0 denotes a nonpolar residue and 1 denotes a polar residue. Thus, for example, 000 represents nonpolar–nonpolar–nonpolar, 001 represents nonpolar–nonpolar–polar, and 111 represents polar–polar–polar.

The 3-gram profile was dominated by 111 (polar–polar–polar), with a mean relative frequency of 0.2494 (range 0.1658– 0.4769). The next most frequent patterns were 011 (nonpolar–polar–polar) and 110 (polar–polar–nonpolar), with means of 0.1550 and 0.1540, respectively (Figure 21). The pattern 101 (polar–nonpolar–polar) had a mean of 0.1255, followed by 001 (nonpolar–nonpolar–polar) and 100 (polar–nonpolar–nonpolar) with means of 0.0942 and 0.0916. The least frequent 3-grams were 000 (nonpolar–nonpolar–nonpolar) and 010 (nonpolar–polar–nonpolar), with means of 0.0665 and 0.0637, respectively (Figure 21). Overall, the 65 SRY sequences exhibited a strongly polar-biased hydropathy 3-gram profile (Table 23).

**Figure 21:**
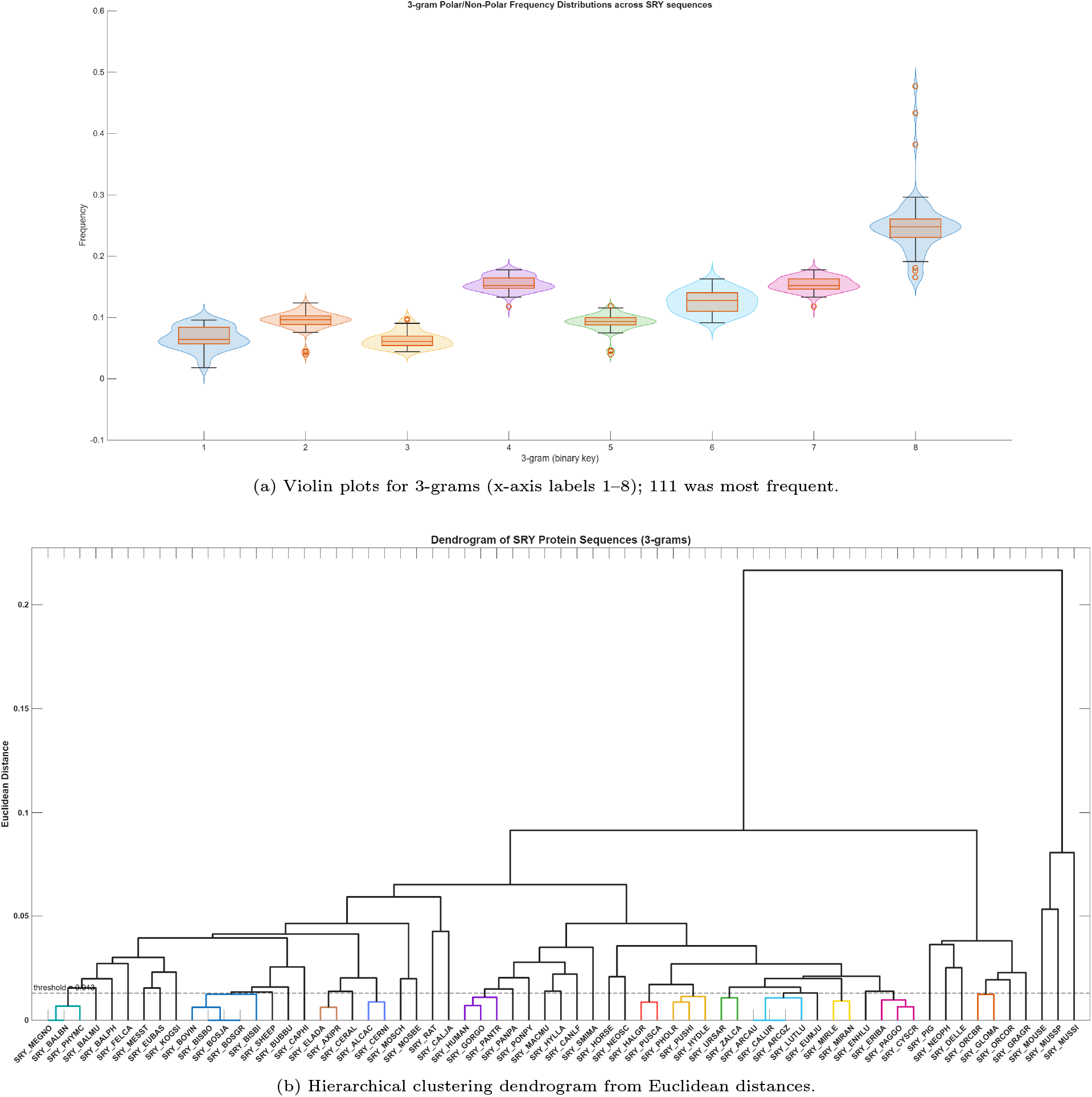
Hydropathy-based 3-gram frequency profiles of 65 SRY sequences.

**Table 23:** Summary of hydropathy-based 3-gram relative frequencies across 65 SRY sequences.

| 3-gram | Hydrophathy pattern | Mean | Range | Min sequence | Max sequence |
| --- | --- | --- | --- | --- | --- |
| 000 | Nonpolar-nonpolar-nonpolar | 0.0665 | 0.0179–0.0957 | *SRY_MOUSE* | *SRY_PUSHI* |
| 001 | Nonpolar-nonpolar-polar | 0.0942 | 0.0397–0.1237 | *SRY_MUSSP* | *SRY_NEOPH* |
| 010 | Nonpolar-polar-nonpolar | 0.0637 | 0.0441–0.0979 | *SRY_BUBBU* | *SRY_NEOPH* |
| 011 | Nonpolar-polar-polar | 0.1550 | 0.1179–0.1778 | *SRY_MOUSE* | *SRY_MOSCH* |
| 100 | Polar-nonpolar-nonpolar | 0.0916 | 0.0397–0.1186 | *SRY_MUSSP* | *SRY_NEOPH* |
| 101 | Polar-nonpolar-polar | 0.1255 | 0.0913–0.1630 | *SRY_PUSCA* | *SRY_CERAL* |
| 110 | Polar-polar-nonpolar | 0.1540 | 0.1179–0.1778 | *SRY_MOUSE* | *SRY_MOSCH* |
| 111 | Polar-polar-polar | 0.2494 | 0.1658–0.4769 | *SRY_ORCBR*, *SRY_GLOMA*, *SRY_GRAGR* | *SRY_MOUSE* |

Considerable variation was observed among SRY sequences, as noticed in 2-grams. The most polar-enriched profiles were found in SRY_MOUSE (*F*_111_ = 0.4769), SRY_MUSSP (0.4334), and SRY_MUSSI (0.3819). In contrast, the lowest *F*_111_ values (0.1658) occurred in SRY_ORCBR, SRY_GLOMA, and SRY_GRAGR. SRY_NEOPH showed the highest frequencies of 001 (0.1237), 010 (0.0979), and 100 (0.1186), indicating a more mixed hydropathy pattern. The highest 011 and 110 frequencies were observed in SRY_MOSCH (0.1778 for both patterns), whereas the lowest 011 and 110 frequencies occurred in SRY_MOUSE (0.1179 for both) (Table 23).

These relative frequencies were used as feature vectors to compute pairwise Euclidean distances among all 65 SRY sequences, producing the 65 *×* 65 distance matrix that was subsequently subjected to hierarchical clustering (Figure 21), which was presented in Table 24.

**Table 24:**
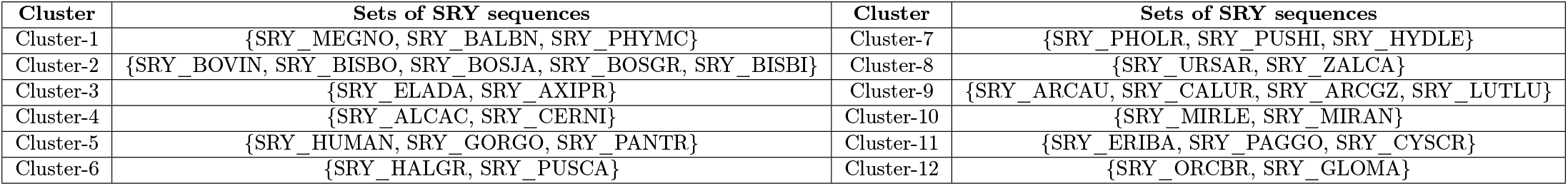
List of proximal SRY sequences based on proximity of 3-grams hydropathy profiles.

##### Relative frequency of 4-grams and associated phylogenetic relationships of SRY sequences

The binary hydropathy profiles of all 65 SRY sequences were scanned with a sliding window of length four to extract the complete set of overlapping 4-grams. For a given sequence, *F_w_* denotes the relative frequency of the 4-gram *w ∈ {*0000, 0001*, . . . ,* 1111*}*, with 0 and 1 representing nonpolar and polar residues, respectively.

A clear polar bias emerged from the 4-gram spectra across SRY sequences. The pattern 1111 (polar–polar–polar–polar) was the single most abundant word, averaging 0.1495 across sequences and ranging from 0.0859 to 0.3856. It was followed jointly by 0111 (nonpolar–polar–polar–polar) and 1110 (polar–polar–polar–nonpolar), each with a mean of 0.1010 and an identical range of 0.0808–0.1194. The patterns 1101 and 1011 occupied the next tier, averaging 0.0968 and 0.0849, respectively. At the opposite end of the spectrum, 0010 (nonpolar–nonpolar–polar–nonpolar) was the rarest word (mean 0.0239), marginally below 0000 (mean 0.0266) (Table 25).

**Table 25:** Summary of hydropathy-based 4-gram relative frequencies across 65 SRY sequences.

| 4-gram | Hydropathy pattern | Mean | Range | Min sequence(s) | Max sequence(s) |
| --- | --- | --- | --- | --- | --- |
| 0000 | Nonpolar–nonpolar–nonpolar–nonpolar | 0.0266 | 0.0077–0.0419 | SRY_MOUSE | SRY_MIRLE, SRY_MIRAN |
| 0001 | Nonpolar–nonpolar–nonpolar–polar | 0.0397 | 0.0103–0.0568 | SRY_MOUSE | SRY_PUSHI |
| 0010 | Nonpolar–nonpolar–polar–nonpolar | 0.0239 | 0.0084–0.0518 | SRY_CAPHI, SRY_SHEEP | SRY_NEOPH |
| 0011 | Nonpolar–nonpolar–polar–polar | 0.0700 | 0.0170–0.0917 | SRY_MUSSP | SRY_PUSCA |
| 0100 | Nonpolar–polar–nonpolar–nonpolar | 0.0346 | 0.0142–0.0622 | SRY_MUSSP | SRY_NEOPH |
| 0101 | Nonpolar–polar–nonpolar–polar | 0.0293 | 0.0100–0.0491 | SRY_PANTR, SRY_PANPA | SRY_CALJA |
| 0110 | Nonpolar–polar–polar–nonpolar | 0.0538 | 0.0257–0.0848 | SRY_MOUSE | SRY_MOSCH |
| 0111 | Nonpolar–polar–polar–polar | 0.1010 | 0.0808–0.1194 | SRY_ORCBR, SRY_GLOMA, SRY_GRAGR, SRY_DELLE | SRY_MEGNO, SRY_BALPH |
| 1000 | Polar–nonpolar–nonpolar–nonpolar | 0.0388 | 0.0103–0.0545 | SRY_MOUSE | SRY_HORSE |
| 1001 | Polar–nonpolar–nonpolar–polar | 0.0528 | 0.0284–0.0829 | SRY_MUSSP | SRY_NEOPH |
| 1010 | Polar–nonpolar–polar–nonpolar | 0.0399 | 0.0265–0.0556 | SRY_BUBBU | SRY_GLOMA, SRY_ORCOR, SRY_GRAGR |
| 1011 | Polar–nonpolar–polar–polar | 0.0849 | 0.0568–0.1234 | SRY_PUSCA | SRY_MUSSI |
| 1100 | Polar–polar–nonpolar–nonpolar | 0.0573 | 0.0231–0.0786 | SRY_MOUSE | SRY_PUSCA |
| 1101 | Polar–polar–nonpolar–polar | 0.0968 | 0.0699–0.1181 | SRY_PUSCA | SRY_CAPHI |
| 1110 | Polar–polar–polar–nonpolar | 0.1010 | 0.0808–0.1194 | SRY_ORCBR, SRY_GLOMA, SRY_GRAGR, SRY_DELLE | SRY_MEGNO, SRY_BALPH |
| 1111 | Polar–polar–polar–polar | 0.1495 | 0.0859–0.3856 | SRY_ORCBR, SRY_GLOMA, SRY_GRAGR | SRY_MOUSE |

Sequence-to-sequence heterogeneity was substantially observed. The strongest polar signature was carried by SRY_MOUSE, whose *F*_1111_ reached 0.3856, followed by SRY_MUSSP (0.3352) and SRY_MUSSI (0.2792). In addition, SRY_ORCBR, SRY_GLOMA, and SRY_GRAGR shared the lowest *F*_1111_ value (0.0859). Several sequences stood out for specific rare words: SRY_PUSCA had the highest 0011 (0.0917) and 1100 (0.0786) frequencies, SRY_NEOPH led in 0010 (0.0518), 0100 (0.0622), and 1001 (0.0829), SRY_MUSSI in 1011 (0.1234), and SRY_CAPHI in 1101 (0.1181). An interesting feature of this SRY sequences was that the 0111 and 1110 columns were numerically identical for every sequence, reflecting the near-symmetry of the two transition directions in the binary encoding (Table 25).

These sixteen-dimensional frequency vectors served as the input features for pairwise Euclidean distance calculation among the 65 SRY sequences, yielding the 65 *×* 65 distance matrix that was subsequently used for hierarchical clustering (Figure 22). List of proximal SRY sequences based on proximity of 4-grams hydropathy profiles was summarized in Table 26.

**Figure 22:**
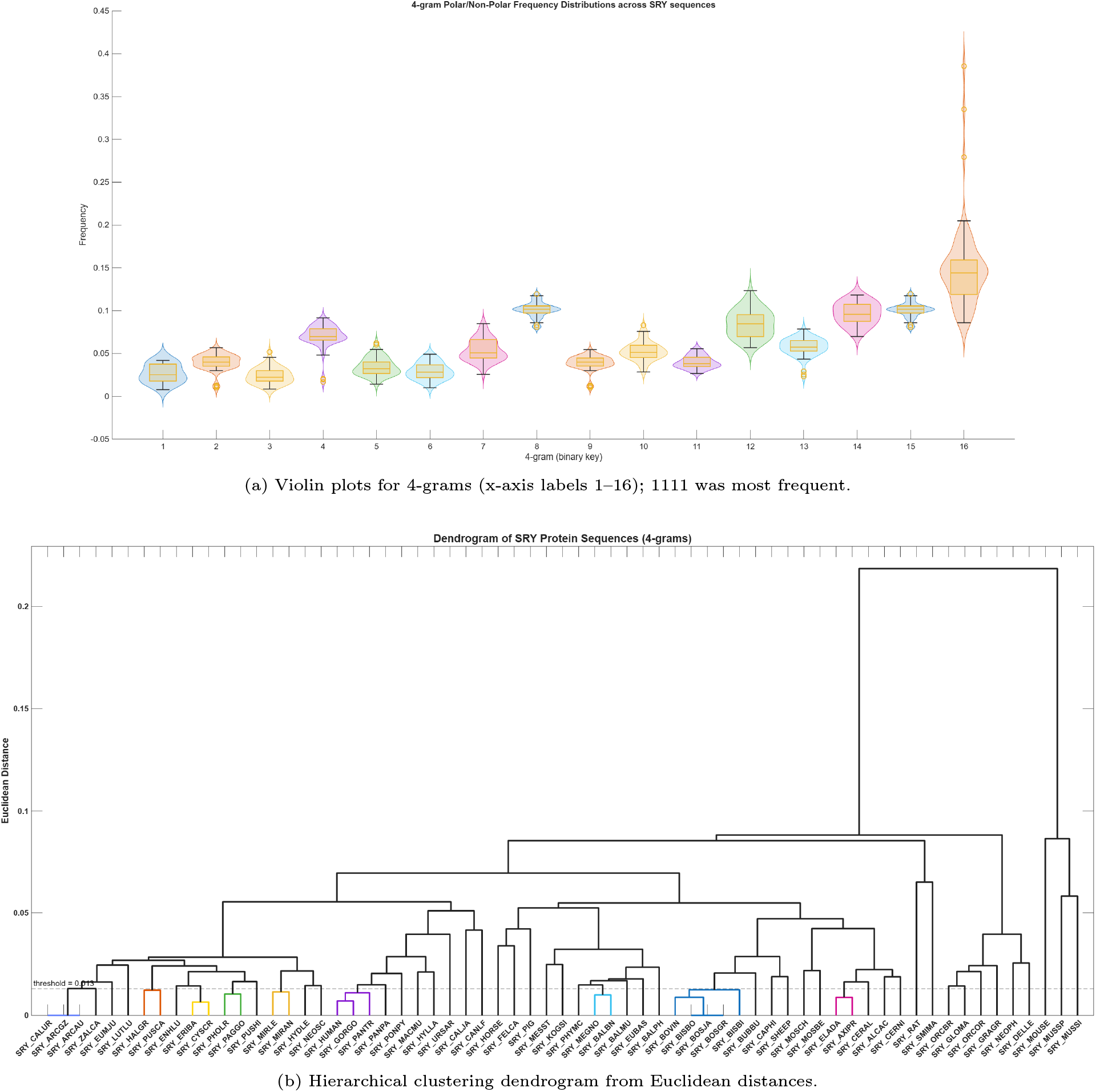
Hydropathy-based 4-gram frequency profiles of 65 SRY sequences.

**Table 26:**
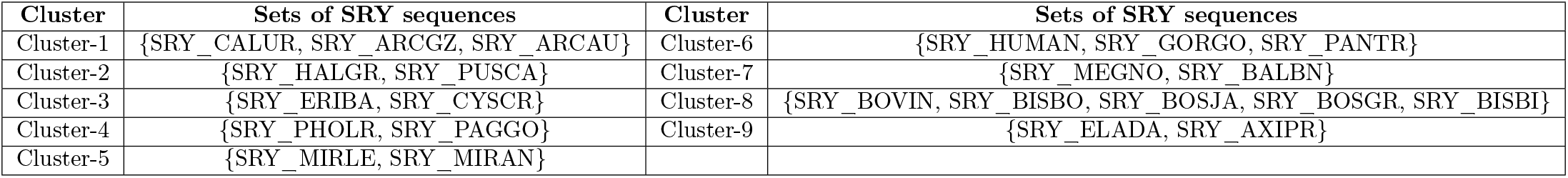
List of proximal SRY sequences based on proximity of 4-grams hydropathy profiles.

#### 4.8.2. Analysis of hydropathy-based n-gram frequency across SRY-HMG subsequences

##### Relative Frequency of 2-grams and associated phylogenetic relationships of SRY-HMG subsequences

The 38 SRY-HMG subsequences were converted into binary hydropathy strings and read with a sliding window of two residues. What stood out when comparing these SRY-HMG subsequences with the SRY protein sequences was how much more even the hydropathy balance became. The polar–polar pair 11 was still the single most common pattern, but it was noticeably less dominant here, averaging 0.3609 and spanning only 0.3382 to 0.4179 (Figure 23). The nonpolar–nonpolar pair 00 moved up into second place with a mean of 0.1898, while the two transition patterns 01 and 10 sat close together at means of 0.2320 and 0.2173 (Figure 23). In other words, the HMG box carried more consecutive nonpolar residues and fewer runs of polar residues than the full-length SRY sequence, which fitted with the structural role this domain was known to play.

**Figure 23:**
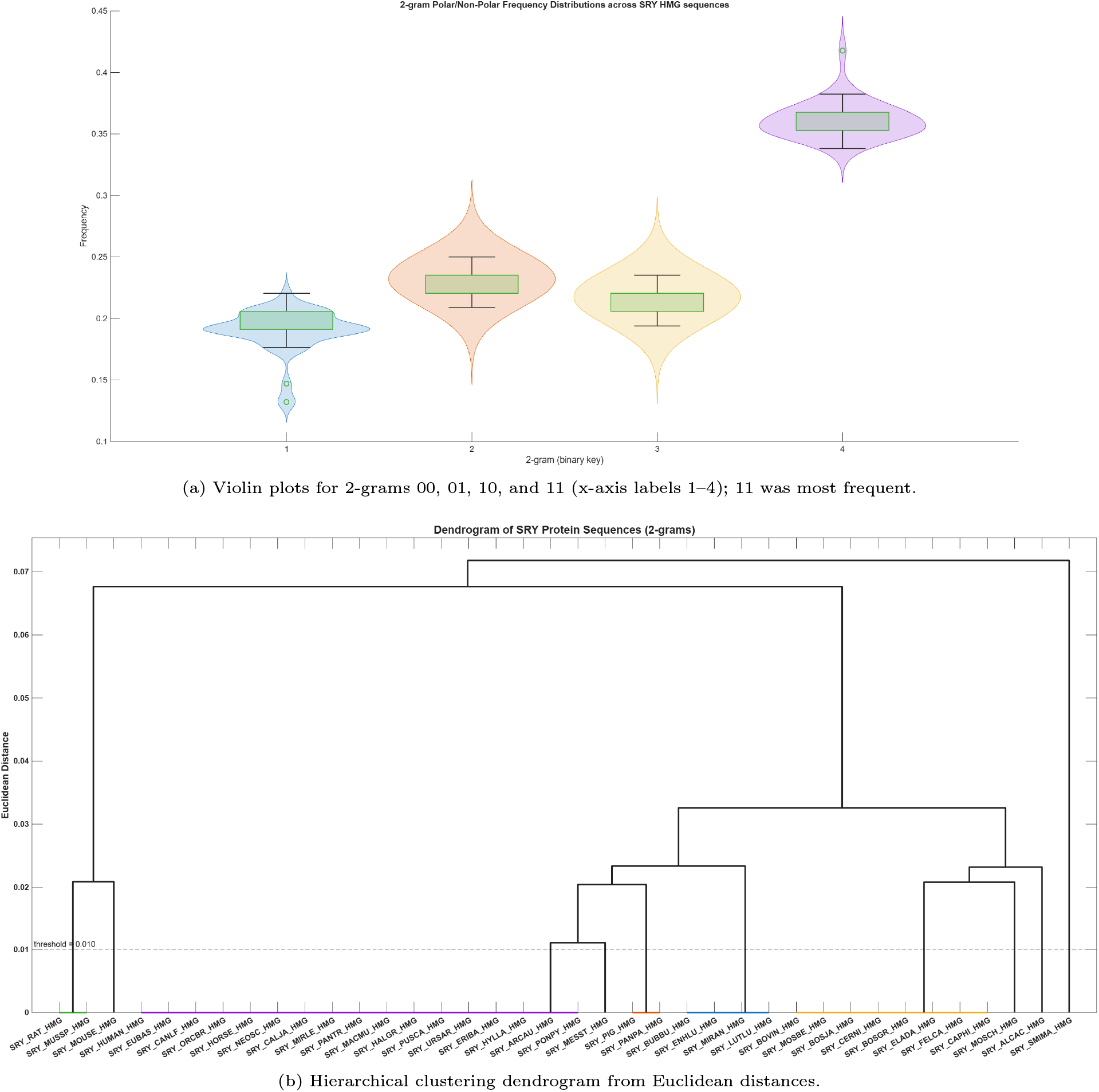
Hydropathy-based 2-gram frequency profiles of 38 SRY-HMG sequences.

Perhaps the most important feature of these SRY-HMG subsequences was how uniform the subsequences were. Many of them shared exactly the same 2-gram profile. Seventeen subsequences, including SRY_HUMAN_HMG, SRY_HORSE_HMG, and SRY_CALJA_HMG, all showed the identical profile (0.1912, 0.2353, 0.2206, 0.3529) (Table 27). Another eight, among them SRY_BOVIN_HMG and SRY_CAPHI_HMG, shared the profile (0.2059, 0.2206, 0.2059, 0.3676). The remaining SRY-HMG sequences were only slightly different. The lowest 00 value (0.1324) belonged to SRY_RAT_HMG and SRY_MUSSP_HMG, while SRY_MOSCH_HMG held the highest (0.2206). SRY_SMIMA_HMG was the outlier at the other end: it had the lowest 01 and 10 frequencies (0.2090 and 0.1940) but the highest 11 frequency (0.4179). SRY_RAT_HMG, SRY_MOUSE_HMG, and SRY_MUSSP_HMG together reached the maximum 01 and 10 values (0.2500 and 0.2353), and SRY_PIG_HMG together with SRY_PANPA_HMG showed the lowest 11 frequency (0.3382) (Table 27).

**Table 27:** Summary of hydropathy-based 2-gram relative frequencies across 38 SRY-HMG subsequences.

| 2-grams | Hydropathy pattern | Mean | Range | Min sequence(s) | Max sequence(s) |
| --- | --- | --- | --- | --- | --- |
| 00 | Nonpolar-nonpolar | 0.1898 | 0.1324-0.2206 | SRY_RAT_HMG, SRY_MUSSP_HMG | SRY_MOSCH_HMG |
| 01 | Nonpolar-polar | 0.2320 | 0.2090-0.2500 | SRY_SMIMA_HMG | SRY_RAT_HMG, SRY_MOUSE_HMG, SRY_MUSSP_HMG |
| 10 | Polar-nonpolar | 0.2173 | 0.1940-0.2353 | SRY_SMIMA_HMG | SRY_RAT_HMG, SRY_MOUSE_HMG, SRY_MUSSP_HMG |
| 11 | Polar-polar | 0.3609 | 0.3382-0.4179 | SRY_PIG_HMG, SRY_PANPA_HMG | SRY_SMIMA_HMG |

These four relative frequencies formed the feature vector for each subsequence and were used to compute pairwise Euclidean distances, producing the 38 *×* 38 distance matrix that was later subjected to hierarchical clustering (Figure 23). List of proximal SRY-HMG sequences based on proximity of 2-grams hydropathy profiles was summarized in Table 28.

**Table 28:** List of proximal SRY-HMG sequences based on proximity of 2-grams hydropathy profiles.

| Cluster | Sets of SRY-HMG subsequences |
| --- | --- |
| Cluster-1 | {SRY_RAT_HMG, SRY_MUSSP_HMG} |
| Cluster-2 | {SRY_HUMAN_HMG, SRY_EUBAS_HMG, SRY_CANLF_HMG, SRY_ORCBR_HMG, SRY_HORSE_HMG, SRY_NEOSC_HMG, SRY_CALJA_HMG, SRY_MIRLE_HMG, SRY_PANTR_HMG, SRY_MACMU_HMG, SRY_HALGR_HMG, SRY_PUSCA_HMG, SRY_URSAR_HMG, SRY_ERIBA_HMG, SRY_HYLLA_HMG, SRY_ARCAU_HMG, SRY_PONPY_HMG} |
| Cluster-3 | {SRY_PIG_HMG, SRY_PANPA_HMG} |
| Cluster-4 | {SRY_BUBBU_HMG, SRY_ENHLU_HMG, SRY_MIRAN_HMG, SRY_LUTLU_HMG} |
| Cluster-5 | {SRY_BOVIN_HMG, SRY_MOSBI_HMG, SRY_BOSJA_HMG, SRY_CERNI_HMG, SRY_BOSGR_HMG, SRY_ELADA_HMG, SRY_FELCA_HMG, SRY_CAPHI_HMG} |

##### Relative Frequency of 3-grams and associated phylogenetic relationships of SRY-HMG subsequences

The binary hydropathy strings of the 38 SRY-HMG subsequences were scanned with a sliding window of three residues, and the relative frequencies of the eight possible 3-grams were calculated (Figure 24). A different picture emerged here compared with the full-length SRY protein sequences. In the SRY-HMG box, the most abundant 3-gram was not the fully polar 111, but the pattern 011 (nonpolar–polar–polar), with a mean of 0.1847 and a range of 0.1515–0.2239. The fully polar 111 followed closely with a mean of 0.1816 (range 0.1642–0.2727), while 110 (polar–polar–nonpolar) averaged 0.1702 (range 0.1364–0.2090). The two transition patterns 101 and 001/100 occupied the middle of the distribution, averaging 0.1164, 0.1042, and 0.1042, respectively. At the bottom end, 000 (nonpolar–nonpolar–nonpolar) was the rarest word (mean 0.0885), followed by 010 (nonpolar–polar–nonpolar; mean 0.0503). The comparatively high frequencies of 011 and 110, together with the reduced 111 frequency, indicated that the HMG box mixed polar and nonpolar residues more evenly than the full-length SRY sequence.

**Figure 24:**
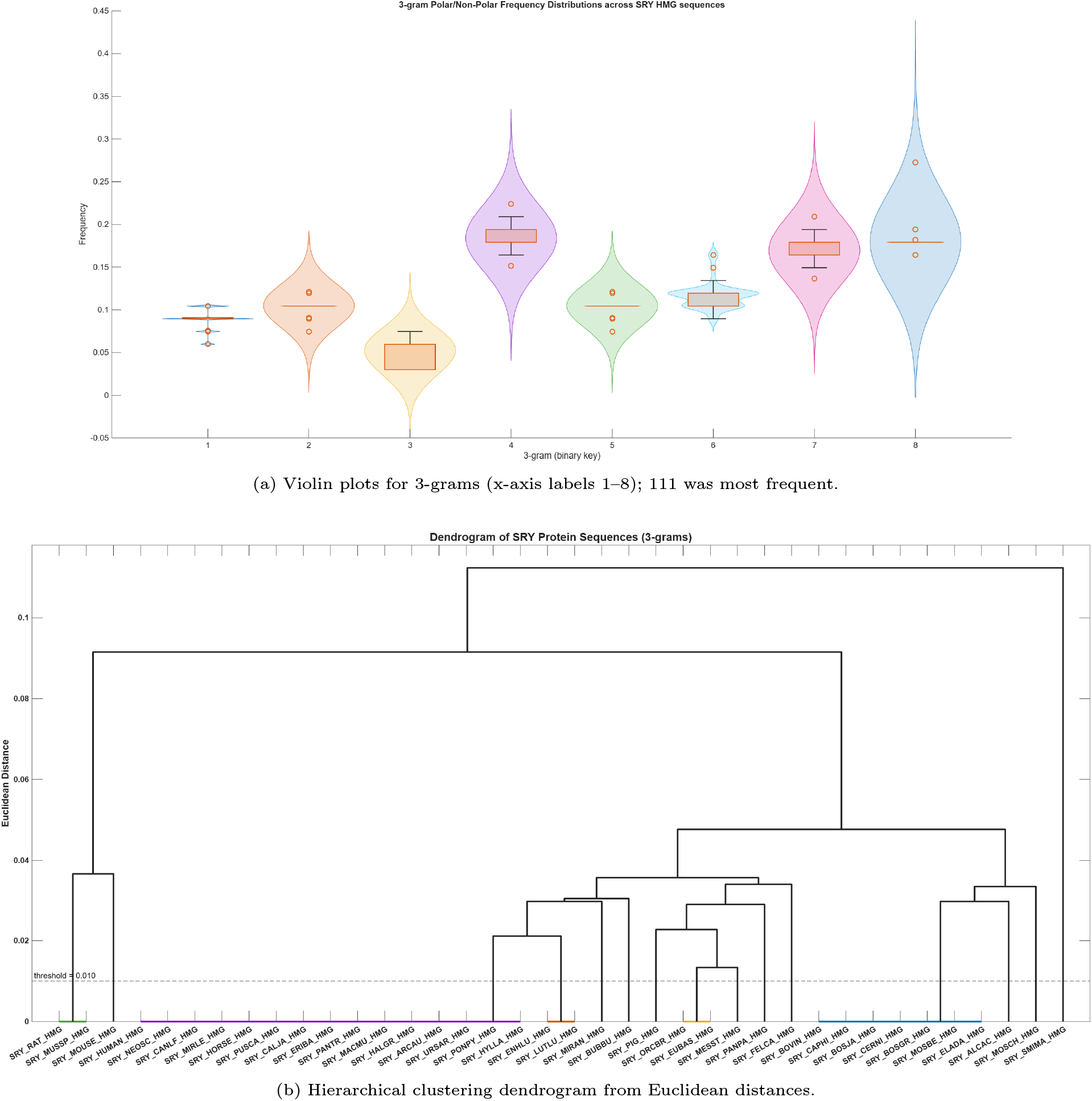
Hydropathy-based 3-gram frequency profiles of 38 SRY-HMG sequences.

A single 3-gram profile, (0.0896, 0.1045, 0.0597, 0.1791, 0.1045, 0.1194, 0.1642, 0.1791), was shared by fifteen subsequences, among them SRY_HUMAN_HMG, SRY_HORSE_HMG, SRY_CALJA_HMG, and SRY_CANLF_HMG. A second profile, (0.1045, 0.1045, 0.0299, 0.1940, 0.1045, 0.1045, 0.1791, 0.1791), was shared by seven subsequences, including SRY_BOVIN_HMG, SRY_CAPHI_HMG, and SRY_BOSGR_HMG (Table 29). Only a couple of subsequences stood apart. SRY_SMIMA_HMG was the clear outlier, combining the lowest 011 frequency (0.1515) with the highest 111 frequency (0.2727) and the lowest 110 frequency (0.1364), which pointed to a more polar character than the rest of the SRY-HMG subsequences. SRY_PANPA_HMG had the highest 010 frequency (0.0746), while SRY_MESST_HMG showed the highest 001 and 100 frequencies (0.1212 each). At the other end, SRY_RAT_HMG and SRY_MUSSP_HMG shared the lowest 000 (0.0597), 001/100 (0.0746), and 010 (0.0299) frequencies, and they also reached the maximum 011, 101, and 110 values (0.2239, 0.1642, and 0.2090, respectively) — the mirror image of the pattern seen in SRY_SMIMA_HMG (Table 29).

**Table 29:** Summary of hydropathy-based 3-gram relative frequencies across 38 SRY-HMG subsequences.

| 3-grams | Hydropathy pattern | Mean | Range | Min sequence(s) | Max sequence(s) |
| --- | --- | --- | --- | --- | --- |
| 000 | Nonpolar–nonpolar–nonpolar | 0.0885 | 0.0597–0.1045 | SRY_RAT_HMG,<br>SRY_MOUSE_HMG,<br>SRY_MUSSP_HMG | SRY_BOVIN_HMG,<br>SRY_CAPHI_HMG,<br>SRY_MOSCH_HMG,<br>SRY_MOSBE_HMG,<br>SRY_ELADA_HMG,<br>SRY_ALCAC_HMG,<br>SRY_BOSJA_HMG,<br>SRY_CERNI_HMG,<br>SRY_BOSGR_HMG |
| 001 | Nonpolar–nonpolar–polar | 0.1042 | 0.0746–0.1212 | SRY_RAT_HMG,<br>SRY_MUSSP_HMG | SRY_MESST_HMG |
| 010 | Nonpolar–polar–nonpolar | 0.0503 | 0.0299–0.0746 | SRY_RAT_HMG,<br>SRY_BOVIN_HMG,<br>SRY_CAPHI_HMG,<br>SRY_MUSSP_HMG,<br>SRY_MOSCH_HMG,<br>SRY_MOSBE_HMG,<br>SRY_ELADA_HMG,<br>SRY_ALCAC_HMG,<br>SRY_BOSJA_HMG,<br>SRY_CERNI_HMG,<br>SRY_BOSGR_HMG | SRY_PANPA_HMG |
| 011 | Nonpolar–polar–polar | 0.1847 | 0.1515–0.2239 | SRY_SMIMA_HMG | SRY_RAT_HMG,<br>SRY_MUSSP_HMG |
| 100 | Polar–nonpolar–nonpolar | 0.1042 | 0.0746–0.1212 | SRY_RAT_HMG,<br>SRY_MUSSP_HMG | SRY_MESST_HMG |
| 101 | Polar–nonpolar–polar | 0.1164 | 0.0896–0.1642 | SRY_MOSCH_HMG,<br>SRY_FELCA_HMG | SRY_RAT_HMG,<br>SRY_MUSSP_HMG |
| 110 | Polar–polar–nonpolar | 0.1702 | 0.1364–0.2090 | SRY_SMIMA_HMG | SRY_RAT_HMG,<br>SRY_MUSSP_HMG |
| 111 | Polar–polar–polar | 0.1816 | 0.1642–0.2727 | SRY_RAT_HMG,<br>SRY_MOUSE_HMG,<br>SRY_PIG_HMG,<br>SRY_MUSSP_HMG,<br>SRY_MOSCH_HMG | SRY_SMIMA_HMG |

Furthermore, it was observed that the normalized frequency of 001 and 100 were identical for every subsequence, the same near-symmetry noted earlier for the 2-gram 01/10 pair. These eight relative frequencies formed the feature vector for each subsequence and were used to compute pairwise Euclidean distances, yielding the 38 *×* 38 distance matrix that was subsequently subjected to hierarchical clustering (Figure 24). List of proximal SRY-HMG sequences based on proximity of 3-grams hydropathy profiles was summarized in Table 30.

**Table 30:** List of proximal SRY-HMG sequences based on proximity of 3-grams hydropathy profiles.

| Cluster | Sets of SRY-HMG subsequences |
| --- | --- |
| Cluster-1 | {SRY_RAT_HMG, SRY_MUSSP_HMG} |
| Cluster-2 | {SRY_HUMAN_HMG, SRY_NEOSC_HMG, SRY_CANLF_HMG, SRY_MIRLE_HMG, SRY_HORSE_HMG, SRY_PUSCA_HMG, SRY_CALJA_HMG, SRY_ERIBA_HMG, SRY_PANTR_HMG, SRY_MACMU_HMG, SRY_HALGR_HMG, SRY_ARCAU_HMG, SRY_URSAR_HMG, SRY_PONPY_HMG, SRY_HYLLA} |
| Cluster-3 | {SRY_ENHLU_HMG, SRY_LUTLU_HMG} |
| Cluster-4 | {SRY_ORCBR_HMG, SRY_EUBAS_HMG} |
| Cluster-5 | {SRY_BOVIN_HMG, SRY_CAPHI_HMG, SRY_BOSJA_HMG, SRY_CERNI_HMG, SRY_BOSGR_HMG, SRY_MOSBE_HMG, SRY_ELADA_HMG} |

##### Relative Frequency of 4-grams and associated phylogenetic relationships of SRY-HMG subsequences

The 38 SRY-HMG subsequences were converted into binary hydropathy strings and scanned with a sliding window of four residues. Every overlapping quartet was classified according to the polar/nonpolar status of its four positions, and the relative frequency of each of the sixteen possible 4-grams was recorded. The most abundant 4-gram across the SRY-HMG subsequences was 1111 (polar–polar–polar–polar), with a mean relative frequency of 0.1058 (range 0.0606–0.1846). It was followed closely by 0110 (nonpolar–polar–polar–nonpolar; mean 0.1038) and 1101 (polar–polar–nonpolar–polar; mean 0.1022) (Figure 25). The patterns 0011 and 1011 occupied the next tier, averaging 0.0890 and 0.0834, respectively, while 0111 and 1100 had means of 0.0786 and 0.0706. At the other end of the spectrum, 0101 (nonpolar–polar–nonpolar–polar) was the rarest word (mean 0.0160), just below 0010 (mean 0.0164) (Table 31). Compared with the full-length SRY protein sequences, the SRY-HMG box showed a more even spread of 4-gram frequencies, with no single pattern dominating as strongly as 1111 had done in the full-length sequences.

**Figure 25:**
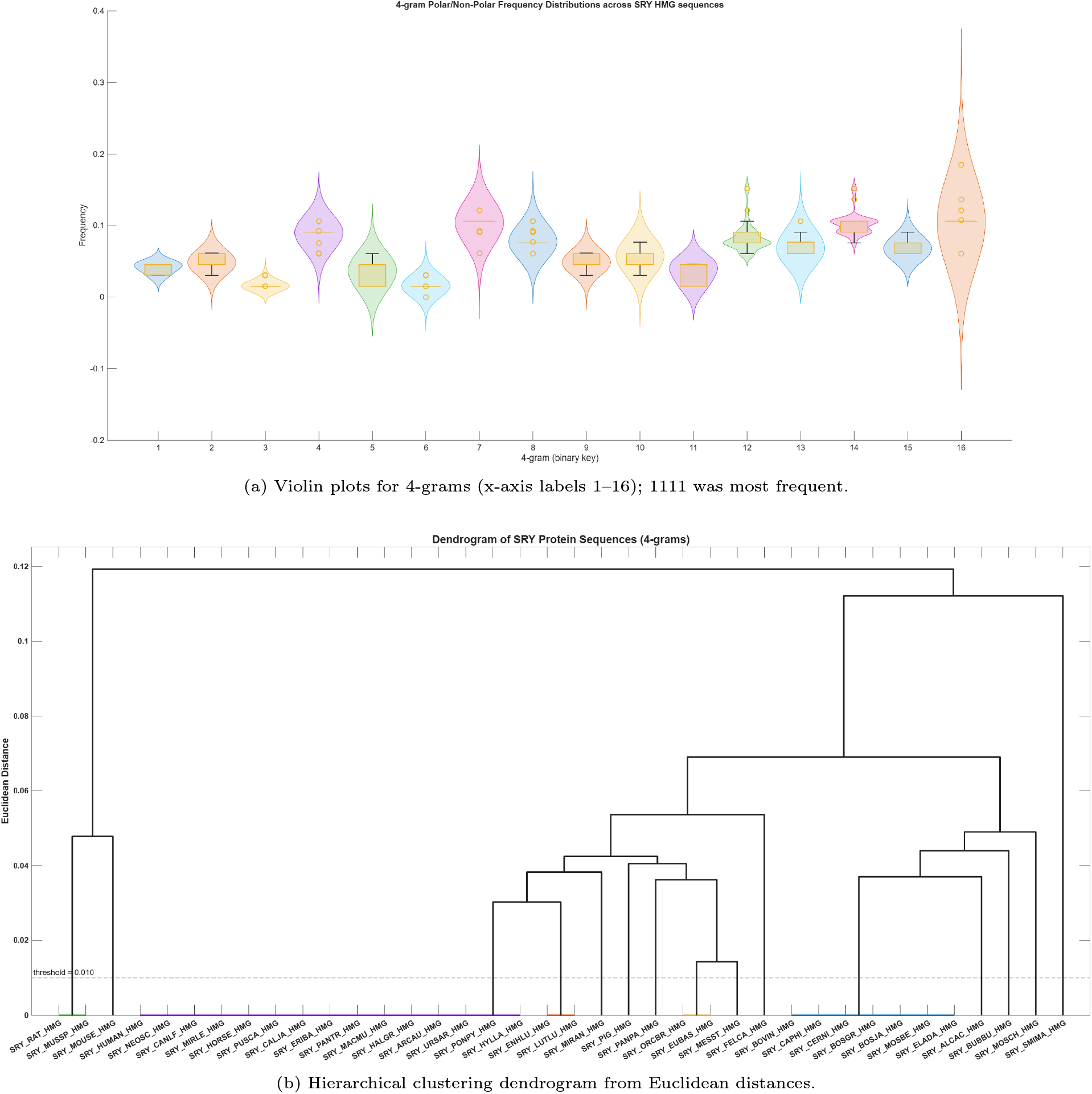
Hydropathy-based 4-gram frequency profiles of 38 SRY-HMG sequences.

**Table 31:**
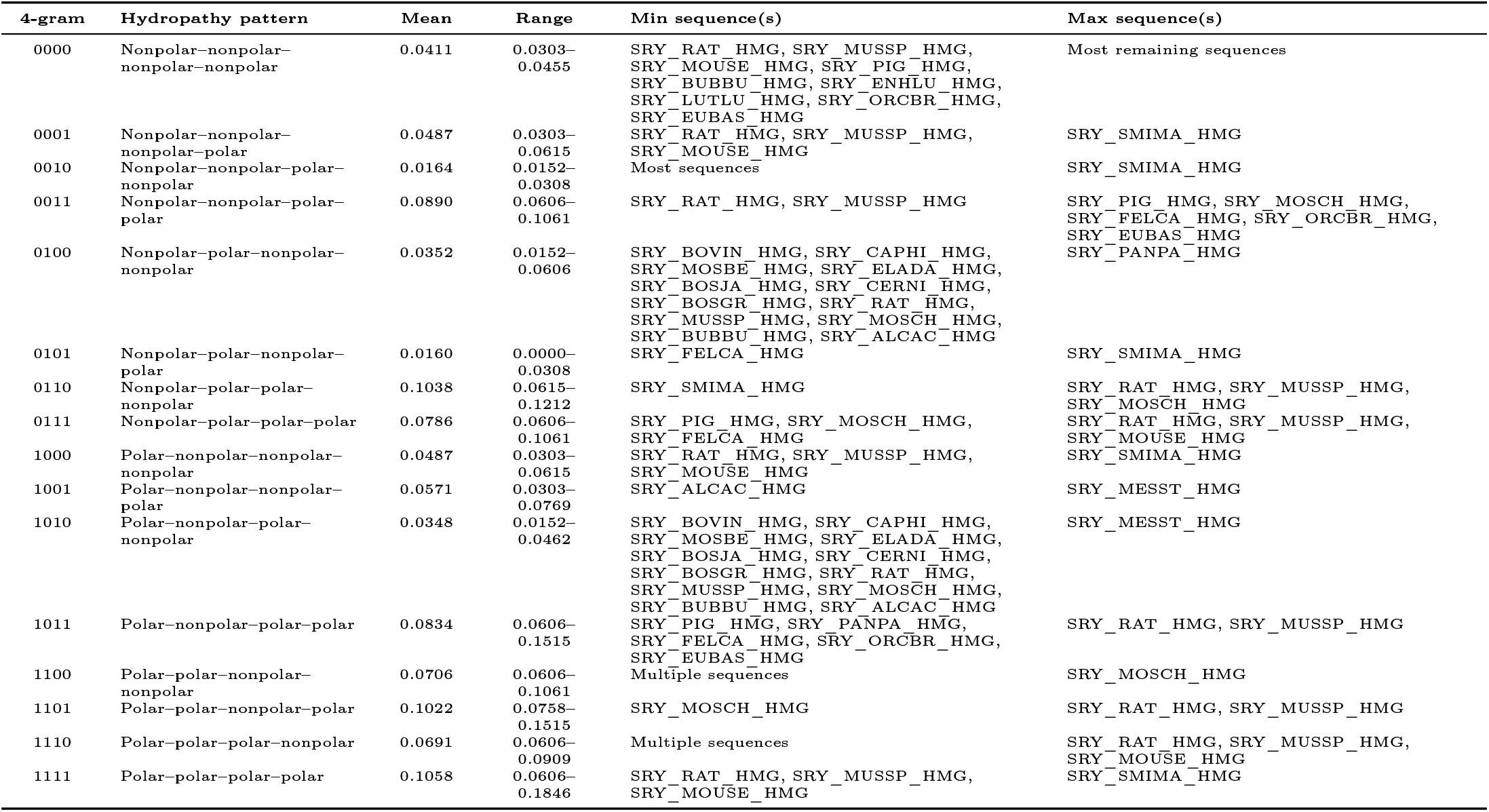
Summary of hydropathy-based 4-gram relative frequencies across 38 SRY-HMG subsequences.

Fifteen of SRY-HMG subsequences, including SRY_HUMAN_HMG, SRY_HORSE_HMG, and SRY_CALJA_HMG, shared an identical 4-gram profile. Seven others, among them SRY_BOVIN_HMG, and SRY_CAPHI_HMG, formed a second identical group. SRY_RAT_HMG and SRY_MUSSP_HMG were indistinguishable from each other, as were SRY_ENHLU_HMG and SRY_LUTLU_HMG, and SRY_ORCBR_HMG and SRY_EUBAS_HMG. Only a few subsequences stood out as distinct. SRY_SMIMA_HMG was the clearest outlier: it had the highest 1111 frequency (0.1846), the highest 0010 frequency (0.0308), and the highest 0101 frequency (0.0308), while also showing the lowest 0110 frequency (0.0615). SRY_FELCA_HMG was unusual in having a 0101 frequency of exactly 0. SRY_MOSCH_HMG reached the maximum 1100 frequency (0.1061) and the minimum 1101 frequency (0.0758) (Table 31). SRY_MESST_HMG held the highest 1001 and 1010 frequencies (0.0769 and 0.0462). SRY_PANPA_HMG had the highest 0100 frequency (0.0606), and SRY_ALCAC_HMG the lowest 1001 frequency (0.0303). SRY_MOUSE_HMG shared the lowest 0001, 1000, and 1111 frequencies with SRY_RAT_HMG and SRY_MUSSP_HMG, while SRY_RAT_HMG and SRY_MUSSP_HMG together reached the maximum 1011 and 1101 frequencies (0.1515 each) (Table 31).

These sixteen relative frequencies formed the feature vector for each subsequence and were used to compute pairwise Euclidean distances, producing the 38 *×* 38 distance matrix that was subsequently subjected to hierarchical clustering (Figure 25). List of proximal SRY-HMG sequences based on proximity of 4-grams hydropathy profiles was summarized in Table 32.

**Table 32:** List of proximal SRY-HMG sequences based on proximity of 4-grams hydropathy profiles.

| Cluster | Sets of SRY-HMG subsequences |
| --- | --- |
| Cluster-1 | {SRY_RAT_HMG, SRY_MUSSP_HMG} |
| Cluster-2 | {SRY_HUMAN_HMG, SRY_NEOSC_HMG, SRY_CANLF_HMG, SRY_MIRLE_HMG, SRY_HORSE_HMG, SRY_PUSCA_HMG, SRY_CALJA_HMG, SRY_ERIBA_HMG, SRY_PANTR_HMG, SRY_MACMU_HMG, SRY_HAMLGR_HMG, SRY_ARCAU_HMG, SRY_URSAR_HMG, SRY_PONPY_HMG, SRY_HYLLA_HMG} |
| Cluster-3 | {SRY_ENHLU_HMG, SRY_LUTLU_HMG} |
| Cluster-4 | {SRY_ORCBR_HMG, SRY_EUBAS_HMG} |
| Cluster-5 | {SRY_BOVIN_HMG, SRY_CAPHI_HMG, SRY_CERNI_HMG, SRY_BOSGR_HMG, SRY_BOSJA_HMG, SRY_MOSBE_HMG, SRY_ELADA_HMG} |

### 4.9. Hydropathy profile-based clustering of SRY and SRY-HMG sequences

The percentages of polar and nonpolar residues were computed for all 65 SRY sequences and all 38 SRY-HMG subsequences. In both sets, polar residues outnumbered nonpolar residues, which was expected given that the polar set contains eleven amino acids and the nonpolar set nine. However, the two sets of SRY sequences and SRY-HMG subsequences differed markedly in both the level and the spread of their polar content.

For the SRY sequences, polar residues made up an average of 61.96% (median 61.64%), with values spanning 57.21% to 76.28%. Nonpolar residues averaged 38.04%, ranging from 23.72% to 42.79%. The distribution was wide, and the extremes were lineage-specific (Figure 26). The highest polar content occurred in SRY_MOUSE (76.28%), SRY_MUSSP (74.93%), and SRY_MUSSI (72.35%), all rodents. The lowest values were shared by SRY_ORCBR and SRY_DELLE (57.21%), followed by SRY_GLOMA (57.71%), SRY_PIG (58.05%), SRY_NEOPH (58.16%), SRY_GRAGR (58.21%), and SRY_PUSCA (58.62%), a group dominated by cetaceans. Fifty of the 65 sequences fell within a narrower band between 59% and 64% (Figure 26).

**Figure 26:**
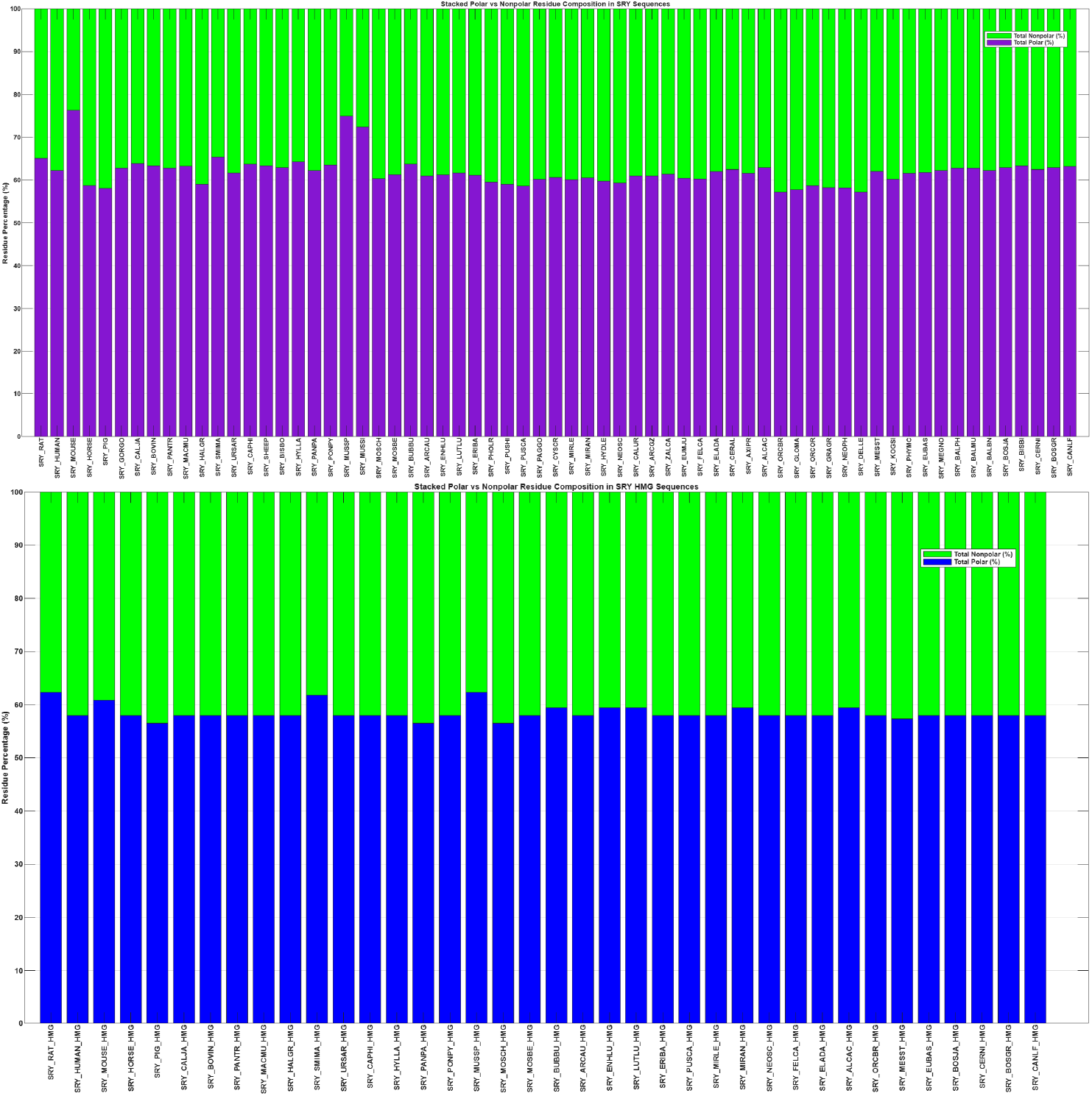
Relative percentage of polar and non-polar residues in SRY and SRY-HMG sequences.

For the SRY-HMG subsequences, the picture was a bit different. Polar residues averaged 58.44%, with a range of only 56.52% to 62.32%. Nonpolar residues averaged 41.56%, ranging from 37.68% to 43.48%. The most notable feature was the uniformity: 25 of the 38 subsequences (65.8%) had exactly 57.97% polar residues. The remaining sequences deviated only slightly (Figure 26). Three subsequences (SRY_PIG_HMG, SRY_PANPA_HMG, and SRY_MOSCH_HMG) had 56.52% polar residues; five (SRY_BUBBU_HMG, SRY_ENHLU_HMG, SRY_LUTLU_HMG, SRY_MIRAN_HMG, and SRY_ALCAC_HMG) had 59.42%; one (SRY_MESST_HMG) had 57.35%; one (SRY_MOUSE_HMG) had 60.87%; one (SRY_SMIMA_HMG) had 61.76%; and two (SRY_RAT_HMG and SRY_MUSSP_HMG) had 62.32%. The maximum polar content within the HMG set was therefore 62.32%, well below the 76.28% seen in SRY_MOUSE (Figure 26).

Polar and nonpolar content among SRY and SRY-HMG sequences revealed a clear contrast. The SRY sequences showed a higher mean polar content (61.96% vs 58.44%) and a much wider range (19.07 percentage points vs 5.80 percentage points). The HMG subsequences, by contrast, were tightly constrained, with nearly two-thirds sharing an identical polar fraction. This difference is consistent with the known biology: the HMG box must maintain a specific fold for DNA binding, and it appears to be under stronger purifying selection than the rest of the protein [52, 62]. The lineage-specific variation seen in SRY—high polar content in rodents, low in cetaceans—was largely absent from the HMG domain [63]. In other words, the compositional freedom of the protein is not mirrored in its DNA-binding domain, which remains close to a fixed polar/nonpolar balance across organisms.

#### 4.9.1. Clustering based on the longest hydropathy stretches across SRY and SRY-HMG sequences

The longest uninterrupted run of polar residues and the longest uninterrupted run of nonpolar residues were located in each of the 65 SRY sequences and each of the 38 SRY-HMG subsequences. The absolute lengths were divided by the total sequence length and expressed as percentages, so that sequences of different sizes could be compared on the same scale. The longest polar and nonpolar stretches for SRY_HUMAN and SRY_HUMAN_HMG are shown in Figure 27. These two normalized values—longest polar stretch percentage and longest nonpolar stretch percentage—formed a two-dimensional feature vector for every SRY/SRY-HMG sequence.

**Figure 27:**
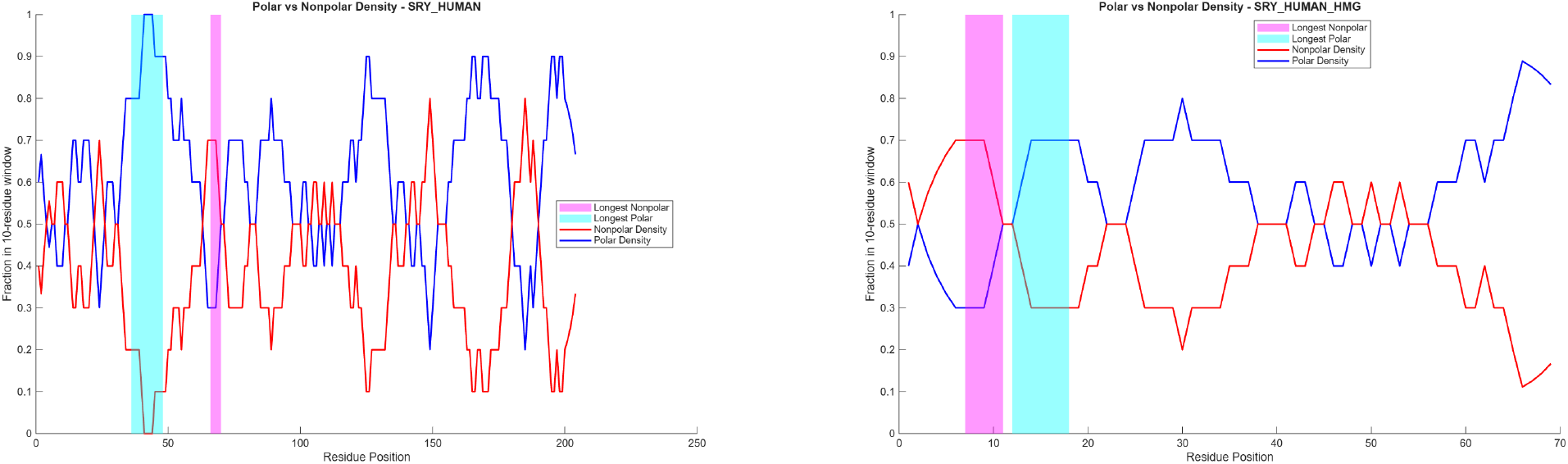
Density of polar and non-polar residues and the longest polar and non-polar stretches of SRY_HUMAN and SRY_HUMAN_HMG.

For SRY protein sequences, the longest polar stretch averaged 4.22% of the sequence (range 2.92%–6.51%), while the longest nonpolar stretch averaged 2.32% (range 1.28%–3.98%). The two measures were therefore not of equal magnitude: polar runs were consistently longer than nonpolar runs, mirroring the overall polar bias of the protein. SRY_RAT carried the longest polar stretch (6.51%), followed by SRY_HUMAN, SRY_GORGO, SRY_PANTR, and SRY_PANPA, all at 6.37% (Figure 28). At the other end, SRY_CAPHI and SRY_SHEEP shared the shortest polar stretch (2.92%). For nonpolar runs, SRY_ORCBR was the clear outlier (3.98%), well above the next highest values of 2.99% in SRY_GLOMA and SRY_ORCOR. SRY_MOUSE had the shortest nonpolar stretch (1.28%), followed by SRY_MUSSP (1.69%). The values spanned a wide range across the 65 SRY sequences, reflecting lineage-specific differences in long-range hydropathy architecture (Figure 28).

**Figure 28:**
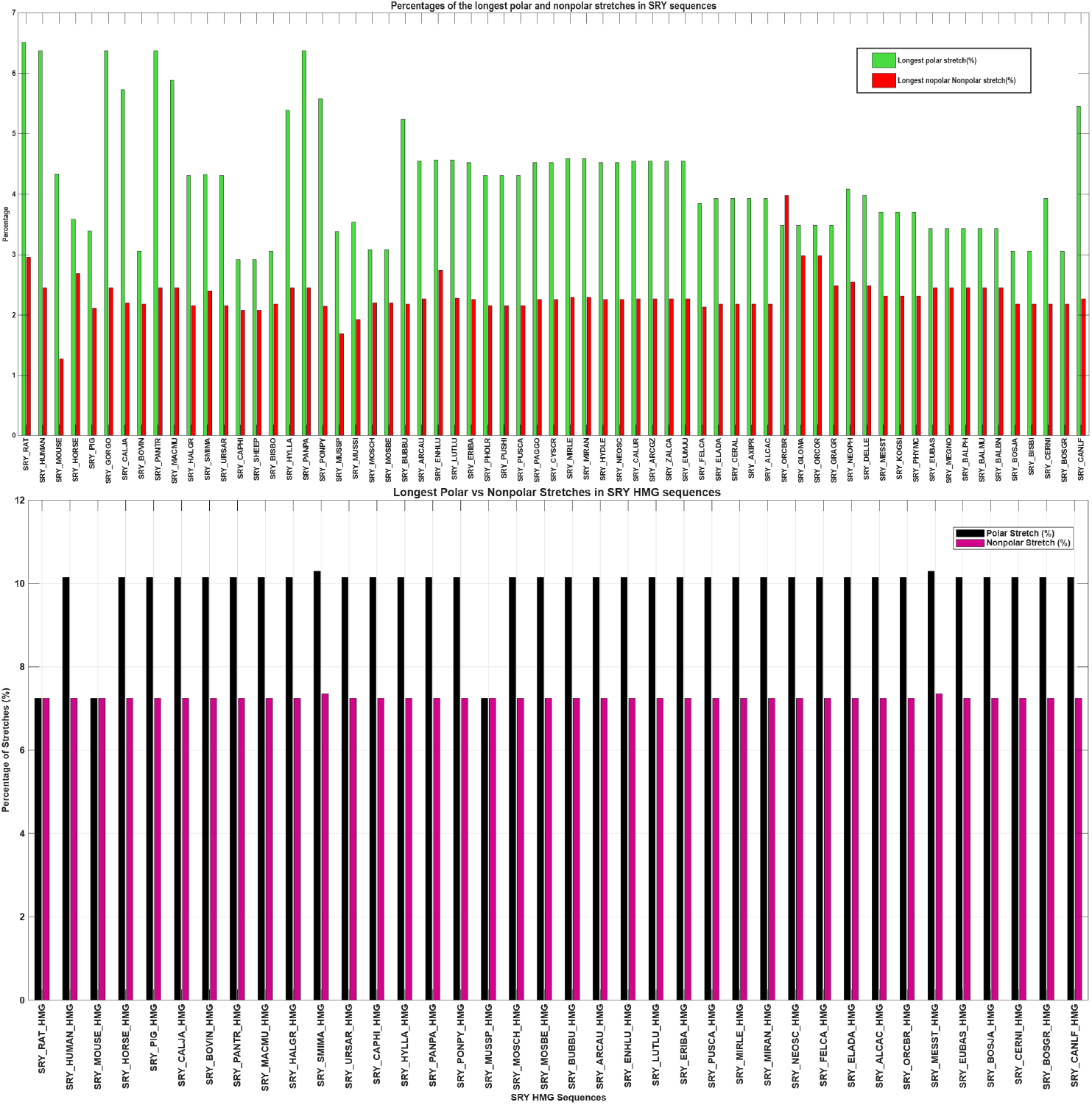
Relative percentage of the longest polar and non-polar stretches in SRY and SRY-HMG sequences.

The SRY-HMG subsequences were organized loudly in form the longest polar and nonpolar residue stretches. Here, the longest polar stretch averaged 9.92% and the longest nonpolar stretch averaged 7.25%. These higher percentages simply reflect the shorter length of the HMG subsequences, not a change in composition. What mattered was the uniformity: 33 of the 38 subsequences (86.8%) had exactly the same values, with a polar stretch of 10.14% and a nonpolar stretch of 7.25% (Figure 28). Three subsequences—SRY_RAT_HMG, SRY_MOUSE_HMG, and SRY_MUSSP_HMG—had a shorter polar stretch of 7.25% but the same nonpolar stretch of 7.25%. Only two, SRY_SMIMA_HMG and SRY_MESST_HMG, deviated slightly upward, with a polar stretch of 10.29% and a nonpolar stretch of 7.35% (Figure 28).

The SRY sequences varied widely in both measures, with polar stretches ranging over 3.59 percentage points and nonpolar stretches over 2.70 percentage points. The SRY-HMG subsequences, by contrast, were almost invariant, with polar stretches spanning only 3.05 percentage points and nonpolar stretches only 0.10 percentage points. The three rodent SRY-HMG subsequences were the only ones with a noticeably shorter polar stretch, but even they retained the same nonpolar stretch as the majority. SRY_SMIMA_HMG and SRY_MESST_HMG differed only marginally. Accordingly, the long-range hydropathy architecture of the SRY-HMG domain was far more homogeneous than that of the full-length SRY protein sequences, consistent with the structural constraints imposed by its DNA-binding function.

These normalized longest-stretch values were used as two-dimensional feature vectors for density-based clustering with DBSCAN with specified *ɛ* = 0.05 and MinPts=3. This analysis yielded six distinct clusters of SRY sequences and two clusters of SRY-HMG subsequences, as presented in Table 33). Furthermore, SRY sequences belonging to the outlier cluster were identified where the longest polar or non-polar stretches deviated substantially from cluster centroids, indicating atypical hydropathy domain architecture.

**Table 33:**
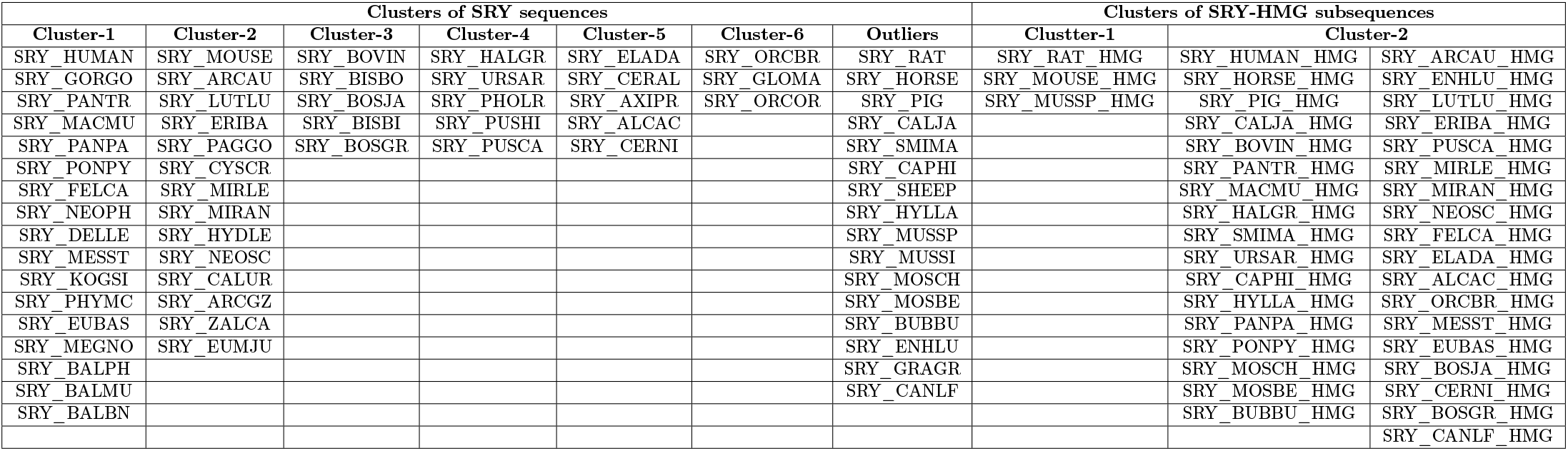
Clustering based on the longest polar and non-polar stretches across SRY and SRY-HMG sequences.

### 4.10. Intrinsic disorder profiles analysis of SRY and SRY-HMG sequences

Figure 29 represented the result of the global disorder analysis of 65 unique SRY proteins and 38 unique HMG domains. As per Figure XA, all SRY proteins were predicted as highly disordered: their PONDR® VSL2-based percent of predicted intrinsically disordered residues (PPIDR) ranged from 94.61% for SRY_HYLLA to 51.27% for SRY_PIG; whereas their average disorder scores (ADS) ranged from 0.7931 for SRI-MOUSE to 0.5119 for SRY_AXIPR. Figure XA also illustrated that HMG domains were expected to contain high levels of intrinsic disorder as well. In fact, only one domain (SRY_BOSGR_HMG) was characterized as moderately disordered (it was positioned in the pink area marked M/M), whereas 6 HMG domains (SRY_PIG_HMG, SRY_CAPHI_HMG, SRY_MOSBE_HMG, SRY_FELCA_HMG, SRY_CERNI_HMG, and SRY_BOVIN_HMG) belonged to the moderately disordered plus category (were located within the light pink area marked “mixed”). The remaining 31 HMG domains were predicted as highly disordered with PPIDR ranging from 43.48% (SRY_ELADA_HMG) to 98.55% (SRY-RAT-HMG) and ADS ranging from 0.5071 (SRY_ELADA_HMG) to 0.6658 (SRY-RAT-HMG).

**Figure 29:**
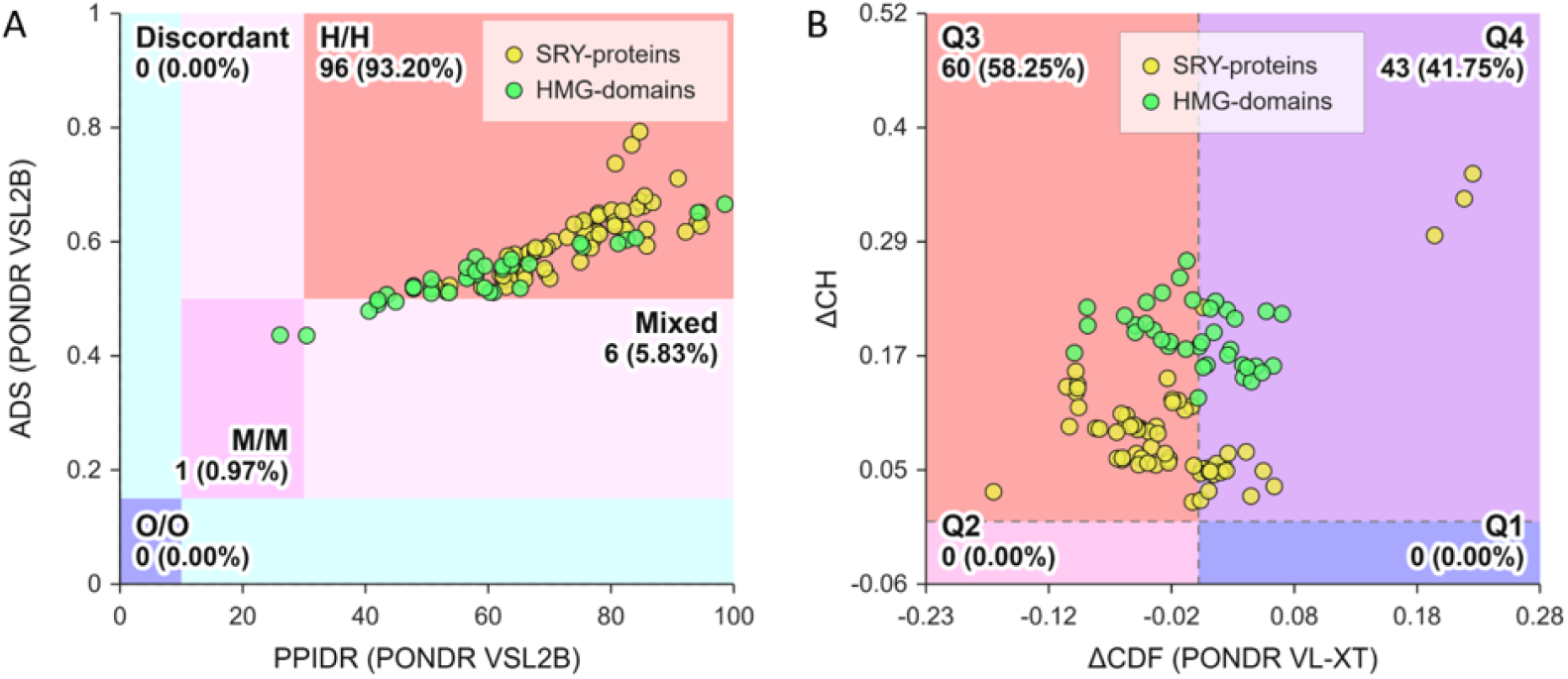
Global disorder analysis of unique SRY proteins (yellow circles) and unique HMG domains (green circles). A. A scatter plot compared the average disorder score (ADS) against the percentage of predicted intrinsically disordered residues (PPIDR) based on PONDR® VSL2 data. Each data point in the plot represented a specific query protein plotted by its respective ADS and PPIDR coordinates. The plot utilized color-coded regions to classify proteins according to accepted structural standards. Here, highly disordered proteins were shown in red, moderately disordered proteins were shown in pink or light pink, and ordered proteins were shown in blue or light blue. To illustrate alignment between the two metrics, the shading intensity varied, where dark shades (dark blue or dark pink) marked regions where both the ADS and PPIDR criteria agreed, whereas light shades (light blue or light pink) marked regions where only one of the two criteria was met. B. Classification of query proteins used a combined ΔCH–ΔCDF plot. This two-dimensional graph integrated outputs from charge-hydropathy (CH) analysis (protein net charge vs. hydropathy plot) and cumulative distribution function (CDF) analysis (disorder score frequency vs. disorder scores plot). The X-axis displayed the protein’s disorder frequency deviation from the CDF boundary (ΔCDF), whereas the Y-axis showed its distance from the CH boundary (ΔCH). Proteins were categorized into four structural states based on their quadrant location: Quadrant 1 (blue, bottom right: structured), Quadrant 2 (pink, bottom left: molten globular/hybrid), Quadrant 3 (red, top left: highly disordered), and Quadrant 4 (violet, top right: mixed).

Highly disordered status of unique SRY proteins and unique HMG domains was further supported by Figure XB showing the ΔCH-ΔCDF plot for these datasets. Here, all SRY proteins and HMG domains were located within the quadrants Q3 and Q4. In this plot, quadrant Q3 contained proteins predicted as disordered by both charge-hydropathy (CH) plot and cumulative distribution function (CDF) analysis [64, 65, 66, 67]. Typically, these were highly disordered proteins with minimal residual structure. On the other hand, proteins in quadrant Q4 were predicted as disordered by CH plot but ordered by the CDF analysis. Although these proteins possessed low overall hydropathy and a high net charge, they were classified as ordered by the CDF. Therefore, Q4 proteins, which were typically quite rare (with only a limited number of proteins found in Q4 during typical proteome-wide analyses), exhibited a specific structural paradox. This paradox could be resolved by considering their distinct biophysical traits, such as the presence of localized, highly hydrophobic clusters strategically positioned to drive cooperative folding. Even though the average hydropathy was low, the local hydropathy within the core was high enough to lock the protein (or a part of it) into a stable conformation [68, 69, 70]. Some of these proteins were characterized by high environmental responsiveness and served as illustrative examples of proteins with conditional disorder. Acting as molecular switches, they remained disordered under physiological conditions but transitioned cleanly to a highly ordered state upon binding a partner, shifting pH, or encountering specific ions.

To provide additional information on the unique characteristics of the intrinsic disorder propensity distribution within SRY protein sequences, Figure 30 represented the per-residue disorder profiles generated by the RIDAO platform for three specific proteins. These included the least and most disordered SRY proteins based on their mean disorder prediction (MDP) values, e.g., SRY_ENHLU (8.20%) and SRY_MOUSE (73.21%), respectively, as well as the human SRY protein (25.98%). As shown in Figure 30, different predictors evaluated the disorder profiles of these proteins differently. For example, ADS values derived for SRY_ENHLU by different tools utilized in this study could be arranged as follows: 0.3583 (PONDR® VLXT) < 0.3701 (MDP) < 0.3886 (PONDR® FIT) < 0.4322 (PONDR® VL3) < 0.4430 (IUPred_short) < 0.4849 (IUPred_long) < 0.5459 (PONDR® VSL2). On the other hand, the PPIDR values for this protein are ranged as: 8.22% (MDP) < 24.66% (PONDR® FIT) < 26.48% (PONDR® VLXT) < 35.62% (IUPred_short) < 41.10% (IUPred_long) < 45.66% (PONDR® VL3) < 69.41% (PONDR® VSL2). In the case of SRY_MOUSE, we see the different arrangements. ADS: 0.1638 (PONDR® VLXT) < 0.6231 (MDP) < 0.6421 (PONDR® VL3) < 0.6916 (IUPred_short) < 0.7005 (PONDR® FIT) < 0.7474 (IUPred_long) < 0.7931 (PONDR® VSL2); PPIDR: 12.24% (PONDR® VLXT) < 69.13% (IUPred_short) < 73.21% (MDP) < 73.72% (IUPred_long) < 76.53% (PONDR® FIT) < 81.12% (PONDR® VL3) < 84.69% (PONDR® VSL2) Human SRY protein is characterized by the different disorder metrics as well. ADS: 0.3928 (IUPred_short) < 0.3991 (MDP) < 0.4321 (IUPred_long) < 0.4667 (PONDR® VLXT) < 0.4858 (PONDR® FIT) < 0.5261 (PONDR® VL3) < 0.6173 (PONDR® VSL2); PPIDR: 21.57% (IUPred_short) < 25.98% (MDP) < 32.35% (IUPred_long) < 41.67% (PONDR® VLXT) < 45.10% (PONDR® FIT) < 56.86% (PONDR® VL3) < 92.16% (PONDR® VSL2).

**Figure 30:**
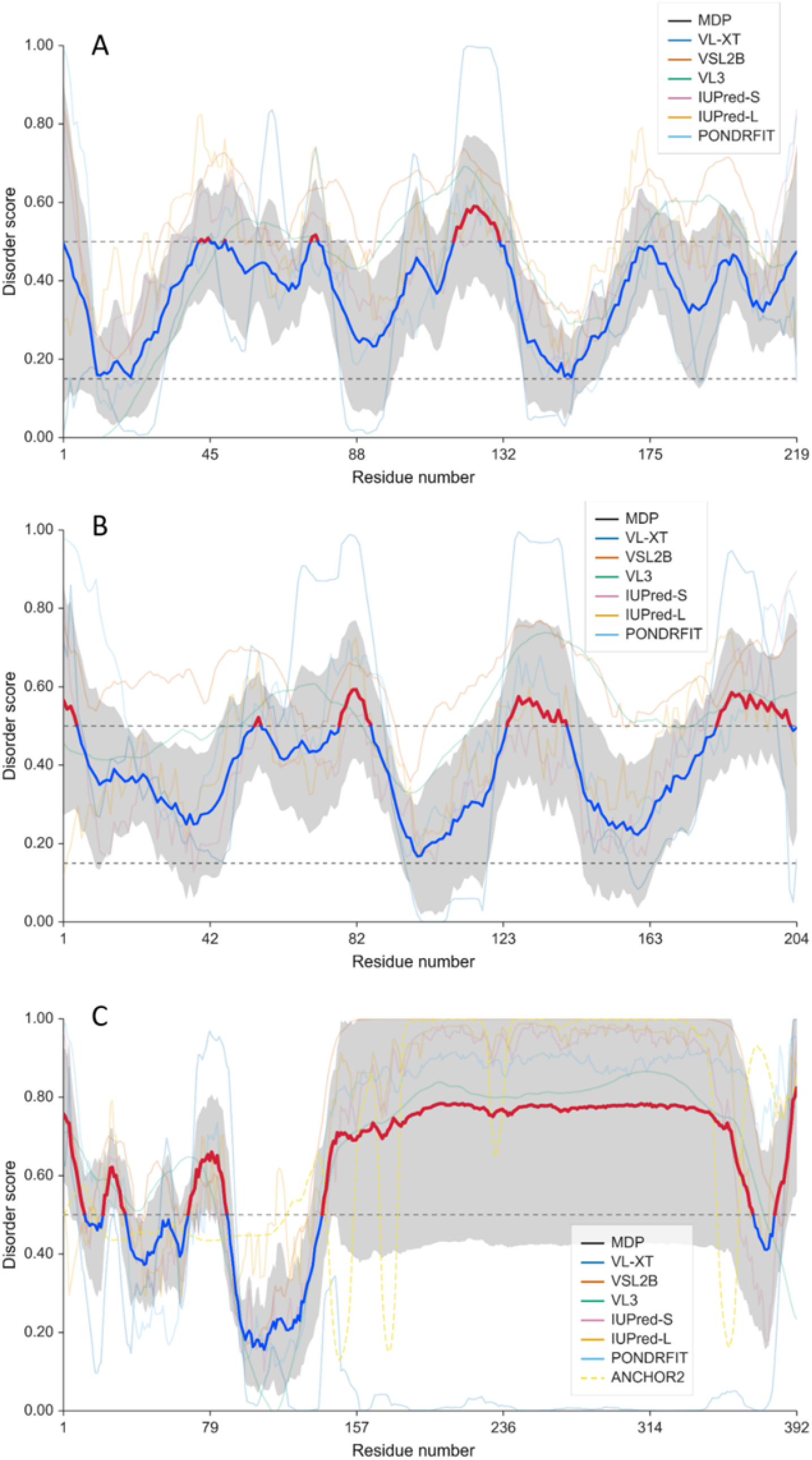
Per-residues intrinsic disorder profiles generated by RIDAO [28]. A. SRY protein from sea otter (*Enhydra lutris*; SRY_ENHLU; UniProt ID: Q6TC27). B. Human SRY protein (*Homo sapiens*; SRY_HUMAN; UniProt ID: Q05066). C. Mouse SRY protein (*Mus musculus*; SRY_MOUSE; UniProt ID: Q05738).

The observed predictor variability could be understood by taking into account that the computational algorithms relied on distinct metrics, such as amino acid composition, hydropathy, charge–charge interactions, or machine learning models, to estimate disorder, which frequently yielded contrasting local propensity scores. However, the mouse SRY protein stood out from this expected variability. As illustrated in Figure YD, although the metrics generated by most of the predictors generally agreed (with ADS values ranging between 0.6231 and 0.7923, and PPIDR values between 69.13% and 84.69%), the profile generated by PONDR® VLXT was noticeably different. This tool predicted the C-terminal two-thirds of the protein (residues 132–385) to be a highly ordered segment, whereas all other tools predicted this long region to be mostly disordered. This discrepancy originated from the fact that this segment contained a highly repetitive sequence with multiple poly-glutamine (polyQ) tracts, with a total glutamine content of 53.36% (119 glutamine residues within this 223-residue-long region spanning residues 144–366). Because PONDR® VLXT was highly sensitive to local amino acid composition and was trained on distinct, non-redundant datasets, it misinterpreted these low-complexity, glutamine-rich repeats as rigid, well-folded structures. In contrast, modern predictors correctly recognized that such repetitive, low-complexity regions lacked the structural constraints required for gaining ordered, well-folded structure, thereby categorizing them as intrinsically disordered.

Finally, Figures 31A, 31C, and 31E provided three-dimensional structural models generated for SRY_ENHLU, SRY_HUMAN, and SRY_MOUSE by AlphaFold PMID: 34265844, whereas Figures 31B, 31D, and 31F showed conformational ensembles generated for these proteins by AFflecto PMID: 40133775. According to AlphaFold structural predictions, all three SRY proteins shared a conserved, small helical HMG domain. In SRY_ENHLU and SRY_HUMAN, the central HMG domain was flanked by extensive intrinsically disordered regions (IDRs), as indicated by low model confidence scores.

**Figure 31:**
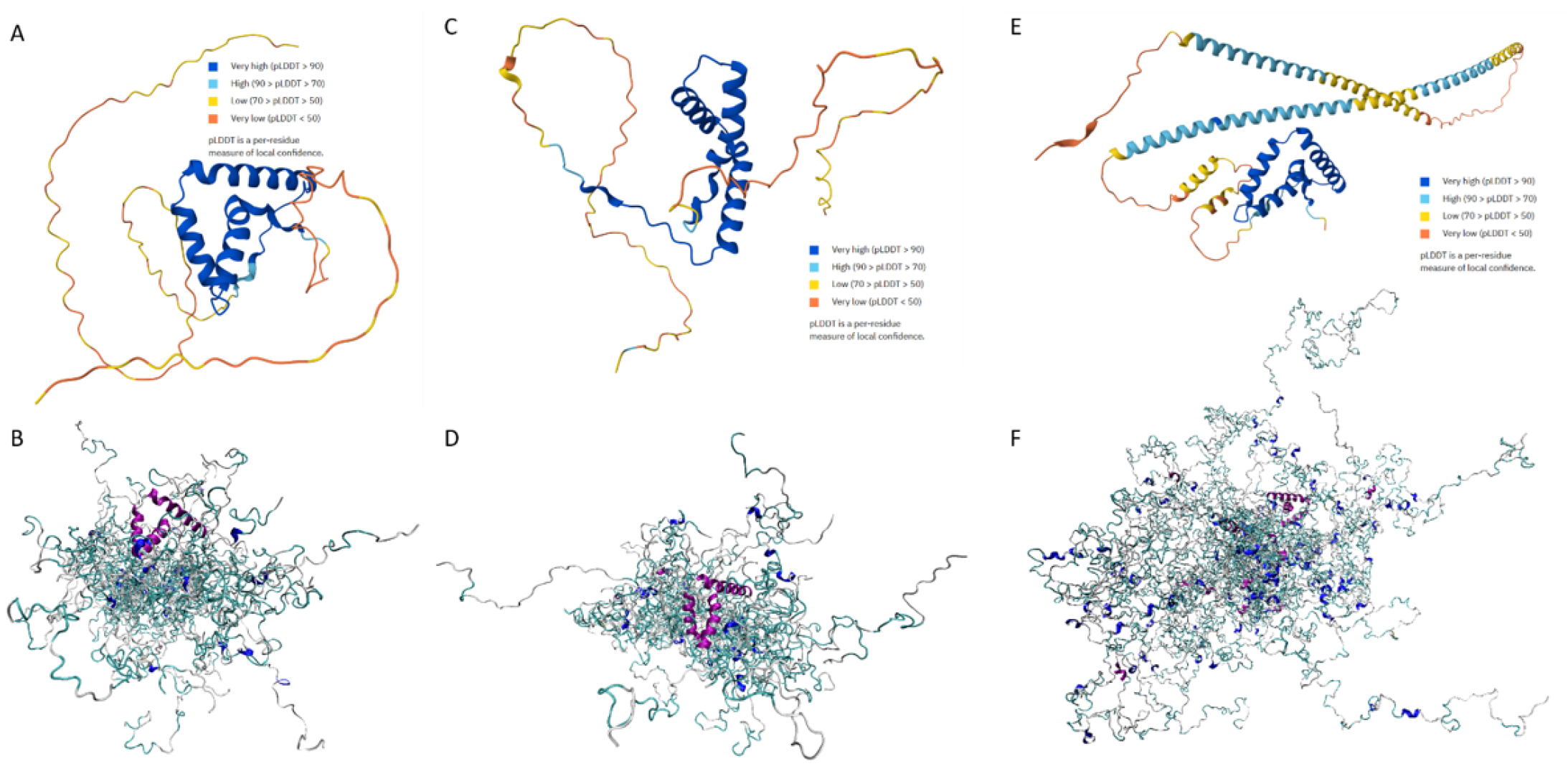
Structural characterization of selected SRY proteins. A, C, and E: 3D models were generated for SRY_ENHLU, SRY_HUMAN, and SRY_MOUSE by AlphaFold [31]. The structures were colored based on the per-residue model confidence (pLDDT), which ranged between 0 and 100. Regions with very high (pLDDT *>* 90), high (90 *>* pLDDT *>* 70), low (70 *>* pLDDT *>* 50), and very low model confidence (pLDDT *<* 50) were shown by blue, cyan, yellow, and orange colors, respectively. Regions with low pLDDT could be unstructured in isolation. B, D, and F: 3D conformational ensembles were generated for SRY_ENHLU, SRY_HUMAN, and SRY_MOUSE by AFflecto [32]. Each ensemble included 25 models. Plots were generated using Visual Molecular Dynamics (VMD) software for molecular visualization [71].

Conversely, for SRY_MOUSE, AlphaFold predicted a starkly distinct topology featuring two long, standalone *α*-helices projecting from the core (see Figure 31E). These terminal expansions exhibited high sequence divergence and were characteristic of the rodent SRY lineage, which frequently incorporated large, repetitive tracts. In computational models, these low-complexity sequences often yielded artificial secondary structures. Therefore, although AlphaFold showed remarkable accuracy in predicting ordered proteins and domains, this highly efficient tool had a tendency to “hallucinate” rigid conformations within intrinsically disordered regions (IDRs) or rapidly evolving, low-complexity repetitive sequences. In these instances, the algorithm often assigned high-confidence local scores (pLDDT) to regions that were biochemically unstructured in vivo. This over-prediction underscored a known limitation: deep-learning models trained on the Protein Data Bank (PDB) were inherently biased toward finding ordered states, sometimes forcing evolutionary noise or highly variable repeat tracts into spurious helical or extended topologies [72, 73, 74, 75, 76].

To resolve structural artifacts in AlphaFold models, we employed web server AFflecto that generated large conformational ensembles, thereby transitioning from a single static structure to an expansive conformational ensemble that accurately captured the flexibility of IDRs. By extracting per-residue confidence (pLDDT) scores from AlphaFold (AF) models, AFflecto effectively distinguished rigid domains from flexible regions [32]. It evaluated structural context to systematically categorize these IDRs as terminal tails, connecting linkers, or internal loops. To accurately chart the massive conformational diversity of these flexible regions, AFflecto leveraged computationally efficient stochastic sampling algorithms, such as Sequential Random Sampling (SRS) and Torsional Random Sampling (TRS) [32]. Furthermore, it addressed a prominent limitation in static deep-learning predictions by incorporating a specialized method to identify conditionally folded secondary structural elements (CF SSEs). These were dynamic segments that AlphaFold often confidently mispredicted as natively folded elements but actually required binding partners or specific conditions to stabilize. Ultimately, AFflecto transformed static structural predictions into realistic, heterogenous protein ensembles [32]. These ensembles provided essential starting configurations for atomistic molecular dynamics (MD) simulations and offered vital templates for back-calculating and interpreting solution-state experimental data, such as NMR and small-angle X-ray scattering (SAXS). Conformational ensembles generated by AFflecto for SRY_ENHLU, SRY_HUMAN, and SRY_MOUSE were presented in Figures 31B, 31D, and 31F, which clearly showed that, with the exception of their structured HMG domains, these proteins were highly disordered. By sampling the vast, highly flexible conformational space of the flanking tails and regions surrounding the central HMG box, the AFflecto server captured the highly dynamic structural heterogeneity characteristic of these transcription factors in solution. Note that no long helices were present in the conformational ensemble generated for SRY_MOUSE, where the extended C-terminal region behaved predominantly as a random coil despite any transient local biases. Therefore, AFflecto successfully avoided the over-packing artifact typical of static machine-learning models, offering a realistic depiction of the expanded volume and structural plasticity of the flanking tails.

### 4.11. Cumulative proximal SRY and SRY-HMG sequences

Intersecting the clusters obtained from eight independent feature-based analyses yielded a set of robust proximal groups for both the SRY protein sequences and the SRY-HMG box subsequences. For the 65 SRY sequences, five clusters remained (Table 34, Figure 32). The largest contained seven sequences from seven mammals namely ARCAU, CALUR, ARCGZ, PAGGO, MIRLE, CYSCR, and MIRAN. For the 38 SRY-HMG subsequences, eight clusters remained (Table 35, Figure 33), consisting of five pairs and three singletons.

**Figure 32:**
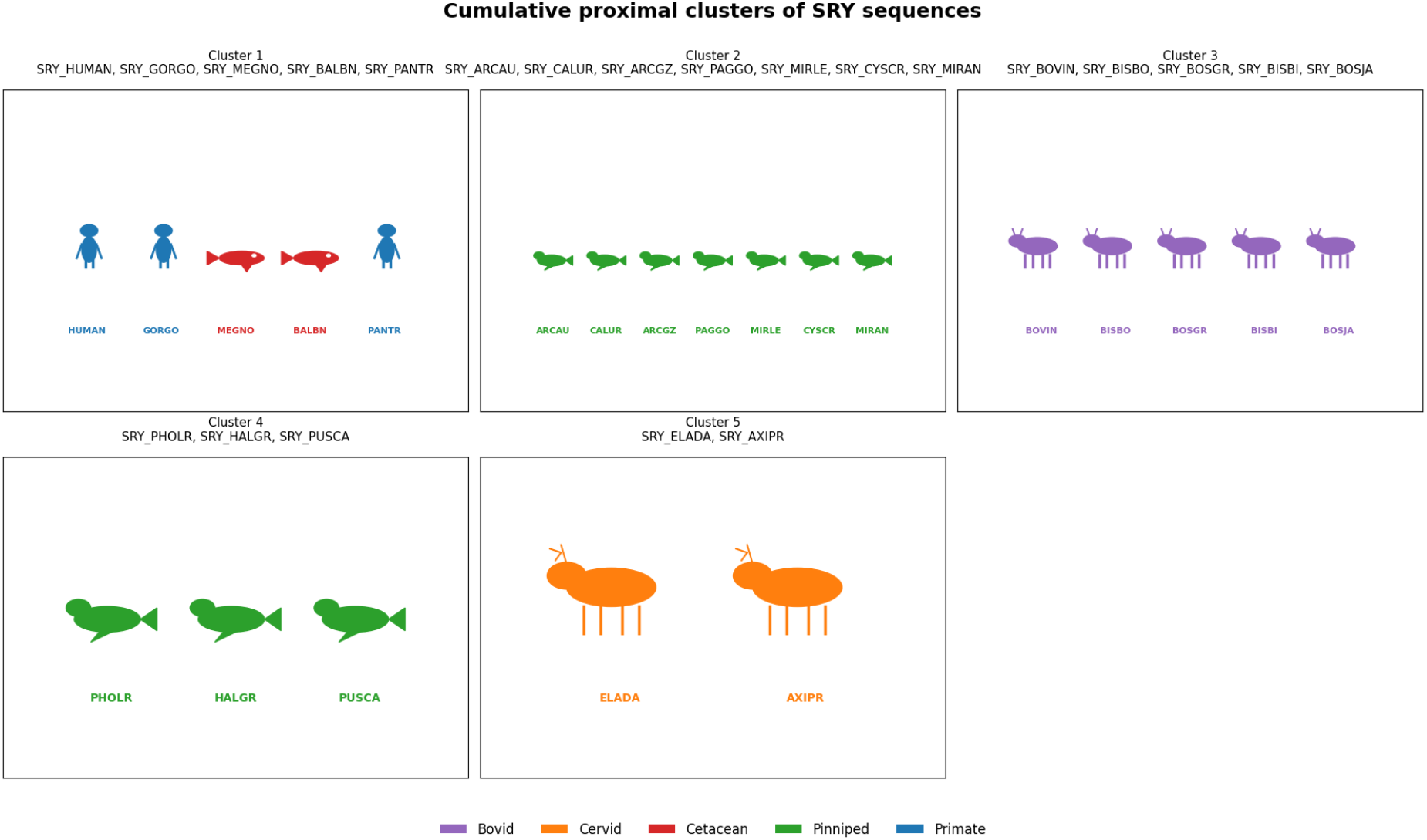
Cumulative proximal clusters of 22 full-length SRY sequences across all feature analyses. Colors denote taxonomic groups: Primates (blue), Cetaceans (red), Pinnipeds (green), Bovids (purple), and Cervids (orange).

**Figure 33:**
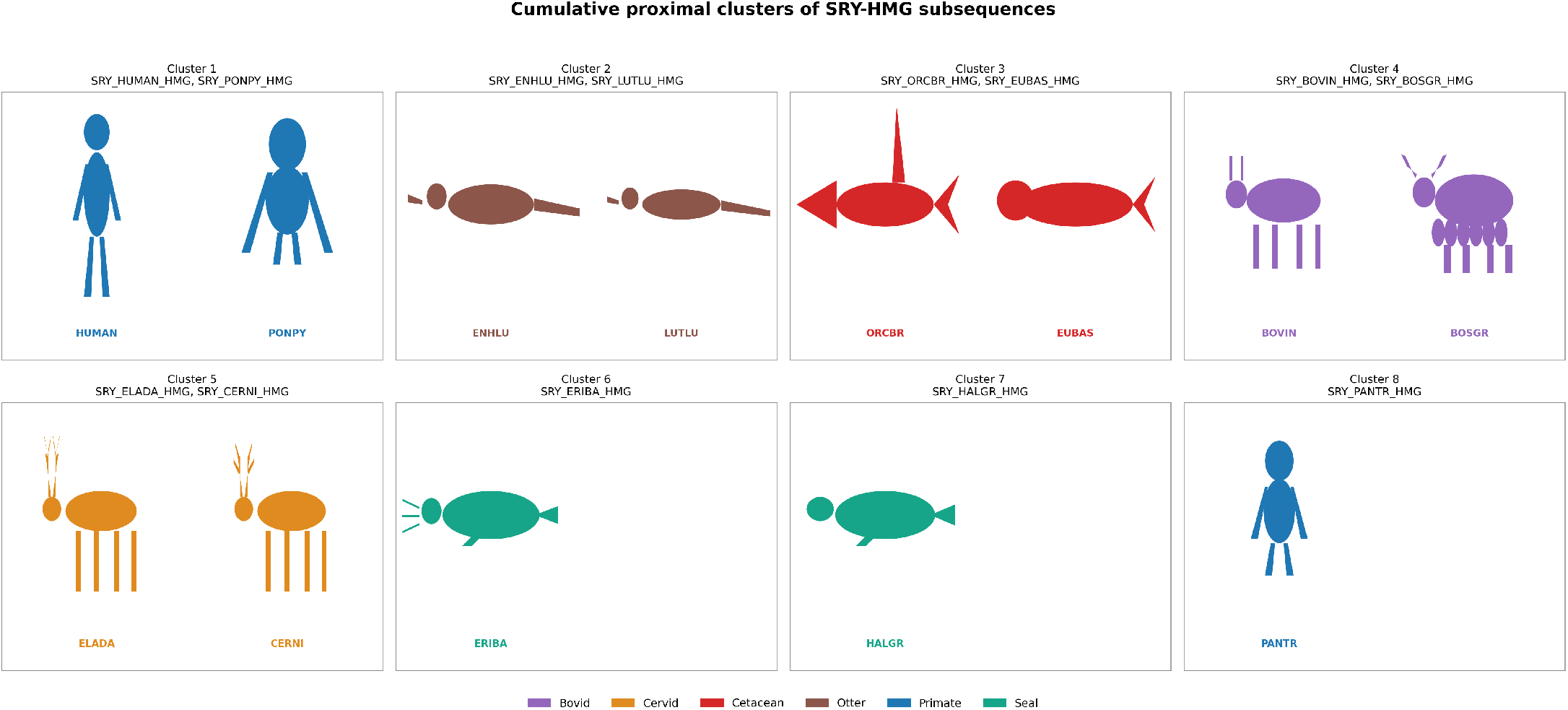
Cumulative proximal clusters of 13 SRY-HMG subsequences across all feature analyses. Colors denote taxonomic groups: Primates (blue), Otters (brown), Cetaceans (red), Bovids (purple), Cervids (orange), and Seals (teal).

**Table 34:** Cumulative list of proximal SRY sequences clusters containing only sequences present in all eight feature-based analyses (amino acid frequency, sequence homology, dipeptide composition, longest polystring, 2-, 3-, and 4-gram hydropathy profiles, and longest polar/nonpolar stretches).

| Cluster | Set of proximal SRY sequences |
| --- | --- |
| Cluster-1 | {SRY_HUMAN, SRY_GORGO, SRY_MEGNO, SRY_BALBN, SRY_PANTR} |
| Cluster-2 | {SRY_ARCAU, SRY_CALUR, SRY_ARCGZ, SRY_PAGGO, SRY_MIRLE, SRY_CYSCR, SRY_MIRAN} |
| Cluster-3 | {SRY_BOVIN, SRY_BISBO, SRY_BOSGR, SRY_BISBI, SRY_BOSJA} |
| Cluster-4 | {SRY_PHOLR, SRY_HALGR, SRY_PUSCA} |
| Cluster-5 | {SRY_ELADA, SRY_AXIPR} |

**Table 35:** Cumulative list of proximal SRY-HMG subsequence clusters containing only sequences present in all eight feature-based analyses (amino acid frequency, sequence homology, dipeptide composition, longest normalized polystring lengths, 2-, 3-, and 4-gram hydropathy profiles, and longest polar/nonpolar stretches).

| Cluster | Set of SRY-HMG subsequences |
| --- | --- |
| 1 | {SRY_HUMAN_HMG, SRY_PONPY_HMG} |
| 2 | {SRY_ENHLU_HMG, SRY_LUTLU_HMG} |
| 3 | {SRY_ORCBR_HMG, SRY_EUBAS_HMG} |
| 4 | {SRY_BOVIN_HMG, SRY_BOSGR_HMG} |
| 5 | {SRY_ELADA_HMG, SRY_CERNI_HMG} |
| 6 | {SRY_ERIBA_HMG} |
| 7 | {SRY_HALGR_HMG} |
| 8 | {SRY_PANTR_HMG} |

Comparison of the cumulative proximal clusters derived from various feature-based analyses revealed a single conserved proximal pairing between the 22 full-length SRY sequences (distributed across 5 clusters) and the 13 SRY-HMG subsequences (distributed across 8 clusters) (Table 34, Table 35). While six organisms (HUMAN, PANTR, BOVIN, BOSGR, HALGR, and ELADA) are present in both cumulative lists, but only BOVIN and BOSGR maintain their proximal relationship across both analyses. In the full-length SRY analysis, SRY_BOVIN and SRY_BOSGR co-cluster in Cluster-3 alongside SRY_BISBO, SRY_BISBI, and SRY_BOSJA. In the SRY-HMG analysis, SRY_BOVIN_HMG and SRY_BOSGR_HMG form Cluster-4 exclusively together. The remaining overlapping organisms fail to maintain their proximal partnerships: HUMAN and PANTR cluster together in full-length Cluster-1, but diverge in the HMG analysis, with SRY_HUMAN_HMG in Cluster-1 and SRY_PANTR_HMG as a singleton in Cluster-8. Likewise, HALGR is proximal to PHOLR and PUSCA in the full-length analysis (Cluster-4), but becomes a singleton in the HMG analysis (Cluster-7); and ELADA shifts its proximal partner from AXIPR (full-length Cluster-5) to CERNI (HMG Cluster-5).

## 5. Discussion and concluding remarks

In this study we employed a couple of quantitative sequence-derived features—amino acid composition, correlation patterns, Shannon entropy, dominant amino acid profiles, dipeptide prevalence, binary hydropathy n-gram spectra, polar/nonpolar composition, and cumulative feature-based clustering—to characterize 65 full-length SRY proteins and 38 SRY-HMG domains across mammals. Collectively, these analyses established a consistent picture: the SRY-HMG domain is a compositionally constrained unit under strong purifying selection, whereas the SRY protein is shaped by rapidly evolving N- and C-terminal regions that introduce substantial lineage-specific variation. This distinction provides a quantitative framework for understanding the functional architecture of SRY and the evolutionary forces acting on it.

The amino acid composition of SRY proteins was enriched in glutamine, lysine, arginine, and serine—residues associated with transcriptional regulation, DNA interaction, and post-translational modification—whereas glycine, isoleucine, cysteine, and tryptophan were frequently depleted. These patterns indicated conservation of residues important for regulatory and DNA-binding functions while limiting amino acids associated with structural rigidity. Consistently, it was found that the SRY-HMG domains exhibited strong enrichment of lysine and arginine, reflecting the established role of the HMG box as the DNA-binding domain responsible for sequence recognition and DNA bending [6, 36, 35]. The near-complete absence of cysteine and depletion of acidic residues within the SRY-HMG domains further highlighted the strong evolutionary conservation of this region. As the SRY-HMG box mediates activation of downstream targets, particularly SOX9, which initiates testis differentiation, maintenance of its characteristic basic amino acid composition is crucial for the male sex-determining function of SRY [6, 4, 77]. While SRY proteins were predominantly serine-rich with lineage-specific enrichment of glutamine, lysine, and arginine, the SRY-HMG domains were mainly enriched in basic residues, with lysine predominating in 14 domains and arginine in 21 domains. In contrast to the variable depletion patterns in SRY protein sequences, the SRY-HMG domains showed a highly conserved depletion profile characterized by the near-complete absence of cysteine and the absence of aspartic acid in SRY_RAT_HMG, SRY_MOUSE_HMG, and SRY_MUSSP_HMG.

Correlation analysis revealed distinct patterns of amino acid co-occurrence. In SRY proteins, strong positive correlations such as Q-H, L-S, R-G, and C-S, together with negative associations including Q-E, R-Q, R-H, and H-S, indicated lineage-specific compositional profiles. The inverse relationship between glutamine and arginine reflected the occurrence of glutamine-rich and arginine-rich SRY variants among mammals. On the other side, the SRY-HMG domains exhibited a more structured correlation network, with positive associations among Q-M, M-W, H-T, and S-T, and strong negative correlations such as A-S, G-M, and Q-R. The persistence of the Q-R negative correlation in both datasets highlighted a conserved compositional pattern within SRY proteins. Overall, the greater diversity of correlations in SRY sequences compared with the SRY-HMG domains was consistent with the strong evolutionary conservation of the SRY-HMG box, which contains residues essential for sequence-specific DNA binding, DNA bending, and activation of SOX9 during mammalian male sex determination [6, 35, 36, 4, 77].

Shannon entropy analysis yielded that SRY proteins varied more widely in sequence complexity (3.4272–4.1901) than their HMG domains (3.8965–4.1063), with rodent SRY proteins showing the lowest entropy (e.g., SRY_MOUSE, 3.4272) and SRY_SMIMA the highest (4.1901), whereas SRY-HMG domains clustered near 4.0. This pattern suggested that the SRY-HMG box was under stronger functional constraint as the essential DNA-binding domain, while the non-HMG regions were more free to diverge and accumulated lineage-specific amino acid changes, consistent with earlier reports of rapid evolution outside the SRY-HMG box [9, 78, 79]. The lower entropy in rodents might have reflected lineage-specific divergence and low-complexity or repetitive sequences in the non-HMG region, whereas the narrower HMG entropy range indicated conservation of the domain’s overall compositional diversity despite some tolerated variation.

Dominant amino acid profiling across the 65 SRY sequences revealed substantial compositional diversity, with 49 distinct classes and most clusters containing few organisms, the largest comprising five bovine sequences sharing the profile ‘V-S-E-R-K-R-R-H-S-S-A-Y’. Recurring dominant residues such as glutamic acid, arginine, lysine, serine, and glutamine in the central regions indicated a conserved compositional framework, whereas the N- and C-terminal regions exhibited greater variation and contributed most to interorganism differences. Murine SRY proteins, particularly SRY_MOUSE, SRY_MUSSP, and SRY_MUSSI, displayed extended glutamine-dominated windows, a lineage-specific feature consistent with known low-complexity regions in rodent SRY [78]. The SRY profile was more complex than that of the HMG-box domain, which showed a compact and conserved profile enriched in arginine and lysine. The repeated enrichment of arginine and lysine in the SRY-HMG box across taxa underscored the importance of positively charged residues for DNA binding and bending, while the overall conservation of the domain’s dominant profile indicated strong purifying selection. Dipeptide prevalence analysis of the 65 SRY proteins revealed a markedly skewed distribution, with the majority of the 400 possible dipeptides falling into rare or absent categories, and only a small subset being highly prevalent. Highly prevalent dipeptides were dominated by polar–polar and polar–charged couples, whereas rare and absent dipeptides were enriched for bulky nonpolar and cysteine-containing couples, consistent with strong purifying selection acting on the functional core of the protein. This compositional bias was less extreme than that observed for the isolated SRY-HMG domain, likely because the sequences included the variable N- and C-terminal regions that diluted the conserved signature of the DNA-binding domain. The persistence of polar and charged dipeptides in the prevalent categories supported the importance of solvent-exposed and DNA-contacting surfaces for SRY-HMG box function [23], while the exclusion of sterically demanding nonpolar couples reflected structural constraints on the domain fold.

Binary hydropathy n-gram analyses revealed a clear contrast between SRY proteins and their SRY-HMG subsequences. In SRY sequences, polar patterns dominated, with 11, 111, and 1111 being the most frequent words and murine sequences (SRY_MOUSE, SRY_MUSSP, and SRY_MUSSI) showing the strongest polar bias, whereas species such as SRY_ORCBR, SRY_GLOMA, and SRY_GRAGR had the lowest polar frequencies. In contrast, the SRY-HMG subsequences exhibited a more even hydropathy balance: although 11 remained the most common 2-gram, its frequency was reduced, 00 increased, and 011 and 110 were the most abundant 3-grams, while 111 was less dominant and the 4-gram frequencies were more evenly spread. Many SRY-HMG subsequences shared identical 2-gram and 3-gram profiles, with only a few outliers such as SRY_SMIMA_HMG (more polar) and SRY_RAT_HMG/SRY_MUSSP_HMG (less polar). This pattern suggested that the SRY-HMG box maintained a mixed polar–nonpolar architecture compatible with its DNA-binding and bending function [23], whereas the SRY sequences were enriched in polar runs partly, because of their variable N- and C-terminal regions. The greater uniformity of the SRY-HMG hydropathy profiles supported stronger purifying selection on this domain, while the broader variation in sequences was consistent with rapid evolution outside the HMG box [9, 78, 79].

Polar/nonpolar composition analysis revealed that although polar residues predominated in both SRY proteins and their SRY-HMG domains, the two sets differed markedly in both mean polar content and spread. SRY sequences had a higher mean polar content (61.96%) and a wide range (57.21–76.28%), with the highest values in rodents such as SRY_MOUSE (76.28%) and the lowest in cetaceans such as SRY_ORCBR and SRY_DELLE (57.21%). In contrast, the SRY-HMG domains were tightly constrained, with a mean of 58.44%, a narrow range (56.52–62.32%), and 25 of 38 subsequences sharing exactly 57.97% polar residues. This uniformity suggested that the SRY-HMG box was under strong purifying selection to maintain a specific fold for DNA binding [52, 62], whereas the lineage-specific variation seen in SRY, such as the high polar content in rodents and low polar content in cetaceans, was largely absent from the SRY-HMG domain [63]. The compositional freedom of the protein was therefore not mirrored in its DNA-binding domain, which remained close to a fixed polar/nonpolar balance across organisms.

Furthermore, across 65 SRY proteins and 38 HMG domains, intrinsic disorder was the dominant predicted state, with all full-length SRY proteins highly disordered (PPIDR 51.27–94.61%; ADS 0.5119–0.7931) and 31 of 38 HMG domains also highly disordered, while Q3/Q4 localization supported a conditional-disorder model rather than uniform disorder. Predictor variability and the AlphaFold artifact in SRY_MOUSE—where polyQ-rich low-complexity repeats were mispredicted as ordered or helical but AFflecto ensembles resolved them as random-coil-like—showed that consensus disorder analysis combined with ensemble modeling was necessary to avoid overestimating structure in rapidly evolving IDRs.

Intersecting the clusters obtained from eight independent feature-based analyses yielded robust proximal groups for both the full-length SRY sequences and the SRY-HMG subsequences, with five clusters retained for the 22 full-length SRY proteins and eight clusters for the 13 SRY-HMG domains. The cumulative proximity of BOVIN and BOSGR at both the SRY and SRY-HMG levels was consistent with their close phylogenetic relationship, whereas the other shared organisms displayed domain-specific patterns: HUMAN grouped with GORGO, MEGNO, BALBN, and PANTR at the SRY level, but with PONPY_HMG at the HMG level; ELADA grouped with AXIPR at the full-length level, but with CERNI_HMG at the SRY-HMG level; and HALGR grouped with PHOLR and PUSCA at the full-length level but became a singleton at the HMG level. These differences likely arose because the full-length protein includes disordered terminal regions that can obscure domain-level similarities and introduce lineage-specific clustering signals [9, 79]. The fact that some SRY-HMG domain clusters were more consistent with taxonomic expectations, such as the bovid pair, supported the view that this domain evolves more slowly and under stronger purifying selection than the flanking regions [52, 62]. The presence of several singletons and unique clusters in both datasets may have reflected genuine species-specific divergence or, alternatively, incomplete taxon sampling and the inherent limitations of feature-based clustering.

The quantitative signatures established here carry direct implications for understanding SRY biology. The SRY-HMG domain is the only region of SRY that is conserved across mammals and is both necessary and sufficient for sequence-specific DNA binding and bending [62, 52]. The tight clustering of SRY-HMG entropy values near 4.0, the narrow polar content range (5.80 percentage points), the repeated enrichment of arginine and lysine in dominant windows, and the prevalence of polar–polar and polar–charged dipeptides collectively indicate that the SRY-HMG domain operates within a narrow compositional window. This window is presumably dictated by the need to maintain a functional DNA-binding surface, a proper hydrophobic core, and the structural flexibility required for DNA bending [62]. The observation that 25 of 38 SRY-HMG subsequences share exactly 57.97% polar residues is particularly striking and suggests that the domain has been optimized to a specific polar/nonpolar balance that cannot be substantially altered without compromising function. On the other side, the SRY protein exhibited a much wider compositional range, with rodents showing the highest polar content (SRY_MOUSE, 76.28%) and cetaceans the lowest (SRY_ORCBR and SRY_DELLE, 57.21%). This variation is not random; it reflects the known rapid evolution of the non-HMG regions, which contain transactivation domains, repetitive elements, and lineage-specific regulatory motifs [9, 78, 79]. The glutamine-rich windows in murine SRY, for example, are consistent with the well-documented glutamine-rich transactivation domain in mouse SRY, which is not conserved in other mammals [78]. The high polar content in rodents may therefore reflect the expansion of polar-rich low-complexity regions, whereas the low polar content in cetaceans may reflect lineage-specific loss or reduction of such regions.

Several additional observations warrant discussion. First, the outlier status of SRY_SMIMA_HMG, which combined the lowest 011 frequency with the highest 111 frequency and the lowest 110 frequency in the 3-gram analysis, and the mirror-image pattern in SRY_RAT_HMG and SRY_MUSSP_HMG, suggest that even within the conserved HMG domain, lineage-specific compositional drift can occur. Whether such drift is functionally neutral or has subtle effects on DNA binding or bending remains to be tested experimentally. Second, the near-symmetry of the 01 and 10 transition frequencies in the 2-gram analysis, and the numerical identity of the 0111 and 1110 columns in the 4-gram analysis, reflect the binary encoding of linear sequences and serve as an internal consistency check on the methodology; deviations from this symmetry, such as those observed in specific sequences, may indicate directional compositional biases worth investigating. Third, the discrepancy between full-length SRY and HMG-level clustering for HUMAN, ORCBR, and ELADA suggests that the disordered terminal regions of SRY can obscure domain-level phylogenetic signals, and that domain-specific analyses are necessary for accurate functional inference. Fourth, the presence of several singletons and small clusters in both datasets may reflect genuine species-specific divergence, incomplete taxon sampling, or the inherent limitations of feature-based clustering; future work with denser taxon sampling and complementary methods such as phylogenetic reconciliation or structural modelling could help distinguish these possibilities. Fifth, the persistent Q-R negative correlation in both full-length and HMG datasets highlights a conserved compositional constraint that may be linked to the functional dichotomy between glutamine-rich and arginine-rich regions in different lineages.

Taken together, these observations suggest that the conserved polar/nonpolar balance of the SRY-HMG domain— particularly the 57.97% polar fraction shared by most subsequences—likely reflects a functional optimum for DNA binding and bending, with deviations from this balance being potentially deleterious. The lineage-specific compositional extremes seen in full-length SRY, such as the high polar content in rodents and low polar content in cetaceans, appear to reflect adaptive or relaxed selection on the non-HMG regions rather than on the DNA-binding domain itself. The HMG domain may tolerate a limited amount of compositional drift, as seen in SRY_SMIMA_HMG and SRY_RAT_HMG/SRY_MUSSP_HMG, without loss of core function, consistent with the reported relaxation of purifying selection on SRY in some lineages [79]. The cumulative clustering approach, by integrating eight independent feature-based analyses, provides a more robust grouping of sequences than any single feature alone, and the clusters identified here may serve as a foundation for future functional and evolutionary studies of SRY across mammals.

Projection of the unique full-length SRY proteins and SRY-HMG box subsequences into a multidimensional feature space resolved nearly every sequence as compositionally distinct, revealing individuality not captured by alignment-based homology. Multivariate proximity across this space proved to be a stringent criterion of compositional similarity, satisfied by only a single mammalian pair. These findings demonstrated that SRY protein and its HMG box sequence each carried a distinct compositional grammar, quantifiable through alignment-based as well as alignment–independent signatures, with compositional individuality being the rule rather than the exception. This framework offered a new lens for understanding SRY’s functional architecture and the relationship between sequence composition and reproductive function. SRY evolution was thus shaped by domain-specific selective pressures: the HMG box maintained a constrained compositional profile, while flanking regions accumulated lineage-specific diversity. Denser sampling and experimental validation would be essential to determine whether this compositional drift carried functional consequences.

## Acknowledgments

We thank Debaleena Nawn for her invaluable help in developing various quantitative methods that we employ in Mathematical Genomics. APA was supported by the Ministry of Science and Higher Education of the Russian Federation, grant No 075 *−* 15 *−* 2025 *−* 525 (Agreement dated 30 May 2025).

## Author contributions statement

SSH formulated and designed the study protocol. SSH, NM, MS, GWD II, APA, and VNU conducted the theoretical experiments and obtained the results. SSH, NM, MS, GWD II, APA, and VNU performed the quantitative analyses. AG, VNU, and SSH supervised the project. All authors reviewed, checked, and approved the final version of the manuscript.

## References

[1] Y. Katsura, H. X. Kondo, J. Ryan, V. Harley, Y. Satta, The evolutionary process of mammalian sex determination genes focusing on marsupial sry s, BMC Evolutionary Biology 18 (1) (2018) 3.

[2] L. Zhao, P. Koopman, Sry protein function in sex determination: thinking outside the box, Chromosome research 20 (1) (2012) 153–162.

[3] Y.-S. Chen, J. D. Racca, M. A. Weiss, Tenuous transcriptional threshold of human sex determination. i. sry and swyer syndrome at the edge of ambiguity, Frontiers in Endocrinology 13 (2022) 945030.

[4] R. Sekido, R. Lovell-Badge, Sex determination involves synergistic action of sry and sf1 on a specific sox9 enhancer, Nature 453 (7197) (2008) 930–934.

[5] N. Gonen, C. R. Futtner, S. Wood, S. A. Garcia-Moreno, I. M. Salamone, S. C. Samson, R. Sekido, F. Poulat, D. M. Maatouk, R. Lovell-Badge, Sex reversal following deletion of a single distal enhancer of sox9, Science 360 (6396) (2018) 1469–1473.

[6] J. D. Racca, Y.-S. Chen, A. R. Brabender, U. Battistin, M. A. Weiss, M. M. Georgiadis, Role of nucleobase-specific interactions in the binding and bending of dna by human male sex determination factor sry, Journal of Biological Chemistry 300 (9) (2024).

[7] V. R. Harley, S. Layfield, C. L. Mitchell, J. K. Forwood, A. P. John, L. J. Briggs, S. G. McDowall, D. A. Jans, Defective importin *β* recognition and nuclear import of the sex-determining factor sry are associated with xy sex-reversing mutations, Proceedings of the National Academy of Sciences 100 (12) (2003) 7045–7050.

[8] S. Eggers, S. Sadedin, J. A. Van Den Bergen, G. Robevska, T. Ohnesorg, J. Hewitt, L. Lambeth, A. Bouty, I. M. Knarston, T. Y. Tan, et al., Disorders of sex development: insights from targeted gene sequencing of a large international patient cohort, Genome biology 17 (1) (2016) 243.

[9] L. S. Whitfield, R. Lovell-Badge, P. N. Goodfellow, Rapid sequence evolution of the mammalian sex-determining gene sry, Nature 364 (6439) (1993) 713–715.

[10] Y.-S. Chen, J. D. Racca, P. W. Sequeira, N. B. Phillips, M. A. Weiss, Microsatellite-encoded domain in rodent sry functions as a genetic capacitor to enable the rapid evolution of biological novelty, Proceedings of the National Academy of Sciences 110 (33) (2013) E3061–E3070.

[11] L. Zhao, E. T. Ng, T.-L. Davidson, E. Longmuss, J. Urschitz, M. Elston, S. Moisyadi, J. Bowles, P. Koopman, Structure–function analysis of mouse sry reveals dual essential roles of the c-terminal polyglutamine tract in sex determination, Proceedings of the National Academy of Sciences 111 (32) (2014) 11768–11773.

[12] M. H. Bailey, W. U. Meyerson, L. J. Dursi, L.-B. Wang, G. Dong, W.-W. Liang, A. Weerasinghe, S. Li, Y. Li, S. Kelso, et al., Retrospective evaluation of whole exome and genome mutation calls in 746 cancer samples, Nature communications 11 (1) (2020) 4748.

[13] A. Zielezinski, S. Vinga, J. Almeida, W. M. Karlowski, Alignment-free sequence comparison: benefits, applications, and tools, Genome biology 18 (1) (2017) 186.

[14] S. F. Banani, H. O. Lee, A. A. Hyman, M. K. Rosen, Biomolecular condensates: organizers of cellular biochemistry, Nature reviews Molecular cell biology 18 (5) (2017) 285–298.

[15] M. Hardenberg, A. Horvath, V. Ambrus, M. Fuxreiter, M. Vendruscolo, Widespread occurrence of the droplet state of proteins in the human proteome, Proceedings of the National Academy of Sciences 117 (52) (2020) 33254–33262.

[16] A. Hatos, S. C. Tosatto, M. Vendruscolo, M. Fuxreiter, Fuzdrop on alphafold: visualizing the sequence-dependent propensity of liquid–liquid phase separation and aggregation of proteins, Nucleic acids research 50 (W1) (2022) W337–W344.

[17] M. H. Smith, The amino acid composition of proteins, Journal of Theoretical Biology 13 (1966) 261–282.

[18] S. S. Hassan, K. Hungyo, A. Hromić-Jahjefendić, V. N. Uversky, Mathematical sequence analyses of cystic fibrosis transmembrane conductance regulator (cftr): Cross-species skeletal frameworks in cystic fibrosis, Current Genomics (2026).

[19] S. S. Hassan, M. Sil, S. Chakraborty, A. Goswami, P. Basu, D. Nawn, V. N. Uversky, Possible functional proximity of various organisms based on the bioinformatics analysis of their taste receptors, International Journal of Biological Macromolecules 222 (2022) 2105–2121.

[20] S. S. Hassan, D. Nawn, P. Goswami, N. Mukherjee, M. Sil, S. Roy, A. Goswami, S. Das, V. N. Uversky, Quantitative deciphering of mammalian histamine receptors through mathematical genomics, Biochemical and Biophysical Research Communications (2026) 153717.

[21] J. G. Kimmel, T. J. Buchinger, S. Tamrakar, B. Huerta, D. L. Larson, E. A. Baker, T. G. Zorn, K. T. Scribner, W. Li, Spatiotemporal patterns in profiles of amino acids indicates they are unlikely singular olfactory cues underlying natal homing in migratory fishes, Freshwater Biology 70 (2) (2025) e14382.

[22] B. J. Strait, T. G. Dewey, The shannon information entropy of protein sequences, Biophysical journal 71 (1) (1996) 148–155.

[23] J. Lehmann, A. Libchaber, B. D. Greenbaum, Fundamental amino acid mass distributions and entropy costs in proteomes, Journal of Theoretical Biology 410 (2016) 119–124.

[24] S. S. Hassan, S. Ghosh, D. Attrish, P. P. Choudhury, A. A. Aljabali, B. D. Uhal, K. Lundstrom, N. Rezaei, V. N. Uversky, M. Seyran, et al., Possible transmission flow of sars-cov-2 based on ace2 features, Molecules 25 (24) (2020) 5906.

[25] F. Sievers, D. G. Higgins, The clustal omega multiple alignment package, in: Multiple sequence alignment: Methods and protocols, Springer, 2020, pp. 3–16.

[26] F. Madeira, N. Madhusoodanan, J. Lee, A. Eusebi, A. Niewielska, A. R. Tivey, R. Lopez, S. Butcher, The embl-ebi job dispatcher sequence analysis tools framework in 2024, Nucleic acids research 52 (W1) (2024) W521–W525.

[27] M. Vendruscolo, M. Fuxreiter, Sequence determinants of the aggregation of proteins within condensates generated by liquid-liquid phase separation, Journal of Molecular Biology 434 (1) (2022) 167201.

[28] G. W. Dayhoff, V. N. Uversky, Rapid prediction and analysis of protein intrinsic disorder, Protein Science 31 (12) (2022) e4496.

[29] K. Rajagopalan, S. M. Mooney, N. Parekh, R. H. Getzenberg, P. Kulkarni, A majority of the cancer/testis antigens are intrinsically disordered proteins, Journal of cellular biochemistry 112 (11) (2011) 3256–3267.

[30] N. Milkovic, T. Mittag, B. Kragelund, K. Skriver, Intrinsically disordered proteins: Methods and protocols, Methods Mol Biology Clifton NJ 2141 (2020) 685–702.

[31] J. Jumper, R. Evans, A. Pritzel, T. Green, M. Figurnov, O. Ronneberger, K. Tunyasuvunakool, R. Bates, A. Žídek, A. Potapenko, et al., Highly accurate protein structure prediction with alphafold, nature 596 (7873) (2021) 583–589.

[32] M. Pajkos, I. Clerc, C. Zanon, P. Bernadó, J. Cortés, Afflecto: A web server to generate conformational ensembles of flexible proteins from alphafold models, Journal of Molecular Biology 437 (15) (2025) 169003.

[33] R. Sekido, I. Bar, V. Narváez, G. Penny, R. Lovell-Badge, Sox9 is up-regulated by the transient expression of sry specifically in sertoli cell precursors, Developmental biology 274 (2) (2004) 271–279.

[34] R. Sekido, Sry: A transcriptional activator of mammalian testis determination, The international journal of biochemistry & cell biology 42 (3) (2010) 417–420.

[35] E. C. Murphy, V. B. Zhurkin, J. M. Louis, G. Cornilescu, G. M. Clore, Structural basis for sry-dependent 46-x, y sex reversal: modulation of dna bending by a naturally occurring point mutation, Journal of molecular biology 312 (3) (2001) 481–499.

[36] S. Ferrari, V. R. Harley, A. Pontiggia, P. N. Goodfellow, R. Lovell-Badge, M. E. Bianchi, Sry, like hmg1, recognizes sharp angles in dna., The EMBO journal 11 (12) (1992) 4497–4506.

[37] S. V. Kathuria, Y. H. Chan, R. P. Nobrega, A. Özen, C. R. Matthews, Clusters of isoleucine, leucine, and valine side chains define cores of stability in high-energy states of globular proteins: Sequence determinants of structure and stability, Protein Science 25 (3) (2016) 662–675.

[38] U. Narendra, L. Zhu, B. Li, J. Wilken, M. A. Weiss, Sex-specific gene regulation: the doublesex dm motif is a bipartite dna-binding domain, Journal of Biological Chemistry 277 (45) (2002) 43463–43473.

[39] H. R. Kilgore, R. T. Raines, n *π*\* interactions modulate the properties of cysteine residues and disulfide bonds in proteins, Journal of the American Chemical Society 140 (50) (2018) 17606.

[40] F. Hecht, I. S. Harris, Lysosomal cystine: an unexpected alarm bell for cysteine scarcity, Trends in cell biology 33 (12) (2023) 1007–1009.

[41] J. D. Racca, Y.-S. Chen, J. D. Maloy, N. Wickramasinghe, N. B. Phillips, M. A. Weiss, Structure-function relationships in human testis-determining factor sry: An aromatic buttress underlies the specific dna-bending surface of a high mobility group (hmg) box*, Journal of Biological Chemistry 289 (47) (2014) 32410–32429.

[42] J. D. Racca, Y.-S. Chen, Y. Yang, N. B. Phillips, M. A. Weiss, Human sex determination at the edge of ambiguity: inherited xy sex reversal due to enhanced ubiquitination and proteasomal degradation of a master transcription factor, Journal of Biological Chemistry 291 (42) (2016) 22173–22195.

[43] D. G. Ross, J. Bowles, P. Koopman, S. Lehnert, New insights into sry regulation through identification of 5’conserved sequences, BMC molecular biology 9 (1) (2008) 85.

[44] M. E. Turner, D. Ely, J. Prokop, A. Milsted, Sry, more than testis determination?, American Journal of Physiology-Regulatory, Integrative and Comparative Physiology 301 (3) (2011) R561–R571.

[45] M. A. Weiss, Molecular mechanisms of male sex determination: the enigma of sry, in: DNA Conformation and Transcription, Springer, 2005, pp. 159–173.

[46] R. Rimini, A. Pontiggia, F. Spada, S. Ferrari, V. Harley, P. Goodfellow, M. Bianchi, Interaction of normal and mutant sry proteins with dna, Philosophical Transactions of the Royal Society of London. Series B: Biological Sciences 350 (1333) (1995) 215–220.

[47] V. H. Salinas, R. Ranganathan, Coevolution-based inference of amino acid interactions underlying protein function, elife 7 (2018) e34300.

[48] H. Okuyama, A. Tokunaga, S. Hayashi, K. Nakasako, D. Tsukamoto, T. Matsuo, K. Tamura, M. Ito, The tolerated functional decline model: Relaxed constraint in the molecular evolution of mammalian master sex-determining gene sry, Gene (2025) 149844.

[49] K. F. Meyer, L. G. Freitas Filho, K. I. Silva, P. A. Trauzcinsky, C. Reuter, M. B. M. Souza, The xy female and swyer syndrome, Urology Case Reports 26 (2019) 100939.

[50] N. Behtash, M. Karimi Zarchi, Dysgerminoma in three patients with swyer syndrome, World journal of surgical oncology 5 (1) (2007) 71.

[51] A. Hassan-Abdallah, G. Zhao, Z.-w. Chen, F. S. Mathews, M. Schuman Jorns, Arginine 49 is a bifunctional residue important in catalysis and biosynthesis of monomeric sarcosine oxidase: A context-sensitive model for the electrostatic impact of arginine to lysine mutations, Biochemistry 47 (9) (2008) 2913–2922.

[52] W. Yang, Y. Xu, J. Wu, W. Zeng, Y. Shi, Solution structure and dna binding property of the fifth hmg box domain in comparison with the first hmg box domain in human upstream binding factor, Biochemistry 42 (7) (2003) 1930–1938.

[53] A. D. Baxevanis, D. Landsman, The hmg-1 box protein family: classification and functional relationships, Nucleic Acids Research 23 (9) (1995) 1604–1613.

[54] J. Teter, K. Boczkowski, Occurrence of tumors in dysgenetic gonads, Cancer 20 (8) (1967) 1301–1310.

[55] J. Assumpção, C. Benedetti, A. Maciel-Guerra, G. Guerra, M. Baptista, M. Scolfaro, M. de Mello, Novel mutations affecting sry dna-binding activity: the hmg box n65h associated with 46, xy pure gonadal dysgenesis and the familial non-hmg box r30i associated with variable phenotypes, Journal of Molecular Medicine 80 (12) (2002) 782–790.

[56] S. Andonova, R. Robeva, M. Sirakov, K. Mainhard, A. Tomova, S. Ledig, P. Kumanov, A. Savov, A novel sry gene mutation p. f109l in a 46, xy female with complete gonadal dysgenesis, Sexual Development 9 (6) (2016) 333–337.

[57] P. Canto, D. Soderlund, E. Reyes, J. Mendez, Mutations in the desert hedgehog (dhh) gene in patients with 46, xy complete pure gonadal dysgenesis, The Journal of Clinical Endocrinology & Metabolism 89 (9) (2004) 4480–4483.

[58] C. Capito, M.-D. Leclair, A. Arnaud, A. David, S. Baron, N. Corradini, Y. Héloury, 46, xy pure gonadal dysgenesis: clinical presentations and management of the tumor risk, Journal of pediatric urology 7 (1) (2011) 72–75.

[59] A. Coutin, A. Hamy, M. Fondevilla, B. Savigny, J. Paineau, J. Visset, Pure 46xy gonadal dysgenesis, Journal de Gynecologie, Obstetrique et Biologie de la Reproduction 25 (8) (1996) 792–796.

[60] M. Ferguson-Smith, D. Alexander, P. Bowen, R. Goodman, B. Kaufmann, H. Jones, Jr, R. Heller, Clinical and cytogenetical studies in female gonadal dysgenesis and their bearing on the cause of turner’s syndrome, Cytogenetic and Genome Research 3 (6) (2004) 355–383.

[61] Y. Luan, Z. Tang, Y. He, Z. Xie, Intra-domain residue coevolution in transcription factors contributes to dna binding specificity, Microbiology Spectrum 11 (2) (2023) e03651–22.

[62] C. Read, P. D. Cary, C. Crane-Robinson, P. C. Driscoll, M. Carrillo, D. G. Norman, The structure of the hmg box and its interaction with dna, in: Nucleic acids and molecular biology, Springer, 1995, pp. 222–250.

[63] K. R. Richard, S. W. McCarrey, J. M. Wright, Dna sequence from the sry gene of the sperm whale (physeter macrocephalus) for use in molecular sexing, Canadian Journal of Zoology 72 (5) (1994) 873–877.

[64] V. N. Uversky, J. R. Gillespie, A. L. Fink, Why are “natively unfolded” proteins unstructured under physiologic conditions?, Proteins: structure, function, and bioinformatics 41 (3) (2000) 415–427.

[65] A. K. Dunker, P. Romero, Z. Obradovic, E. C. Garner, C. J. Brown, Intrinsic protein disorder in complete genomes, Genome informatics 11 (2000) 161–171.

[66] C. J. Oldfield, Y. Cheng, M. S. Cortese, C. J. Brown, V. N. Uversky, A. K. Dunker, Comparing and combining predictors of mostly disordered proteins, Biochemistry 44 (6) (2005) 1989–2000.

[67] B. Xue, C. J. Oldfield, A. K. Dunker, V. N. Uversky, Cdf it all: consensus prediction of intrinsically disordered proteins based on various cumulative distribution functions, FEBS letters 583 (9) (2009) 1469–1474.

[68] A. Mohan, W. J. Sullivan Jr, P. Radivojac, A. K. Dunker, V. N. Uversky, Intrinsic disorder in pathogenic and non-pathogenic microbes: discovering and analyzing the unfoldomes of early-branching eukaryotes, Molecular BioSystems 4 (4) (2008) 328–340.

[69] F. Huang, C. Oldfield, J. Meng, W.-L. Hsu, B. Xue, V. N. Uversky, P. Romero, A. K. Dunker, Subclassifying disordered proteins by the ch-cdf plot method, in: Biocomputing 2012, World Scientific, 2012, pp. 128–139.

[70] F. Huang, C. J. Oldfield, B. Xue, W.-L. Hsu, J. Meng, X. Liu, L. Shen, P. Romero, V. N. Uversky, A. K. Dunker, Improving protein order-disorder classification using charge-hydropathy plots, BMC bioinformatics 15 (Suppl 17) (2014) S4.

[71] W. Humphrey, A. Dalke, K. Schulten, Vmd: visual molecular dynamics, Journal of molecular graphics 14 (1) (1996) 33–38.

[72] K. M. Ruff, R. V. Pappu, Alphafold and implications for intrinsically disordered proteins, Journal of molecular biology 433 (20) (2021) 167208.

[73] C. J. Wilson, W.-Y. Choy, M. Karttunen, Alphafold2: a role for disordered protein/region prediction?, International journal of molecular sciences 23 (9) (2022) 4591.

[74] A. O. Stevens, Y. He, Benchmarking the accuracy of alphafold 2 in loop structure prediction, Biomolecules 12 (7) (2022) 985.

[75] M. Mehdiabadi, S. C. Tosatto, D. Piovesan, Modeling intrinsically disordered regions from alphafold2 to alphafold3, Protein Science 35 (1) (2026) e70426.

[76] S. Gopalan, S. Narayanan, Hallucinations in alphafold3 for intrinsically disordered proteins with disorder in biological process residues, arXiv preprint arXiv:2510.15939 (2025).

[77] D. Wilhelm, S. Palmer, P. Koopman, Sex determination and gonadal development in mammals, Physiological reviews 87 (1) (2007) 1–28.

[78] P. K. Tucker, B. L. Lundrigan, The nature of gene evolution on the mammalian y chromosome: lessons from sry, Philosophical Transactions of the Royal Society of London. Series B: Biological Sciences 350 (1333) (1995) 221–227.

[79] P. Pamilo, P. Gertsch, P. Thorén, P. Seppä, Molecular population genetics of social insects, Annual Review of Ecology and Systematics 28 (1) (1997) 1–25.

